# Overcoming Limitations to Deep Learning in Domesticated Animals with TrioTrain

**DOI:** 10.1101/2024.04.15.589602

**Authors:** Jenna Kalleberg, Jacob Rissman, Robert D. Schnabel

**Affiliations:** University of Missouri, Division of Animal Sciences, Columbia, MO, 65201 USA; University of Missouri, Genetics Area Program, Columbia, MO, 65201 USA

## Abstract

Variant calling across diverse species remains challenging as most bioinformatics tools default to assumptions based on human genomes. DeepVariant (DV) excels without joint genotyping while offering fewer implementation barriers. However, the growing appeal of a “universal” algorithm has magnified the unknown impacts when used with non-human genomes. Here, we use bovine genomes to assess the limits of human-genome-trained models in other species. We introduce the first multi-species DV model that achieves a lower Mendelian Inheritance Error (MIE) rate during single-sample genotyping. Our novel approach, TrioTrain, automates extending DV for species without Genome In A Bottle (GIAB) resources and uses region shuffling to mitigate barriers for SLURM-based clusters. To offset imperfect truth labels for animal genomes, we remove Mendelian discordant variants before training, where models are tuned to genotype the offspring correctly. With TrioTrain, we use cattle, yak, and bison trios to build 30 model iterations across five phases. We observe remarkable performance across phases when testing the GIAB human trios with a mean SNP F1 score >0.990. In HG002, our phase 4 bovine model identifies more variants at a lower MIE rate than DeepTrio. In bovine F1-hybrid genomes, our model substantially reduces inheritance errors with a mean MIE rate of 0.03 percent. Although constrained by imperfect labels, we find that multi-species, trio-based training produces a robust variant calling model. Our research demonstrates that exclusively training with human genomes restricts the application of deep-learning approaches for comparative genomics.

## INTRODUCTION

Machine learning applications in genomics are not new (Benzer 1959; Camin and Sokal 1965; Beyer et al. 1974). Nevertheless, the type and scale of available data continue to change what is possible through genomics-specific, deep-learning (DL) tools. Foundational, large-scale sequencing consortiums for human genomes often initiate similar work in other species (Altshuler et al. 2010; Jarvis et al. 2022; Liao et al. 2023; Kaminow et al. 2022; Nurk et al. 2022), particularly within the bovine genomics community (Hayes and Daetwyler 2019; Zhou et al. 2022a; Smith et al. 2023; Kalbeisch et al. 2024). However, aspirations of fully understanding genome biology require effectively translating between disciplines. Unfortunately, species-agnostic tool development has not kept pace with the exponential growth of sequenced biodiversity. Despite widespread variation in ploidy, chromosome number, or kinship, many bioinformatic tools expect the human genome by default. However, optimism surrounds deep-learning technologies and their potential translation into non-human species (Webb 2018).

Context matters when evaluating the performance of any statistical inference tool. When trained exclusively with human genomes, new data from other species can affect model precision (Sundaram et al. 2018). However, the burden of customizing leading-edge tools often delays their use with animal genomes. Thus, comparative genomics researchers often rely on human-based default parameters that may be inappropriate for other species (Genereux et al. 2020). Human-centric tools include the most popular variant calling software, the GATK Best Practices (McKenna et al. 2010). Species-specific alterations to the Best Practices are common but rarely standardized, meaning researchers must tediously check published workflows for outdated deviations. Pervasive inconsistencies in human genomics motivated the Genome In A Bottle (GIAB) consortium to create the first “highly accurate set of genotypes across a genome” (Zook et al. 2014). These GIAB Reference Materials (RMs) were foundational advancements that remain the authority for assessing variant discovery methods (Olson et al. 2023; Chin et al. 2020; Wagner et al. 2022).

Identifying genetic variation depends upon three components: the reference genome, the sequencing technologies, and the algorithms applied (Bickhart et al. 2020; Atkinson et al. 2023). The resources from GIAB enable objective decisions when comparing approaches (Olson et al. 2022a). For example, recent work found that variant discovery software can significantly impact genotype quality compared to the alignment method (Barbitoff et al. 2022). Benchmarking competitions have illustrated that incomplete knowledge limits heuristics-based variant discovery methods like GATK. In contrast, deep-learning-based methods can identify the unknown (Sapoval et al. 2022). The GIAB RMs were intended to help standardize communicating optimal method selection (Olson et al. 2022b). However, as the gold standard of reliable genomic variation, they have become the default source of training data for deep learning genomics models, such as DeepVariant (DV). As the first deep learning model to achieve a small yet significant improvement over GATK, DeepVariant’s innovation stems from transforming variant calling into an image classification problem (Poplin et al. 2018a). After identifying features from pileup alignments and transforming these into multi-layer tensors, the Convolution Neural Network (CNN) uses the context surrounding a putative variant when genotyping. Despite relying heavily on the human GIAB reference materials for training, subsequent research has repeatedly demonstrated DeepVariant’s superiority (Supernat et al. 2018; Zhao et al. 2020; Lin et al. 2022; Olson et al. 2022a; Barbitoff et al. 2022). Furthermore, DV avoids GATK’s scalability limitations by achieving comparable accuracy without requiring joint genotyping (Poplin et al. 2018b; Chen et al. 2023). Meanwhile, the haplotype-aware PEPPER-Margin-DeepVariant pipeline is now the standard approach for small variant detection with PacBio’s long-read sequencing (Shafin et al. 2021). Unsurprisingly, DeepVariant’s success inspired the rapid development of DL methods for variant discovery (Sahraeian et al. 2019; Luo et al. 2020; Ip et al. 2020; Ramachandran et al. 2021; Khazeeva et al. 2022; Su et al. 2022; Popic et al. 2023; Ahsan et al. 2023). Deep learning technologies saturate the leading edge of genomics, yet most overlook the challenges facing non-human genomics, limiting their relevance across biological diversity.

The revolutionizing impact of deep-learning-based approaches is attributed to publicly accessible genomic data and reliable variant callsets (Olson et al. 2023). Unfortunately, these resources only exist for human genomes, stagnating tool development for other species. Preliminary research illustrated DV’s potential with insects and plants (Yun et al. 2018; Day and Poplin 2019). However, without extensive re-training, “DeepVariant has limited applicability to non-human SNV calling” (Sapoval et al. 2022). Despite the unknown impact, the human-based default model has seen a rapid expansion in comparative genomics. DV offers substantially fewer implementation barriers than GATK, enticing researchers across a wide range of species, including domesticated and endangered animals (Leonard et al. 2023; Stenløkk et al. 2022; Flack et al. 2023), birds (Secomandi et al. 2023; Guhlin et al. 2023), plants (Osipowski et al. 2020; Ruperao et al. 2023), and pathogenic organisms (Gunderson et al. 2020; Jesudoss Chelladurai et al. 2023). As the ’default’ genome, models intended for humans are often implicitly trusted to be the optimal method in other diploid species. Here, we address the non-trivial technical barriers of customizing DV models for non-human genomes. We describe an alternative strategy for re-training for species lacking GIAB-quality truth callsets. TrioTrain enables automatic model extension with new data on a SLURM-based computing cluster. We provide our best-performing bovine-trained model to use with DeepVariant’s one-step, single-sample variant caller. Our research illustrates the limitations of using the default, human-genome-trained DeepVariant models in other species, as we find that a multi-species training approach improves performance in human and bovine genomes.

## RESULTS

### Initial variant quality indicated a training gap

DeepVariant’s specialty is providing robust models that control the known and unknown systematic bias for a wide range of sequencing technologies. The first non-human extension of DeepVariant (DV) pursued re-training with mosquitos (Yun et al. 2018). As mosquito genomes are appreciably more different from humans than cattle are, we initially expected re-training to be unnecessary. We evaluated the human-trained model in a bovine genome (UMAG Lab ID: 828) by comparing calls from 1kBulls Run8 and DeepVariant (1.0) on chromosome 29. DV identified more unique variants than the 1kBulls GATK pipeline (**Supplemental Figure 1**). However, the SNP transition to transversion (Ts/Tv) ratio was marginally lower than those unique to GAT. We then asked if DV could benefit from further training with bovine genomes.

### Model complexity cannot overcompensate for poor-quality labels

The challenge for training species-specific models is maximizing the quality of the labels used as truth. Here, we infer truth using short-read sequencing (SRS) data and GATK, which introduces a small amount of noise indistinguishable from genuine variation. For example, the initial build of TrioTrain used truth labels from the 1kBulls Run8 callset (2020), as Run 9 was in progress. Automated training enabled comparison between truth-label filtering criteria with existing data. Our preliminary analyses identified an order-of-operations flaw; loss plots indicated overfitting and potentially noisy labels (**Supplemental Note 1**). Subsequent logic testing revealed filtering inconsistencies between the truth VCF and the PopVCF. Avoiding these missteps requires appreciating how every prior step affects the data (Whalen et al. 2021). As such, we recommend TrioTrain users carefully review the limitations of their current genotyping approach before attempting training in new species.

The quality of the 1kBulls Run8 callset constrained our ability to improve performance. Despite applying more strict filters, the tuning loss hovered consistently around 0.2 instead of approaching zero. We then reviewed concordance across three recent 1kBulls callsets (2019– 2021). To our knowledge, this is the most extensive validation with these data, requiring >1.5M CPU hours (**Supplemental Note 2**). Across these genotyping runs, 3659 samples were re-processed. In theory, the genotypes for these samples should be identical except due to increasing sample size and updating parameters; our results indicate significantly lower recall and precision within the 1kBulls data than expected (**Supplemental Table 1**). These findings reveal substantial variability between genotyping runs using the same underlying raw data. Our discovery affects all downstream analyses that assume accurate and consistent genotypes.

Due to the limitations of the 1kBulls callsets, we drastically improved our GATK-based variant calling by iteratively tuning parameters during Variant Quality Score Recalibration (VQSR). Unfortunately, most users settle for performing VQSR once with default parameters based on human data. Instead, we tested a range of values for each user-tunable parameter, resulting in approximately 300–1,000 VQSR runs (Methods). We then compared our optimal parameters against the GATK defaults and the 1kBulls Run9 (2021). Our internal workflow identifies more variants within the 95 percent tranche, producing 2x fewer Mendelian Inheritance Errors (MIE) than the globally used 1kBulls Run9 callset (**Supplemental Figure 14**). While our research demonstrates significant room for improvement in the overall quality of variant call sets in animal genomics, we find that default values intended for human genomes introduce errors.

### Using TrioTrain to design bovine-specific curriculum

A lack of high-quality benchmarking variants has thwarted most pursuits for non-human DV models (**Supplemental Table 2**). However, as one of the highest quality, population-scale cohorts in animal genomics, the UMAGv1 bovine callset allows examining if existing sequence data can provide viable alternatives. TrioTrain was designed to enable training with any diploid mammalian species. We built our bovine models using trios with publicly available SRS data, targeting multiple breeds and variable coverage levels (Methods). Our work addresses several knowledge gaps. First, we asked if leveraging inheritance patterns during training and increasing the training sample size could offset imperfect truth labels. Second, we describe the genome characteristics that alter model behavior. Lastly, we identify opportunities to improve DeepVariant using the diverse ancestry with bovids while offering strategies for choosing samples.

Foundational research used trios to assess inheritance patterns and improve assembly quality (Koren et al. 2018). In the mosquito-specific DV approach, the parental genomes were never given to the model but instead used to curate inherited variants within the offspring (Yun et al. 2018). As de novo variants are rarer than systematic errors, removing MIEs maximizes label accuracy within a trio. With TrioTrain, we modified the mosquito-specific strategy based on pedigree structure, genome length, and sample size. We hypothesized that a model trained on the parental genomes would correctly genotype inherited variants in the offspring. Therefore, a family trio is divided by treating a parental genome as a training dataset, while a single offspring is used for model tuning. Each training, tuning, and testing dataset is constrained to the autosomes and X chromosome. Pedigree information is not explicitly provided to the model; instead, the optimal checkpoint achieves the maximum F1 score when genotyping the offspring. TrioTrain begins by initializing weights from the default human-trained model (**Figure 1**).

**Figure 1.**
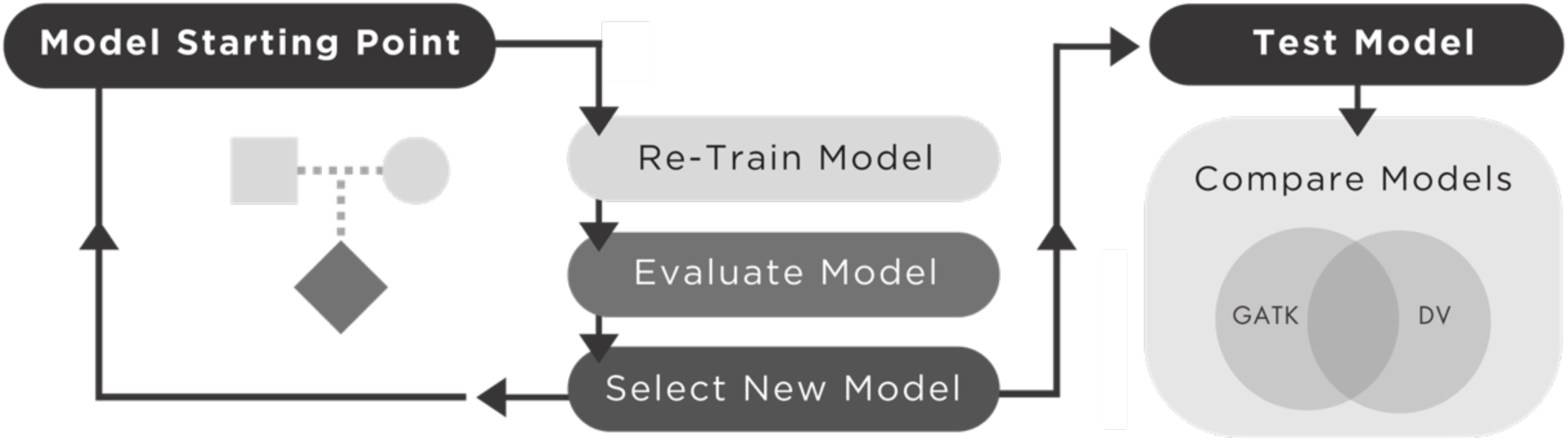
Enabling customization of DeepVariant. Workflow diagram of TrioTrain’s iterative re-training process. All re-training begins with weights from an existing model as the starting point. The first iteration uses the default checkpoint (1.4) without the allele frequency (AF) channel, while subsequent iterations include the AF channel. Two iterations are performed for each trio, one for each parent. An iteration covers all labeled examples within a parent-child duo (CHR 1-29, X). Each iteration produces a candidate checkpoint assessed using a set of test genomes previously unseen by the model. The resulting variant calls are compared against test-specific GATK-derived truth labels (UMAGv1) to observe changes over the training period.

Our sample size of trios allows for two sequential iterations of training, one for each parent genome (Methods). Within a trio, both iterations are tuned using replicate labeled examples from the child to prioritize weights that distinguish inherited variants from noise. The number of steps varies based on the examples produced for a parent, but all are covered once to limit potential overfitting. When tuning with the offspring, stratified performance metrics are calculated continuously. The checkpoint with the maximum F1 score in the offspring is selected as the warm-starting point for the next iteration. Then, the entire re-training process is repeated until all trios have been exhausted. Each training iteration is tested using an independent set of genomes unseen by DV (Methods).

We created multiple DV models to address knowledge gaps hindering training in species without GIAB-quality callsets. We built 30 model iterations, representing 15 family trios (Methods). We then subset these iterations into five phases to summarize our findings. We began by training with 9 multi-breed cattle trios to illustrate a ’cattle-specific’ model, split into Phases 1 and 2. The terminal checkpoint for Phase 1 was chosen to represent an upper-limit performance threshold when relying on typical public SRS data for training. The second phase demonstrates the effects of increasing training volume with breed-specific replicates. The remaining 6 trios are used in Phases 3 – 5. For Phase 3, we identified two bison trios with the same father from our UMAGv1 dataset. Prior research suggested “an advantage to using a diverse population” when building DV models (Chen et al. 2023). With readily available data from a related but divergent species, we compared a multi-species, ’bovine-specific’ model against the typical species-specific approach. Next, we explored alternative strategies for achieving higher-quality truth labels. The ongoing bovine pangenome project includes trio-binned, reconstructed parental assemblies.

Phase 4 uses three F1-hybrid cross trios, including both cattle subspecies, Angus-Brahman (Low et al. 2020), and two cattle-outgroup crosses, Highlander-Yak and Simmental-Bison (Rice et al. 2020; Heaton et al. 2021; Oppenheimer et al. 2021). While the Phase 4 genotypes represent our highest-confidence bovine truth variants, the offspring’s increased heterozygosity exacerbates the genotype class imbalances (**Supplemental Table 3**). Therefore, we used the alternative assemblies to build synthetic diploid (SynDip) offspring (Methods). For Phase 5, we used replicate examples from the Angus and Brahman parents but unique examples from the SynDip offspring. These previously unseen datatypes represent potential intermediate avenues for curating truth labels.

### Truth label quality restricts performance

During training, performance metrics for all checkpoints are stratified by genotype class and variant type. We compared these metrics for the selected optimal checkpoint across all 30 iterations (**Figure 2**, **Supplemental Figure 3**). The outcomes during Phases 1 – 2 suggest that training on existing callsets can achieve remarkable performance (Average F1 score ∼0.98). However, this required exhaustively improving our genotyping accuracy before training. Therefore, we caution that our findings represent an upper limit threshold rather than typical performance with non-human SRS data. Across all iterations, we find INDELs skew selected checkpoint performance, with lower precision indicating more False Positives (FPs). Since INDELs are underrepresented in our UMAGv1 truth labels, some genuine variants are misclassified as FPs during tuning (**Supplemental Table 4**). Based on this expectation, we suspect the skewed iterations (Trios 2, 5, and 9) may significantly benefit from INDEL truth labels from long-read sequencing (LRS). While most bovine training iterations perform remarkably well, without a ground truth callset, we likely overestimate performance by some small degree. However, we can use these findings to identify sample characteristics affecting learning success between iterations.

**Figure 2.**
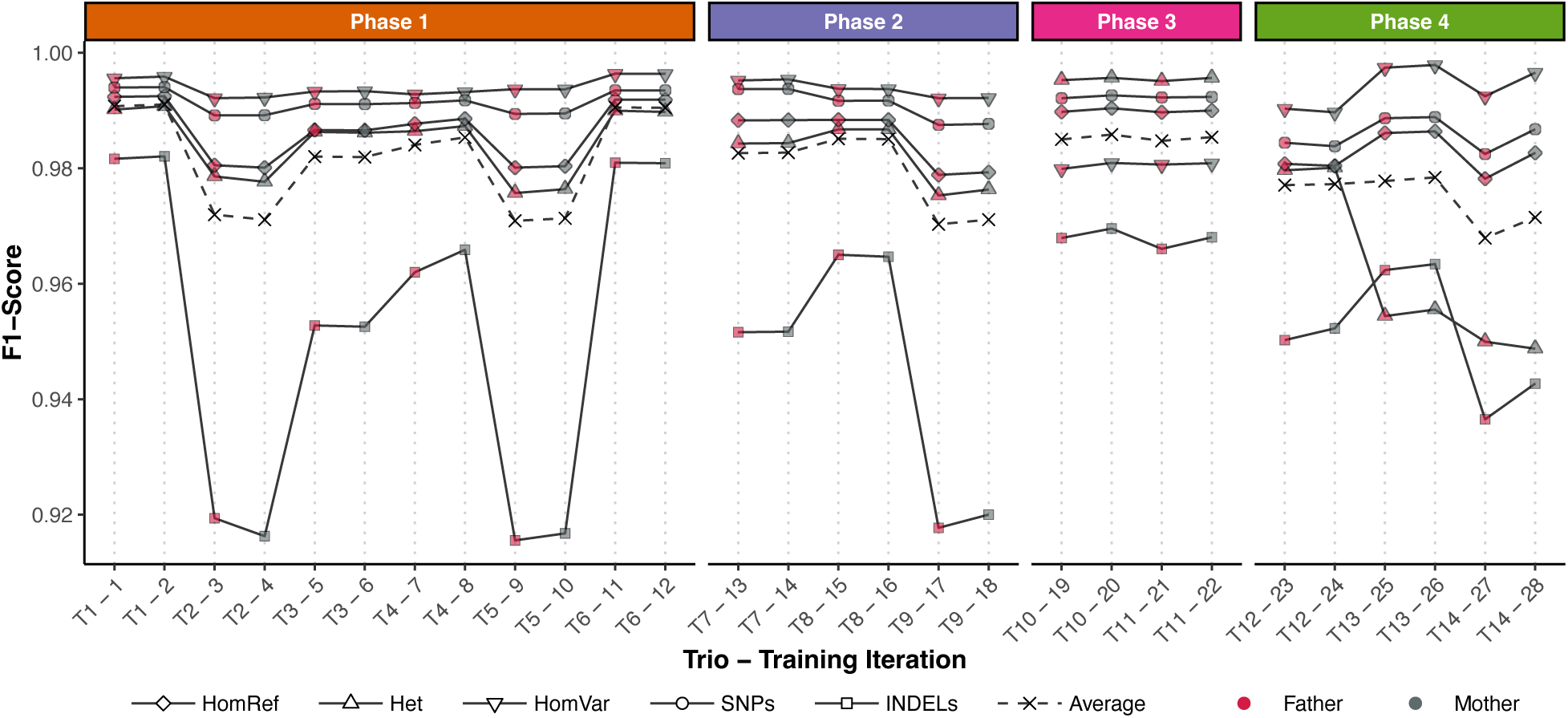
Comparing learning across phases. Training performance across multiple iterations is illustrated using F1-score, calculated as TP / TP + ½ (FP + FN). Each line represents stratified performance using variant type (INDELs, SNPs), genotype class (HomRef, Het, HomAlt), and the average across categories. The minimum y-axis value is 0.92, with perfect prediction achieved at 1. Chronological trios are marked on the x-axis from left to right, spanning 28 of 30 sequential iterations (Trio# - Iteration#). Each trio begins with labeled examples from the father, followed by the mother (C.H.R. 1 – 29, X). As expected, models underperform with INDELs due to relying on short-read WGS data and GATK for curating the UMAGv1 bovine truth labels. However, overall performance during model selection is robust, with a median F1-Score for selected checkpoints at 0.9865. TP = True Positive, FP = False Positive, FN = False Negative.

### Factors that influence training behavior

The nature of deep-learning-based methods obscures which features are relevant for accurate prediction. Previous work suggests the importance of coverage depth, demographic history, variant distribution and density, and genome complexity (Zou et al. 2019; Whalen et al. 2021; Musolf et al. 2022). Including the allele frequency (AF) channel stabilized model performance in lower coverage human genomes (Chen et al. 2023). Since agricultural genomics cohorts often prioritize a larger sample size with lower coverage, we also included the population allele frequencies from the UMAGv1 cohort. Although we see benefits of including more context, iterations where the trio’s mean coverage approaches 25x performed incrementally better (**Table 1**, **top**). However, inferring model behavior with a discrete attribute overlooks DeepVariant’s ability to learn complex and hidden interactions. Instead, our experiment design structures the training phases into biological questions as we explore the nuances of DV’s behavior with these data. For example, in Figure 2, the outlier iterations represent three Hereford trios with mean coverage <25x (**Table 1**, **middle**). These samples have fewer homozygous alternate (HomAlt) and more heterozygous (Het) genotypes due to their similarity with the Hereford reference genome (Rosen et al. 2020). The Het/HomAlt ratio indicates a source of genotype class imbalance, where balanced data approaches 1. Exclusively training with human genomes limits DV to expect distributions based on the GIABv4.2.1 truth sets, where the mean Het/HomAlt ratio is 1.46 (**Supplemental Table 3**). We give new context to DV by re-training with bovine genomes, as the class distribution deviates from prior experience. The skewed metrics observed stem from the model inflating INDEL HomAlt genotypes relative to the truth labels. Therefore, in addition to coverage bias, the outlier trios’ behavior is partially explained by the Het/HomAlt ratio approaching 2, compared to the human-genome expectation of ∼1.5. Investigating why a subset of training genomes underperform gives insight into any unintended biases in the model’s training curriculum. Our bovine model will undoubtedly be used with other Hereford samples, making it imperative for DV to experience similar data during training. As such, these iterations’ value is not in achieving perfect prediction; instead, they provide contrast emphasizing how TrioTrain users can choose training genomes strategically.

**Table 1.** Sample characteristics influence training performance. Overall, the best-performing samples have a mean sequence coverage depth near 25x, illustrating the importance of varying training coverage. <u>Top:</u> Best-performing trios (F1-score > 0.99). These include samples with more Homozygous Alternate (HomAlt) genotypes from non-reference cattle breeds compared to Heterozygous (Het) genotypes. The Het/HomAlt ratio is more balanced (⟶ 1) while being closer to the training data (GIAB mean ∼ 1.5). <u>Middle:</u> Underperforming trios during training (F1-score <0.99)). These trios are purebred or crossbred Herefords, resulting in fewer HomAlt genotypes and skewing the genotype class balance (⟶ 2). <u>Bottom:</u> Contrasting genome characteristics in divergent trios used during training. Although samples similar to the reference are essential to maintain generalizability, we recommend prioritizing sample diversity when selecting individuals to include in a training dataset.

| Trio |  |  | Checkpoint<br>F1-Score | Mean<br>Coverage | Het/HomAlt Ratio |  |  |
| --- | --- | --- | --- | --- | --- | --- | --- |
| Name | Breed | Sample |  |  | ALL | SNP | INDEL |
| High-Performing Trios |  |  |  |  |  |  |  |
| Trio1 | Angus (AA) | Father | 0.992378 | 35.96 | 1.61 | 1.61 | 1.57 |
|  |  | Mother | 0.992497 | 23.10 | 1.61 | 1.61 | 1.56 |
|  |  | Offspring | - | 23.64 | 1.50 | 1.51 | 1.46 |
| Trio6 | Tyrolean Grey (TG.) | Father | 0.991886 | 25.30 | 1.74 | 1.74 | 1.69 |
|  |  | Mother | 0.991871 | 24.10 | 1.85 | 1.85 | 1.80 |
|  |  | Offspring | - | 24.90 | 1.76 | 1.76 | 1.71 |
| Mean |  |  | 0.992158 | 26.17 | 1.68 | 1.68 | 1.63 |
| Low-Performing Trios |  |  |  |  |  |  |  |
| Trio2 | Hereford (HE) | Father | 0.980533 | 17.47 | 2.10 | 2.11 | 1.95 |
|  |  | Mother | 0.980109 | 12.92 | 2.07 | 2.08 | 1.92 |
|  |  | Offspring | - | 14.45 | 1.51 | 1.52 | 1.41 |
| Trio5 | Hereford (HE) | Father | 0.980117 | 13.51 | 1.90 | 1.91 | 1.78 |
|  |  | Mother | 0.980370 | 15.08 | 2.14 | 2.15 | 2.03 |
|  |  | Offspring | - | 15.89 | 2.00 | 2.02 | 1.86 |
| Trio9 | Holstein-Hereford (HH) | Father | 0.978856 | 17.47 | 2.10 | 2.11 | 1.95 |
|  |  | Mother | 0.979315 | 14.62 | 2.21 | 2.23 | 2.08 |
|  |  | Offspring | - | 14.79 | 2.22 | 2.24 | 2.08 |
| Mean |  |  | 0.979883 | 15.13 | 2.03 | 2.04 | 1.90 |
| Multi-Species Trios |  |  |  |  |  |  |  |
| Trio10 | Bison (BI) | Father | 0.989776 | 22.69 | 0.211 | .0211 | 0.222 |
|  |  | Mother | 0.990404 | 23.72 | 0.220 | 0.219 | 0.232 |
|  |  | Offspring | - | 17.95 | 0.216 | 0.215 | 0.227 |
| Trio12 | Angus (AA) | Father | 0.980774 | 36.99 | 1.623 | 1.627 | 1.576 |
|  | Brahman (BR) | Mother | 0.980420 | 43.32 | 1.776 | 1.769 | 1.872 |
|  | AA-BR F1 Hybrid | Offspring | - | 50.01 | 4.915 | 4.914 | 4.931 |

Continuous learning requires identifying new data sources that address training gaps. Our results indicate that training improves with breeds that differ from the reference genome. Both trios 1 and 6 have a mean Het/HomAlt ratio of 1.7, resulting in a higher tuning F1 score due to having similar levels of imbalance relative to the human training data (**Table 1**, **top**). However, deviating from prior expectations can improve performance if these data offer new context. For example, when training with bison trios, the drastic shift in the Het/HomAlt ratio unexpectedly improves prediction (**Table 1**, **bottom**). Aligning bison genomes to the cattle reference results in more HomAlt genotypes, severely altering the Het/HomAlt ratio from >1.5 to <0.5 (mean Het/HomAlt = 0.216). Although Phase 3 trains on more HomAlt examples, the precision of Het variants improves (mean F1 score = 0.99543) while the F1 score for HomAlts decreases. Counterintuitively, attempting to direct learning on specific categories by training with more of these data can have unanticipated impacts. We hypothesize that providing novel HomAlt examples from bison allows the model to resolve ambiguous signals that obscure Het variants. Our results indicate that truth variants from a related species may provide additional context for accurate variant discovery. Although model builders may intuitively gravitate towards ’breed-specific’ or ’species-specific’ explanations, we emphasize that the model remains unaware of those labels. Instead, these differences become evident to DV through differing characteristics between the priors and the new training callsets. Our findings suggest that model builders should prioritize diversity and quality when selecting their training samples.

### Improving Generalization with TrioTrain

Model generalization is contingent upon variation content and distribution differences. Identifying which models are useful requires challenging the model with previously unseen data. After completing an iteration, TrioTrain automatically performs variant calling with the optimal checkpoint using at least one independent genome. Like the trios, these samples must have a set of truth labels. When extending DV with cattle genomes, we use a multi-breed group of 15 – 19 bovine genomes from the UMAGv1 cohort that were not used previously during either training or tuning (Methods). Most of these genomes are missing either pedigree data or parental genotypes, so we are unable to exclude Mendelian discordancies from their truth labels. The UMAGv1 callset has half the number of Mendelian errors as the 1kBulls Run9 callset (Methods). As such, these data represent our closest approximation of a benchmarking callset for bovine genomes.

We followed the NIST-GIAB standards during model testing to accurately evaluate a novel model in new datasets (Methods). However, we are evaluating performance in relative terms across the training iterations. Our ability to gauge performance depends on the definition of “truth.” For our purposes, we define a False Positive (FP) as the variants discovered by DeepVariant that GATK misses. Additionally, any variant missed by DV found in the UMAGv1 truth labels is considered a False Negative (FN). Finally, True Positives (TP) are an exact match between DeepVariant and GATK. These definitions are important caveats for interpreting our findings. For example, some variants categorized as FPs could also be novel, so interpreting performance requires caution.

We find the testing F1 score rapidly increases with the first cattle-specific model compared to the human-based defaults (**Figure 3**). However, subsequent iterations during the first three phases have diminishing returns, particularly with SNPs. Although INDEL performance incrementally improves, the SNP F1 score remains consistent despite adding more trios. Ultimately, these results illustrate that if sample selection is strategic, other species with fewer trios may improve generalization in fewer iterations. For example, Phase 2 includes additional replicates of breeds used in previous iterations, three Holstein or Hereford crosses. Although dairy breeds such as Holsteins are common in public sequencing datasets, our results suggest prioritizing diverse training samples over pursuing additional iterations within the same breed. Next, in Phase 3, the two bison trios contain the same father. As such, we recommend focusing on unique trios instead of extending training with a half-sib offspring. Increasing training sample size without new context by including replicates of highly similar genomes offers minimal benefit to generalizability. Ultimately, our result demonstrates how sample quality rather than quantity can be more effective.

**Figure 3.**
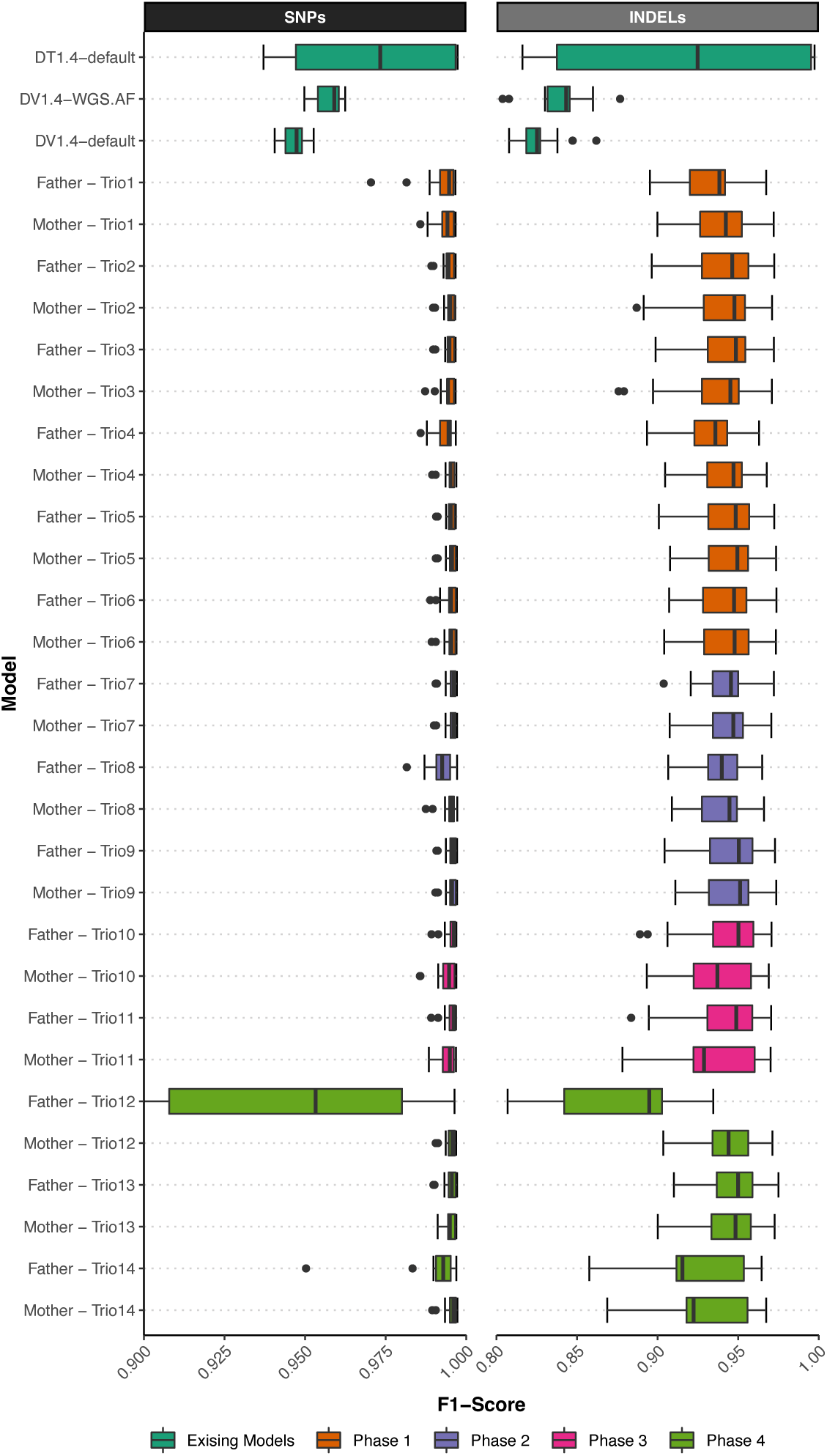
Comparing variants from bovine genomes. (Next Page) Each box-and-whisker represents a model’s F1-score in a set of bovine samples previously unseen by the model. Sample size differs by model, as N=6 with DeepTrio, while N=19 with Phases 1 – 3, and starting in Phase 4, N decreases by one with each additional trio until N=15. For each test genome, F1-score is calculated by comparing DV variant calls against the UMAG v1 truth labels. Models are presented chronologically from top to bottom on the y-axis: two alternative human-trained models, the re-training starting point, followed by 28 of 30 training iterations. The x-axis scale begins at 0.9 and 0.8 for SNPs and INDELs, respectively, where 1 would indicate perfect prediction. As with training, SNPs outperform INDELs. In these samples, we find the human-trained models underperforms relative to all bovine-trained models. Note that iteration 23 (Father-Trio12) is the first iteration with a hybrid-cross offspring. This unique case extends the whisker SNPs out of view (minimum F1-Score=0.826). However, the next iteration allows the model to recover, indicating the benefits of our trio-based training approach.

Performance gains continue with INDELS because fewer are provided per iteration relative to SNPs. The UMAGv1 truth labels are overwhelmingly from SNPs (mean SNP:INDELs ratio = 8.39). Therefore, more iterations are required to accumulate the same volume of examples. These findings illustrate that if the data are available, multiple iterations of re-training provide value by covering gaps in training data, specifically any underrepresented prediction classes, such as INDELs or HomAlt variants. For example, performance deviated substantially during the first iterations of Phases 4 – 5 (**Figure 3; Supplemental Figure 4**). While such inconsistent F1 scores imply an inappropriate iteration, performance recovers after a second round of training with the second parent. The wide range of F1 scores is expected, given the drastic shift in the Het:HomAlt ratio between the F1-hybrid offspring and the test genomes. We recommend using repeated iterations with novel data, such as our trio-based approach, to allow the model to adjust. However, a lack of precision across new test sets can distinguish inappropriate models.

For example, when given bovine genomes, all human-trained models behaved unreliably, drastically reducing performance, particularly with Heterozygous and INDEL variants. The higher proportion of INDEL FPs primarily drive the performance decrease. In a well-tuned mammalian model, we expect very few genuine variants DV detects to be incorrectly labeled as FPs, even in new species. Instead, we find that giving cattle samples to the human-trained models considerably increases FPs. Across all three existing models, we observe a mean FP rate of 15.86 percent (SNP mean = 7.60%, INDEL mean = 24.12%) with our bovine test set. However, after one round of re-training, the FP rate drops to 2.12 percent (SNP mean = 0.59%, INDEL mean = 3.64%).

Additionally, re-training with the UMAGv1 truth labels improves concordance between models. With our bovine test set, the existing DV models have an average TP rate of 78.65 percent (SNP mean = 87.76%, INDEL mean = 69.54%). In contrast, the final iteration of TrioTrain’s Phase 4 identifies over 12 percent more, with an average TP rate of 91.50 percent (SNP mean = 96.83%, INDEL mean = 86.18%). A more substantial proportion of TPs and fewer FPs demonstrates improvement relative to the default model built exclusively with human genomes. However, we can only use these findings to contrast changes in the model relative to our GATK-based callsets. With a GIAB-quality truth set for bovine genomes, we could authoritatively select the optimal model for cattle genomics. Overall, we find that an optimal model for human genomes may not be appropriate in all mammals.

### Benchmarking with Human Genomes

To summarize our results, we selected a subset of models, specifically the terminal checkpoints from each phase. These five checkpoints were used to call variants with the human GIAB samples. Callsets from all five models are compared against the existing DV models, including the default WGS, adding allele frequencies (AF) of confident variants, and DeepTrio (v1.4). We use previously described methods to perform benchmarking with the GIAB genomes (Krusche et al. 2019). As expected, the cattle-trained model struggles with INDEL recall in human genomes (**Figure 4**, **Supplemental Figure 5**). Our bovine truth labels likely exclude potentially real INDELs since the curation process relies on GATK with SRS. Known issues include the shorter read length, which introduces mapping errors in highly repetitive regions, and INDEL accuracy, which varies with coverage. Ultimately, our truth set misses some long and more complex INDELS. Despite the imperfect truth set, our Phase 4 bovine model performs remarkably well with the human benchmarking trios. SNP performance between the two allele frequency models is remarkably similar (SNP F1 score: human. AF = 0.99559, bovine.Phase4.AF = 0.990347, Δ = -0.00524). After training with our F1-hybrid truth sets, the bovine AF model produces 2,666 more SNPs in HG002 than the comparable human model (**Table 2**). These findings illustrate that the SNP calls in our GATK-derived truth set from our optimized UMAGv1 dataset are robust

**Figure 4.**
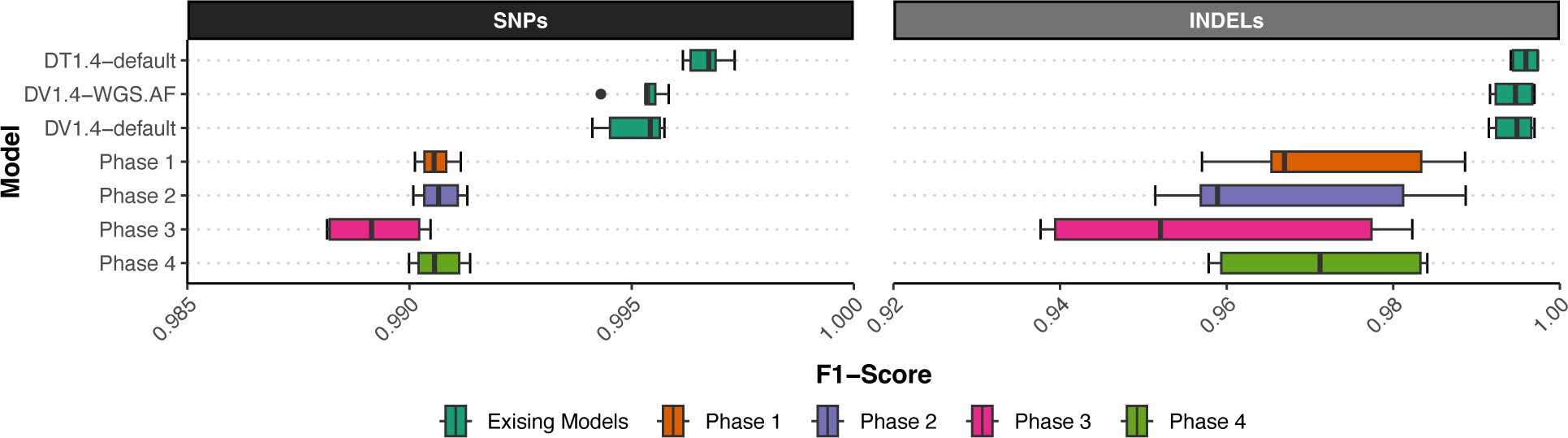
Benchmarking with human GIAB trios. (Next Page) Each box-and-whisker represents a model’s F1-score with the GIAB human trios (N=6). We compared the variants produced by each model against their respective GIAB v4.2.1 benchmark sets. Models are presented chronologically from top to bottom on the y-axis, starting with two alternative human-trained models, then the re-training starting point, followed by the terminal checkpoint from 4 out of 5 TrioTrain Phases. Note that the x-axis scale begins at 0.985 and 0.94 for SNPs and INDELs, respectively, where 1 would indicate perfect prediction. We find INDEL accuracy in the training labels hinders subsequent performance from TrioTrain models. Despite this limitation, we observe remarkable performance from our bovine-trained models with the GIAB human genomes (mean SNP F1-Score > 0.990) apart from the bison-trained model from Phase 3. See supplemental figures for Phase 5 results.

**Table 2.** Model Summary. The table below summarizes the proportional error rate (top) and variant counts (bottom) across the three existing DeepVariant and DeepTrio models. We contrast default models relative to the Phase 4 bovine model and the UMAGv1 callset before removing Mendelian Inheritance Errors (MIEs). Compared to defaults, our multi-species bovine model identifies more variants while achieves a MIE rate in the human GIAB trios. Furthermore, when using the same bovine-trained model in the F1-hybrid trios, we observe a drastic reduction in MIE rate, particularly compared to the unfiltered UMAGv1 callset.

|  |  |  | Human |  | Bovine |  |  |
| --- | --- | --- | --- | --- | --- | --- | --- |
|  |  |  | Ashkenazi Jewish (AJ.) | Han Chinese (HC.) | Angus/ Brahman F1 (AA/BR) | Yak/ Highlander F1 (YK/HI) | Bison/ Simmental F1 (BI/SI) |
|  |  |  | HG002 | HG005 | Trio12 | Trio13 | Trio14 |
| Human | Model | Version | Mendelian Inheritance Error Rate |  |  |  |  |
|  | DeepTrio | default | 0.24% | 0.36% | 0.13% | 0.09% | 0.11% |
|  | DeepVariant | default | 0.45% | 0.53% | 0.23% | 0.17% | 0.19% |
|  | DeepVariant | WGS.AF | 0.39% | 0.48% | 0.17% | 0.12% | 0.11% |
| Bovine | DeepVariant | WGS.AF | 0.21% | 0.53% | 0.03% | 0.03% | 0.03% |
|  | GATK | UMAGv1 | - | - | 3.15% | 2.67% | 5.40% |
| Human | Total Number of Variants |  |  |  |  |  |  |
|  | DeepTrio | default | 4,635,843 | 4,715,887 | 14,375,708 | 28,863,353 | 28,914,856 |
|  | DeepVariant | default | 4,048,333 | 4,647,507 | 14,309,064 | 28,528,607 | 27,985,861 |
|  | DeepVariant | WGS.AF | 4,645,538 | 4,635,384 | 13,502,611 | 26,851,253 | 26,013,975 |
| Bovine | DeepVariant | WGS.AF | 4,700,929 | 4,733,705 | 13,095,833 | 25,419,283 | 24,822,205 |
|  | GATK | UMAGv1 | - | - | 20,933,910 | 32,284,325 | 32,966,132 |

### Fewer Mendelian Inheritance Errors without joint genotyping

As the F1 score depends on the quality of the imperfect bovine truth labels, we used an additional metric, the Mendelian Inheritance Error (MIE) rate, to assess model performance across species. Although TrioTrain models are not directly given pedigree information, our training approach was designed to reduce the number of Mendelian discordant genotypes introduced during variant calling. We infer that a lower MIE rate indicates a better DV model. For these analyses, we used the GIAB Ashkenazi Jewish (AJ) and Han Chinese (HC) trios, along with three bovine F1-hybrids (Methods). We first assessed using bovine-trained models in bovine genomes. All DeepVariant models have a lower MIE rate than the GATK-based UMAGv1 callsets with only PASS variants (**Table 2**). Across the seven DeepVariant models compared, the terminal checkpoint from Phase 4 resulted in proportionally fewer Mendelian discordant variants in all three F1-hybrid bovine trios (**Figure 5**).

**Figure 5.**
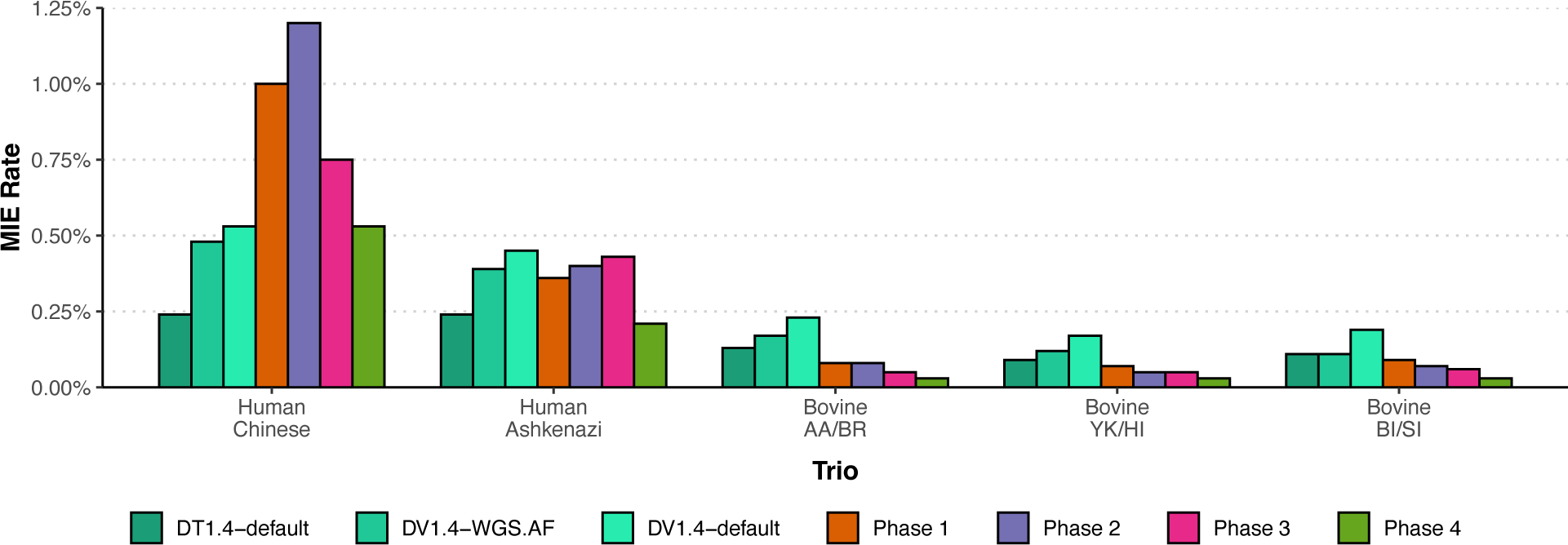
Inheritance error rate in human and bovine trios. Mendelian Inheritance Errors (MIE) were identified in PASS variants in the autosomes and X chromosome for two GIAB human trios and six bovine hybrid-cross trios. The above bar plot contains the number of discordant variants proportional to the total number of variants analyzed (PASS). The x-axis groups different trios, with each bar representing performance from a unique model checkpoint. One of the best-performing models for the GIAB samples is DeepTrio (DT). While DT is a joint caller requiring a minimum of two samples with known pedigree, our TrioTrain-built model reduces the MIE rate in individual samples, enabling drastic speed improvements. The bovine-trained Phase 4 model can produce fewer Mendelian discordances in human and bovine samples. With the Ashkenazi (AJ) trio, our best bovine-trained model produces more variants yet fewer discordant calls than any human-exclusive model, including DT. The Han Chinese (HC) trio deviates from all other trios, with the MIE rate nearly doubling during the first two phases. However, our Phase 4 model achieves an identical MIE rate as the default human model (0.53%). See supplemental figures for Phase 5 results.

Remarkably, the bovine model from Phase 4 decreased the Mendelian discordancy rate slightly compared to the DeepTrio (1.4) model, which uses separate parental and offspring models to identify variants in related duo or trio samples (Kolesnikov et al. 2021). As a joint caller, DeepTrio typically outperforms DeepVariant; however, the wall time to produce results varies significantly, depending on sample coverage and variant content. DeepVariant produces results in as little as six hours when embarrassingly parallelized. Meanwhile, DeepTrio takes at least two days to achieve the same results. In contrast, TrioTrain models reduce the MIE rate for individual samples and do not require joint calling. Thus, the resulting models built with TrioTrain enable the discovery of inherited variants in less time.

For our last bovine training phase, we find that the SynDip-trained model increases the MIE rate in all trios compared to the phase 4 checkpoint (Supplemental Figure 6), illustrating the consequences of altering the model’s expectations of noise by training with synthetic data. As such, we do not consider synthetic data derived from these assemblies a viable source of training data based on the current Quality Values (QV). However, assembly quality is rapidly improving, and thus, further research is needed to re-evaluate our findings as Q100 assemblies are pursued. We recommend estimating the per-base error rate before creating synthetic training data from any assembly (**Supplemental Table 10**).

Next, we compared using the bovine-trained model in human genomes. Our bovine Phase 4 model achieved the lowest Mendelian inheritance error rate in the AJ offspring (HG002) (**Table 2**). Notably, for the AJ family, the bovine-specific TrioTrain model discovered over 65k more variants with a lower rate of Mendelian discordancy than DeepTrio. As the HG002 genome is more commonly used as a general benchmarking sample, we further characterized the two alternative allele frequency models in the AJ trio (**Table 3**). This comparison illustrates the tradeoff of using our bovine-trained model in human genomes: under-calling INDELs. The bovine Phase 4 model calls approximately 31k fewer INDELs than the comparable human model. The observed limitation with INDELs is expected given the bovine truth label curation process; future improvement may be possible with LRS-based truth labels. Notably, performance with the HC offspring (HG005) deviates by performing worse during re-training with cattle – the MIE rate nearly doubles with the first two phases of TrioTrain (**Figure 5**). However, performance in HG005 recovers over the entire training period as the phase 4 bovine model achieves the same Mendelian discordancy rate as the default human model while producing 86k more variants (**Table 2**). While the existing human-trained models identify more variants in bovine samples, they also introduce more variants that violate Mendelian inheritance expectations (**Figure 5**). Although pedigree is not directly given during training, our model training approach implemented with TrioTrain does reduce the number of Mendelian discordant variants in our bovine samples and HG002. Ultimately, we selected the phase 4 checkpoint as our best-performing bovine model.

**Table 3.**
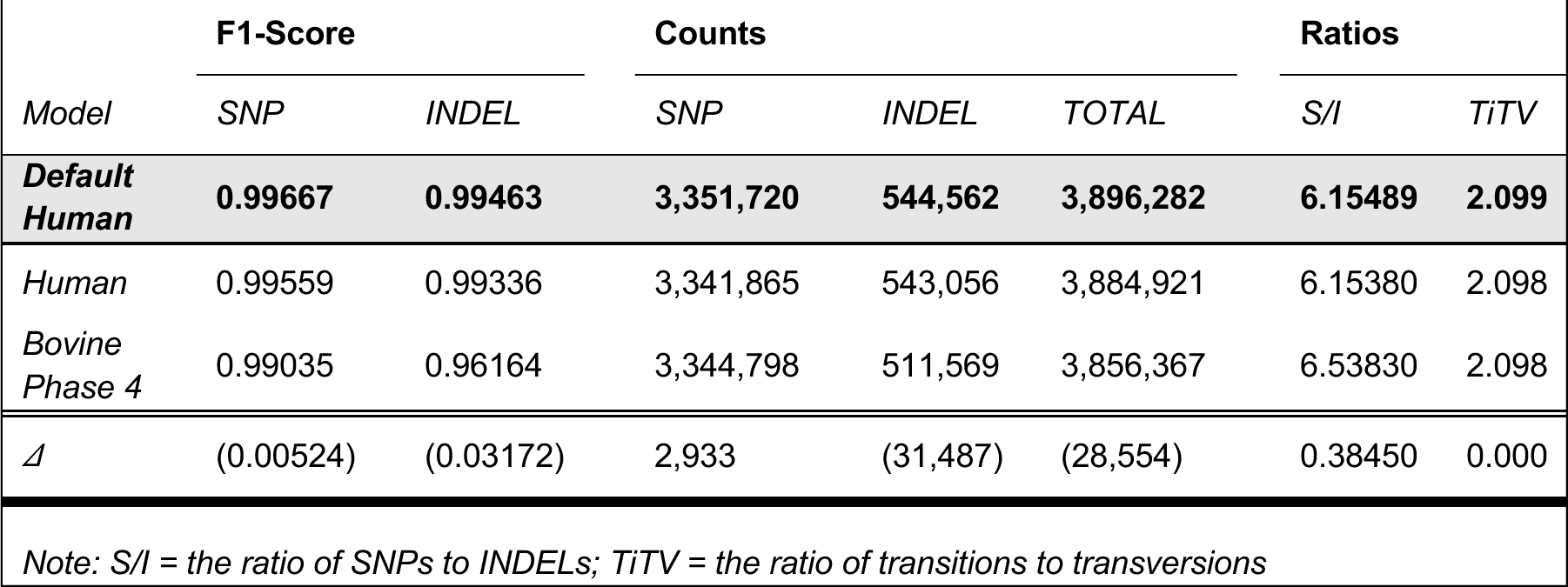
Model comparison using HG002. The table below highlights the differences between the two species-specific allele frequency models against the default DeepVariant model (bold). The analysis is constrained to GIAB-truth regions (4.2.1) for the Ashkenazi Son sample (HG002). The bottom row highlights how performance changes after re-training with bovine genomes. Parentheses indicate that a metric has decreased in the bovine-specific allele frequency model relative to the human-specific alternative. The SNP F1-Score and the TiTv ratio for our bovine Phase 4 model are comparable to the default model, with nearly 3,000 additional SNP variants in benchmarking regions relative to the DeepVariant W.G.S. AF human model. The decrease in INDELs is expected due to limitations with our cattle genotype truth labels.

## DISCUSSION

The remaining challenges within genome biology require technologies designed for the continuously expanding diversity of sequencing data. Variant discovery is rarely the sole objective but the first of many hurdles encountered during comparative genomics research. Currently, analyzing non-human genomes requires compromising between time and quality; the difficulty of optimizing species-specific methods leads many to rely on the defaults intended for the human genome. Studying critically endangered species adds urgency to the decision, as recording population diversity is essential to monitor decline and prioritize intervention. However, the growing prevalence of deep-learning-based methods introduces unknown effects of implicitly trusting a sophisticated model. For example, recent work built a customized model for the endangered Kākāpō parrots (Guhlin et al. 2023). Prior to re-training, the small population size (<200) impacted GATK’s accuracy, resulting in a higher rate of Mendelian inheritance errors compared to DeepVariant (v0.9) (3.38 and 1.28%, respectively). However, using DV to create the Kākāpō truth labels leads to circular training that may unintentionally retain bias towards the human genome. Additionally, the SNP Mendelian inheritance error rate reported for Kākāpō is 2.6x higher than our benchmarks with the default DeepVariant (v1.4) with the GIAB samples (mean: 0.49%). The contrast illustrates that researchers cannot assume the human-genome-trained DV will perform similarly in other species. Despite potential bias from the human genome, the Kākāpō callset was curated by altering Mendelian discordant sites in the offspring to match the parental genotypes. There are always tradeoffs during experiment design, particularly in novel or endangered species, but our research with Trio quantifies their magnitude.

Regardless of method, relying on defaults intended for human genomes has consequences. As with Kākāpō, parallel research using the default human-genome-trained DeepVariant (v1.5) to polish cattle assemblies reported double the SNP error rate (1.06%) relative to our benchmarks with humans (Leonard et al. 2024). For context, prior to removal of Mendelian discordant sites, the initial VQSR-optimized GATK callsets for Phases 1 – 3 trios have a minimum MIE rate of 1.14% (mean = 2.40%, max = 3.92%). The overlap between GATK and DV illustrates the nuances of method selection in comparative genomics. A lack of species-specific benchmark callsets obstructs interpretation and communication. Here, we approximate truth using our VQSR-optimized UMAGv1 callset as an independent callset re-training. Using our bovine-trained model in the GIAB samples demonstrates that our imperfect truth labels provide new context rather than echoing prior bias. For example, we observe a lower MIE rate in HG002 with our Phase 4 model than even DeepTrio (v1.4). Additionally, the bovine-trained model achieves a mean F1 score for SNPs of 0.9906 during benchmarking with the GIAB samples. Lastly, our bovine-trained model reduces the error rate in bovine trios by an order of magnitude compared to the default DeepVariant model. Our findings suggest that implicitly trusting the genotypes from DV in another species exacerbates potential error accumulation. As such, we caution against inferring that performance will replicate consistently in another species. However, exploring the behavior of such models can help identify the hidden unknowns in genomics.

We built TrioTrain to enable others to reproduce our approach for any species with the necessary input data; however, our work revealed the potential of building a ‘golden model’ for comparative genomics. Previous work focused on building species-specific models that require continuous re-training every time the human model is improved. However, developing separate models for each species is cost-prohibitive across the exponentially growing list of species with sequencing data. Instead, we find that training with a bison trio (Phase 3) improves performance in cattle genomes, suggesting the benefits of training within genera. Further research is required to investigate if the human-trained model would benefit from re-training with other hominid species. Ideally, DeepVariant would become generalizable across more broad phylogenic classifications. Our observation of a lower MIE rate in HG002 after re-training in cattle suggests the potential for a ‘golden model’ in mammals. In building TrioTrain, we provide a tool for exploring training across species and the impacts on learning, potentially revealing which phylogenic boundaries achieve optimal performance. Our work highlights the potential for using the behavior of deep-learning models to investigate the persistent challenges in genomics.

For other researchers to have maximum confidence in our new model, we require a GIAB-quality truth set for cattle. However, the UMAGv1 cohort call set vastly exceeds typical variant quality in animal genomics, even compared to the community standard, Run9, from the 1kBulls project. Short of producing species-specific GIAB benchmarks, these data serve as the upper limit of variant quality achievable for any agricultural species. While referred to as the “truth labels,” genotypes used for training in non-human species must be considered preliminary due to a lack of GIAB benchmarking variants. They cannot represent all possible true variants; instead, they are regions we can reliably and accurately genotype. Reliance on SRS data for truth labels inherently excludes highly repetitive and GC-rich regions due to shorter read length and platform-specific limitations. Ongoing work to build pangenomes with telomere-to-telomere quality assemblies will drastically impact bovine variant quality. For example, incorporating long-read technologies will address the current bovine model’s limitations with INDELs. Our future research will use these developing resources to build GIAB-quality truth sets for agriculture, starting with bovines.

Despite known limitations with the labels given to the model for training, we demonstrate achieving improvement to the DeepVariant model by including more diverse training examples. We provide a novel training design addressing the lack of GIAB samples and explore the benefits of training with multiple closely related species. For the first time, we explore training DeepVariant in other species while including the allele frequency channel. During this research, an updated model checkpoint included channels for insert size and the human 1000Genomes allele frequencies. Future research is required to evaluate if significant performance changes are achieved by starting with this combined model checkpoint. Since our model works on individual samples, we achieve comparable performance to trio-based genotyping in less time. Our research demonstrates that a comparative training protocol improves model generalizability. We hypothesize that TrioTrain enables DeepVariant to select model weights to identify inherited variants. Additionally, we show opportunities to create added value for existing data in non-human genomics research. Innovating with deep-learning technologies requires looking beyond human genomes by creating diverse, pedigree-based community resources for other species. With TrioTrain, we provide the first DeepVariant model trained in multiple mammalian species for comparative genomics research.

## METHODS

### Raw sequencing data

The University of Missouri Animal Genomics (UMAG) group maintains a repository containing the publicly available genomes for several agricultural species, including bovines. As new samples are published, the paired-end, short-read Illumina WGS data is obtained from the NCBI’s Sequence Read Archive (SRA). We identify bovine samples based on SRA metadata for taxon ID for multiple species (**Supplemental Note 3**). Where available, we also collect sex, breed, and pedigree information. Samples are aggregated based on BioSample ID and assigned an internal, integer-based UMAG lab ID unique to a single animal. We refer to the bovine cohort throughout as the UMAGv1 cohort (**Supplemental Note 4**). At the time of analysis, the cohort included 5,612 individuals with complete metadata, spanning modern and ancient genomes from related bovine species: Bos taurus taurus (Taurus), Bos taurus indicus (Indicine cattle), Bison bison (American Bison), Bos mutus (Yak), Bos grunniens (Yak), Bos frontalis (Gayal), Bos javanicus (Banteng), and Bos gaurus (Gaur).

### Whole genome sequence alignment

After downloading the raw FASTQ files from SRA, the data are pre-processed (**Supplemental Note 5 – 8**). Briefly, low-quality bases and adapters were trimmed from the reads with Trimmomatic (0.38/0.39) (Bolger et al. 2014) and subsequently aligned to the reference genome, ARS-UCD1.2 with Btau5.0.1 Y chromosome (Rosen et al. 2020) using either BWA-MEM (0.7.17) (Li 2013) or BWA-MEM2 (2.2.1) (Md et al. 2019). The resulting sequence alignment map (SAM) output is sorted, indexed, and compressed into binary alignment map (BAM) files. Duplicate reads are marked with Picard MarkDuplicates (2.18.19 or 2.26.10) (Broad Institute 2019). Our internal pipeline automatically performs realignment and base quality score recalibration (BQSR), producing callable region BED files (**Supplemental Note 9 – 10**). Note that DeepVariant does not require the realignment and recalibration of raw data. Coverage and other summary metrics were calculated using Samtools (1.9) (Danecek et al. 2021) (**Supplemental Note 11**). The average coverage for the entire cohort was 12.77x (minimum = 0.07x, maximum = 182.78x).

### Initial SNV and INDEL variant calling

The processed sequence data was used for germline variant calling for single nucleotide variants (SNVs) and short insert and deletions (INDELs). We refer to the resulting VCF as the UMAGv1 callset. Genotypes for the UMAGv1 cohort were prepared for all 6,893 bovine samples, including outgroup species, using an internal workflow that expands upon the established 1kBulls GATK FASTQ to GVCF guidelines (Hayes and Daetwyler 2019). The 1kBulls protocol uses GATK HaplotypeCaller (4.1) (Poplin et al. 2018b) to generate single sample callsets, which are merged before joint genotyping (**Supplemental Note 12 – 13**). However, using bovine trios, our internal workflow is further optimized during VQSR (**Supplemental Note 14 – 17**). In brief, significant improvement was made during Variant Quality Score Recalibration (VQSR) as parameters were extensively tuned from defaults intended for human genomes.

### Sample Selection

A subset of individuals from the final UMAGv1 callset were selected as the high-quality bovine genotypes used for re-training DeepVariant. We used mean sequencing coverage to select candidate samples from the UMAGv1 cohort with moderate sequencing depth by excluding samples with a mean coverage of <10x. Related samples were identified using pedigree metadata. Candidates for training samples were constrained to parent-offspring trios by excluding samples missing pedigree data or samples with sequencing data for only one parent. We identified 14 unique trios, resulting in 40 unique variant datasets with a mean coverage of 25.99x (minimum = 10.66x, maximum =50.83x). These 14 trios consist of 12 sires with 14 offspring and dams across multiple cattle breeds and crossbreeds (**Supplemental Note 18, Supplemental Table 6**). In addition to the trios, we identified other genomes we could use to represent the final data given to the deep-learning model. These genomes were used to monitor performance changes over the iterative re-training process and assess the generalizability of each model created with TrioTrain. We selected 13 samples from multiple cattle breeds with a mean coverage of 29.63x (minimum = 13.54x; maximum = 60.00x). These testing genomes included representatives from 15 cattle breeds, with eight additional breeds not used during training (**Supplemental Note 19, Supplemental Table 8**).

### Truth Set Curation

From the UMAGv1 cohort dataset, we extracted variants for all three samples within a trio into family VCF files (**Supplemental Note 20**). Briefly, within each family VCF, we removed any Mendelian discordant genotypes. For example, at one locus, if one parent is A/B and the other parent is B/B, but the child has a genotype A/A, that position was dropped from all three individuals. We then split the family VCF into single-sample VCFs. Due to a lack of parental sequencing data, Mendelian discordant variants could not be removed for the non-trio samples used during testing. The single-sample VCFs used for training, tuning, and testing were then filtered to retain PASS filter sites only, where PASS was set to tranche level 99.7. Lastly, to meet the specifications required by DeepVariant for labels, the final VCFs exclude all homozygous reference genotypes.

### Synthetic Diploid Offspring

Synthetic diploid (SynDip) approaches have been proposed as alternatives to GIAB-quality benchmarking data (Li et al. 2018). Ongoing efforts to produce multiple high-quality bovine assemblies could be used to generate high-quality synthetic benchmark variants for cattle (Smith et al. 2023; Zhou et al. 2022b). To compare both strategies of truth label curation in non-human species, we used the haploid parental assemblies to create SynDip offspring for all three trios (**Supplemental Note 21**). Briefly, we created synthetic reads from each parental assembly using NEAT (3.2) (Stephens et al. 2016), introducing no sequencing error, then merged these synthetic reads, resulting in three more offspring replicates. Each synthetic trio consists of the parents’ original short-read WGS data, but the offspring is substituted with the pseudo-diploid synthetic SRS. The SynDip F1-hybrid offspring have unique identifiers, separate truth labels, and a different noise signature relative to the non-synthetic samples. The three unique F1-hybrid offspring and their respective SynDip replicates (n = 6) have a mean coverage of 26.7x (minimum coverage = 14.46x; maximum coverage = 50.01x).

### Shuffling Labeled Examples

First, TrioTrain v0.8 calculates the number of regions based on the variants within the Truth VCF input for a genome (**Supplemental Note 22 – 23**). These regions’ BED files are passed to DeepVariant through the `--regions` flag. We designed region shuffling to batch the complete genome to create representative samples of the total examples. These region-specific steps are submitted across compute nodes as independent SLURM jobs. Crucially, a region-specific tfrecord file contains examples produced from across the genome. Next, within-region shuffling is performed, and then all tfrecord “shards” are concatenated into a single file. Finally, the resulting labeled and shuffled examples across all regions are fed to the DeepVariant model in a randomized order. Our Regions Shuffling process successfully enables the DeepVariant Beam shuffling pipeline on a SLURM-based HPC cluster. Rather than rely on a different resource manager, our pipeline creates embarrassingly parallel shuffling jobs. Furthermore, our design avoids extensive system-administrator support to create a Spark cluster on top of the existing SLURM-controlled cluster. Instead, our approach uses Beam’s Direct Runner to shuffle examples. Direct Runner is constrained to the local memory within a single compute node and is typically applied for pipeline debugging. However, SLURM effectively controls multiple independent Direct Runner jobs simultaneously.

### Initializing model weights

Due to the wide variability in coverage levels within the complete UMAGv1 cohort, we opted to add the allele frequency channel during our re-training. Before re-training beings, an existing model checkpoint from human genomes is used to initialize model weights, otherwise known as “warm starting.” At the time of analysis, the only WGS-specific model available with DeepVariant (1.4) added a new channel for insert size compared to previous model versions. However, this default model did not include an alternative channel encoding population-level allele frequency data from the 1000 Genomes project that had previously been available (1.1) (Chen et al. 2023). Therefore, we used the default checkpoint as the training starting point (**Supplemental Note 24**). Future research is required to evaluate warm starting with an updated human AF model. We created the PopVCF using the UMAGv1 callset by dropping genotypes and retaining only variant allele frequencies for the joint genotypes across nearly the entire cohort (n = 5,512).

### Re-Training

Running TrioTrain requires organizing metadata and file paths into a single metadata CSV file (**Supplemental Note 21**). The number of examples created and used varies per trio (**Supplemental Table 7**). For re-training the network, we used a learning rate of 0.005, a batch size of 32, and defined the training period as one epoch that spans all labeled candidate variants, known as examples, for a training genome. Performance metrics in the offspring from the selected best checkpoint for all 30 iterations were stratified by variant type and genotype (**Supplemental Table 4**).

### Summarizing Performance

We obtained the 1000Genomes population VCF from Google Genomics and the GRCH38 reference genome. We then acquired the GIAB benchmarking files (4.2.1), corresponding sample-specific callsets, and region files from the National Institute for Standards and Technology (NIST) FTP site (**Supplemental Note 25**). We used two trios: (1) Ashkenazi trio with HG002_NA24385_son, HG003_NA24149_father, HG004_NA24143_mother; (2) Han Chinese trio with HG005_NA24631_son, HG006_NA24694_father. Benchmarking parameters were identical to those used to calculate bovine performance (**Supplemental Note 26**). The only difference is switching to the human reference genome, truth regions, and the corresponding model checkpoints. The resulting VCFs were then passed to hap.py (0.3.12) using the appropriate reference genome and truth file. Performance metrics were then calculated using the same Python script for bovine test genomes, with the F1 score plotted with R.

### Validating Performance Using Mendelian Discordancy

Without a benchmarking callset for bovine genomes, we cannot directly compare the F1 score across species. However, “most methods used to identify and genotype genetic variants from next-generation sequencing data ignore the relationships between samples, resulting in significant Mendelian errors, false positives, and negatives” (Cleary et al. 2014). Therefore, we used the Mendelian Inheritance Error (MIE) rate to infer relative performance changes between species. We used Real Time Genomics (RTG) software, rtg-tools mendelian (3.12.1), to calculate the MIE rate (Cleary et al. 2015). For this analysis, we again used the same models from GIAB benchmarking: five terminal checkpoints from the TrioTrain phases and the three existing DeepVariant v1.4.0 models (default WGS, WGS.AF, DeepTrio). However, we used human and bovine trios, specifically, the two GIAB trios and the three F1-hybrid trios described previously. Each sample was re-run using DeepVariant separately, except with DeepTrio, a joint caller; all models created per-sample VCFs as output (**Supplemental Note 27**).

### Data Access

To ensure reproducibility, the exact steps used to obtain raw data listed below can be found with bash scripts on GitHub: https://github.com/jkalleberg/DV-TrioTrain/tree/main/scripts/setup. The complete documentation for our TrioTrain pipeline, including install instructions, is available on GitHub: https://github.com/jkalleberg/DV-TrioTrain.

## COMPETING INTEREST STATEMENT

The authors declare that they have no competing interests.

## ACKNOWLEDGMENTS

The research reported here was partly supported by NIH NLM with award number T32LM012410 and USDA NIFA with award numbers 2020-67015-31675, 2023-67015-39261, and HATCH MO HAAS0001. We thank the MU Animal Genomics group for their feedback and discussions on this work. The computation for this work was performed on the high-performance computing infrastructure operated by Research Support Solutions in the Division of IT at the University of Missouri, Columbia, MO DOI: https://doi.org/10.32469/10355/97710

## AUTHOR CONTRIBUTIONS

RDS and JK conceived the study. RDS performed the initial variant calling with GATK, extensively optimized the VQSR parameters to produce the final UMAGv1 callset, and curated the truth labels. JR developed code to identify callable regions. JK built the TrioTrain pipeline with input from RDS. JK performed all re-training iterations with bovine genomes and performed the subsequent analyses for model benchmarking, comparison, and evaluation. JK and RDS wrote the manuscript. All authors have read and approved the final manuscript.

## Notes

### Competing Interest Statement

The authors have declared no competing interest.

## REFERENCES

1. Ahsan MU, Liu Q, Perdomo JE, Fang L, Wang K. 2023. A survey of algorithms for the detection of genomic structural variants from long-read sequencing data. Nat Methods 20: 1143–1158.

2. Altschul SF, Gish W, Miller W, Myers EW, Lipman DJ. 1990. Basic local alignment search tool. J Mol Biol 215: 403–410. https://linkinghub.elsevier.com/retrieve/pii/S0022283605803602.

3. Altshuler DL, Durbin RM, Abecasis GR, Bentley DR, Chakravarti A, Clark AG, Collins FS, De La Vega FM, Donnelly P, Egholm M, et al. 2010. A map of human genome variation from population-scale sequencing. Nature 467: 1061–1073. 10.1038/nature09534.

4. Atkinson EG, Artomov M, Loboda AA, Rehm HL, MacArthur DG, Karczewski KJ, Neale BM, Daly MJ. 2023. Discordant calls across genotype discovery approaches elucidate variants with systematic errors. Genome Res 33: 999–1005.

5. Barbitoff YA, Abasov R, Tvorogova VE, Glotov AS, Predeus A V. 2022. Systematic benchmark of state-of-the-art variant calling pipelines identifies major factors affecting accuracy of coding sequence variant discovery. BMC Genomics 23: 1–17. 10.1186/s12864-022-08365-3.

6. Benzer S. 1959. On the Topology of the Genetic Fine Structure. Proceedings of the National Academy of Sciences 45: 1607–1620.

7. Beyer WA, Stein ML, Smith TF, Ulam SM. 1974. A molecular sequence metric and evolutionary trees. Math Biosci 19: 9–25. https://linkinghub.elsevier.com/retrieve/pii/0025556474900285.

8. Bickhart DM, McClure JC, Schnabel RD, Rosen BD, Medrano JF, Smith TPL. 2020. Symposium review: Advances in sequencing technology herald a new frontier in cattle genomics and genome-enabled selection. J Dairy Sci 103: 5278–5290. 10.3168/jds.2019-17693.

9. Bolger AM, Lohse M, Usadel B. 2014. Trimmomatic: A flexible trimmer for Illumina sequence data. Bioinformatics 30: 2114–2120.

10. Broad Institute. 2019. Picard Toolkit. GitHub. https://broadinstitute.github.io/picard (Accessed November 24, 2023).

11. Camin JH, Sokal RR. 1965. A Method for Deducing Branching Sequences in Phylogeny. Evolution (N Y) 19: 311.

12. Chen NC, Kolesnikov A, Goel S, Yun T, Chang PC, Carroll A. 2023. Improving variant calling using population data and deep learning. BMC Bioinformatics 24.

13. Chin CS, Wagner J, Zeng Q, Garrison E, Garg S, Fungtammasan A, Rautiainen M, Aganezov S, Kirsche M, Zarate S, et al. 2020. A diploid assembly-based benchmark for variants in the major histocompatibility complex. Nat Commun 11: 1–9. 10.1038/s41467-020-18564-9.

14. Cleary JG, Braithwaite R, Gaastra K, Hilbush BS, Inglis S, Irvine SA, Jackson A, Littin R, Nohzadeh-Malakshah S, Rathod M, et al. 2014. Joint variant and de novo mutation identification on pedigrees from high-throughput sequencing data. Journal of Computational Biology 21: 405–419.

15. Cleary JG, Braithwaite R, Gaastra K, Hilbush BS, Inglis S, Irvine SA, Jackson A, Littin R, Rathod M, Ware D, et al. 2015. Comparing Variant Call Files for Performance Benchmarking of Next-Generation Sequencing Variant Calling Pipelines. bioRxiv 1–6. 10.1101/023754.

16. Clift B, Haussler D, McConnell R, Schneider TD, Stormo GD. 1986. Sequence landscapes. Nucleic Acids Res 14: 141–158.

17. Danecek P, Bonfield JK, Liddle J, Marshall J, Ohan V, Pollard MO, Whitwham A, Keane T, McCarthy SA, Davies RM. 2021. Twelve years of SAMtools and BCFtools. Gigascience 10.

18. Day A, Poplin R. 2019. Analyzing 3024 rice genomes characterized by DeepVariant. Google Cloud Blog. https://cloud.google.com/blog/products/data-analytics/analyzing-3024-rice-genomes-characterized-by-deepvariant (Accessed November 24, 2023).

19. Flack N, Drown M, Walls C, Pratte J, McLain A, Faulk C. 2023. Chromosome-level, nanopore-only genome and allele-specific DNA methylation of Pallas’s cat, Otocolobus manul. NAR Genom Bioinform 5. https://academic.oup.com/nargab/article/doi/10.1093/nargab/lqad033/7103190.

20. Genereux DP, Serres A, Armstrong J, Johnson J, Marinescu VD, Murén E, Juan D, Bejerano G, Casewell NR, Chemnick LG, et al. 2020. A comparative genomics multitool for scientific discovery and conservation. Nature 587: 240–245.

21. Guhlin J, Le Lec MF, Wold J, Koot E, Winter D, Biggs PJ, Galla SJ, Urban L, Foster Y, Cox MP, et al. 2023. Species-wide genomics of kākāpō provides tools to accelerate recovery. Nat Ecol Evol.

22. Gunderson EL, Vogel I, Chappell L, Bulman CA, Lim KC, Luo M, Whitman JD, Franklin C, Choi YJ, Lefoulon E, et al. 2020. The endosymbiont Wolbachia rebounds following antibiotic treatment. PLoS Pathog 16.

23. Hayes BJ, Daetwyler HD. 2019. 1000 Bull Genomes Project to Map Simple and Complex Genetic Traits in Cattle: Applications and Outcomes. Annu Rev Anim Biosci 7: 89–102. https://www.annualreviews.org/doi/10.1146/annurev-animal-020518-115024 (Accessed August 8, 2022).

24. Heaton MP, Smith TPL, Bickhart DM, Vander Ley BL, Kuehn LA, Oppenheimer J, Shafer WR, Schuetze FT, Stroud B, Mcclure JC, et al. 2021. A Reference Genome Assembly of Simmental Cattle, Bos taurus taurus. Journal of Heredity 112: 184– 191.

25. Ip EKK, Hadinata C, Ho JWK, Giannoulatou E. 2020. dv-trio: A family-based variant calling pipeline using DeepVariant. Bioinformatics 36: 3549–3551.

26. Jarvis ED, Formenti G, Rhie A, Guarracino A, Yang C, Wood J, Tracey A, Thibaud-Nissen F, Vollger MR, Porubsky D, et al. 2022. Semi-automated assembly of high-quality diploid human reference genomes. Nature 611: 519–531.

27. Jesudoss Chelladurai JRJ, Abraham A, Quintana TA, Ritchie D, Smith V. 2023. Comparative Genomic Analysis and Species Delimitation: A Case for Two Species in the Zoonotic Cestode Dipylidium caninum. Pathogens 12.

28. Kalbeisch T, Mckay S, Murdoch B, Adelson DL, Almansa D, Becker G, Beckett LM, José Benítez-Galeano M, Biase F, Casey T, et al. 2024. RT2T: A Global Collaborative Project to Study Chromosomal Evolution in the Suborder Ruminantia. ResearchSquare. 10.21203/rs.3.rs-3918604/v1 (Accessed February 4, 2024).

29. Kaminow B, Ballouz S, Gillis J, Dobin A. 2022. Pan-human consensus genome significantly improves the accuracy of RNA-seq analyses. Genome Res 32: 738– 750.

30. Khazeeva G, Sablauskas K, van der Sanden B, Steyaert W, Kwint M, Rots D, Hinne M, van Gerven M, Yntema H, Vissers L, et al. 2022. DeNovoCNN: a deep learning approach to de novo variant calling in next generation sequencing data. Nucleic Acids Res 50: e97.

31. Kolesnikov A, Goel S, Nattestad M, Yun T, Baid G, Yang H, McLean CY, Chang P-C, Carroll A. 2021. DeepTrio: variant calling in families using deep learning. bioRxiv 1– 16.

32. Koren S, Rhie A, Walenz BP, Dilthey AT, Bickhart DM, Kingan SB, Hiendleder S, Williams JL, Smith TPL, Phillippy AM. 2018. De novo assembly of haplotype-resolved genomes with trio binning. Nat Biotechnol 36: 1174–1182. https://www.nature.com/articles/nbt.4277.

33. Krusche P, Trigg L, Boutros PC, Mason CE, De La Vega FM, Moore BL, Gonzalez-Porta M, Eberle MA, Tezak Z, Lababidi S, et al. 2019. Best practices for benchmarking germline small-variant calls in human genomes. Nat Biotechnol 37: 555–560. 10.1038/s41587-019-0054-x.

34. Leonard AS, Crysnanto D, Mapel XM, Bhati M, Pausch H. 2023. Graph construction method impacts variation representation and analyses in a bovine super-pangenome. Genome Biol 24.

35. Leonard AS, Mapel XM, Pausch H. 2024. Pangenome genotyped structural variation improves molecular phenotype mapping in cattle. Genome Res gr.278267.123. http://genome.cshlp.org/lookup/doi/10.1101/gr.278267.123.

36. Li H. 2013. Aligning sequence reads, clone sequences and assembly contigs with BWA-MEM. ArXiv. http://arxiv.org/abs/1303.3997 (Accessed February 4, 2024).

37. Li H, Bloom JM, Farjoun Y, Fleharty M, Gauthier L, Neale B, MacArthur D. 2018. A synthetic-diploid benchmark for accurate variant-calling evaluation. Nat Methods 15: 595–597. 10.1038/s41592-018-0054-7.

38. Liao WW, Asri M, Ebler J, Doerr D, Haukness M, Hickey G, Lu S, Lucas JK, Monlong J, Abel HJ, et al. 2023. A draft human pangenome reference. Nature 617: 312–324.

39. Lin YL, Chang PC, Hsu C, Hung MZ, Chien YH, Hwu WL, Lai FP, Lee NC. 2022. Comparison of GATK and DeepVariant by trio sequencing. Sci Rep 12.

40. Low WY, Tearle R, Liu R, Koren S, Rhie A, Bickhart DM, Rosen BD, Kronenberg ZN, Kingan SB, Tseng E, et al. 2020. Haplotype-resolved genomes provide insights into structural variation and gene content in Angus and Brahman cattle. Nat Commun 11.

41. Luo R, Wong C-L, Wong Y-S, Tang C-I, Liu C-M, Leung C-M, Lam T-W. 2020. Exploring the limit of using a deep neural network on pileup data for germline variant calling. Nat Mach Intell 2: 220–227. https://www.nature.com/articles/s42256-020-0167-4.

42. McKenna A, Hanna M, Banks E, Sivachenko A, Cibulskis K, Kernytsky A, Garimella K, Altshuler D, Gabriel S, Daly M, et al. 2010. The genome analysis toolkit: A MapReduce framework for analyzing next-generation DNA sequencing data. Genome Res 20: 1297–1303.

43. Md V, Misra S, Li H, Aluru S. 2019. Efficient architecture-aware acceleration of BWA-MEM for multicore systems. In Proceedings - 2019 IEEE 33rd International Parallel and Distributed Processing Symposium, IPDPS 2019, pp. 314–324, Institute of Electrical and Electronics Engineers Inc.

44. Musolf AM, Holzinger ER, Malley JD, Bailey-Wilson JE. 2022. What makes a good prediction? Feature importance and beginning to open the black box of machine learning in genetics. Hum Genet 141: 1515–1528.

45. Nurk S, Koren S, Rhie A, Rautiainen M, Bzikadze A V, Mikheenko A, Vollger MR, Altemose N, Uralsky L, Gershman A, et al. 2022. *The complete sequence of a human genome*. https://www.science.org.

46. Olson ND, Wagner J, Dwarshuis N, Miga KH, Sedlazeck FJ, Salit M, Zook JM. 2023. Variant calling and benchmarking in an era of complete human genome sequences. Nat Rev Genet 24: 464–483.

47. Olson ND, Wagner J, McDaniel J, Stephens SH, Westreich ST, Prasanna AG, Johanson E, Boja E, Maier EJ, Serang O, et al. 2022a. PrecisionFDA Truth Challenge V2: Calling variants from short and long reads in difficult-to-map regions. Cell Genomics 2: 100129. https://linkinghub.elsevier.com/retrieve/pii/S2666979X22000581.

48. Olson ND, Wagner J, McDaniel J, Stephens SH, Westreich ST, Prasanna AG, Johanson E, Boja E, Maier EJ, Serang O, et al. 2022b. PrecisionFDA Truth Challenge V2: Calling variants from short and long reads in difficult-to-map regions. Cell Genomics 2: 100129. https://www.sciencedirect.com/science/article/pii/S2666979X22000581.

49. Oppenheimer J, Rosen BD, Heaton MP, Vander Ley BL, Shafer WR, Schuetze FT, Stroud B, Kuehn LA, Mcclure JC, Barfield JP, et al. 2021. A Reference Genome Assembly of American Bison, Bison bison bison. Journal of Heredity 112: 174–183.

50. Osipowski P, Pawełkowicz M, Wojcieszek M, Skarzyńska A, Przybecki Z, Pląder W. 2020. A high-quality cucumber genome assembly enhances computational comparative genomics. Molecular Genetics and Genomics 295: 177–193.

51. Popic V, Rohlicek C, Cunial F, Hajirasouliha I, Meleshko D, Garimella K, Maheshwari A. 2023. Cue: a deep-learning framework for structural variant discovery and genotyping. Nat Methods 20: 559–568.

52. Poplin R, Chang PC, Alexander D, Schwartz S, Colthurst T, Ku A, Newburger D, Dijamco J, Nguyen N, Afshar PT, et al. 2018a. A universal snp and small-indel variant caller using deep neural networks. Nat Biotechnol 36: 983. https://www.nature.com/articles/s41587-021-00861-3.

53. Poplin R, Ruano-Rubio V, Depristo MA, Fennell TJ, Carneiro MO, Van Der Auwera GA, Kling DE, Gauthier LD, Levy-Moonshine A, Roazen D, et al. 2018b. Scaling accurate genetic variant discovery to tens of thousands of samples. bioRxiv 1–22. 10.1101/201178.

54. Ramachandran A, Lumetta SS, Klee EW, Chen D. 2021. HELLO: improved neural network architectures and methodologies for small variant calling. BMC Bioinformatics 22: 1–31.

55. Rice ES, Koren S, Rhie A, Heaton MP, Kalbfleisch TS, Hardy T, Hackett PH, Bickhart DM, Rosen BD, Ley B Vander, et al. 2020. Continuous chromosome-scale haplotypes assembled from a single interspecies F1 hybrid of yak and cattle. Gigascience 9.

56. Rosen BD, Bickhart DM, Schnabel RD, Koren S, Elsik CG, Tseng E, Rowan TN, Low WY, Zimin A, Couldrey C, et al. 2020. De novo assembly of the cattle reference genome with single-molecule sequencing. Gigascience 9.

57. Ruperao P, Gandham P, Odeny DA, Mayes S, Selvanayagam S, Thirunavukkarasu N, Das RR, Srikanda M, Gandhi H, Habyarimana E, et al. 2023. Exploring the sorghum race level diversity utilizing 272 sorghum accessions genomic resources. Front Plant Sci 14.

58. Sahraeian SME, Liu R, Lau B, Podesta K, Mohiyuddin M, Lam HYK. 2019. Deep convolutional neural networks for accurate somatic mutation detection. Nat Commun 10.

59. Sapoval N, Aghazadeh A, Nute MG, Antunes DA, Balaji A, Baraniuk R, Barberan CJ, Dannenfelser R, Dun C, Edrisi M, et al. 2022. Current progress and open challenges for applying deep learning across the biosciences. Nat Commun 13.

60. Secomandi S, Gallo GR, Sozzoni M, Iannucci A, Galati E, Abueg L, Balacco J, Caprioli M, Chow W, Ciofi C, et al. 2023. A chromosome-level reference genome and pangenome for barn swallow population genomics. Cell Rep 42.

61. Shafin K, Pesout T, Chang PC, Nattestad M, Kolesnikov A, Goel S, Baid G, Kolmogorov M, Eizenga JM, Miga KH, et al. 2021. Haplotype-aware variant calling with PEPPER-Margin-DeepVariant enables high accuracy in nanopore long-reads. Nat Methods 18: 1322–1332.

62. Smith TPL, Bickhart DM, Boichard D, Chamberlain AJ, Djikeng A, Jiang Y, Low WY, Pausch H, Demyda-Peyrás S, Prendergast J, et al. 2023. The Bovine Pangenome Consortium: democratizing production and accessibility of genome assemblies for global cattle breeds and other bovine species. Genome Biol 24.

63. Stenløkk K, Saitou M, Rud-Johansen L, Nome T, Moser M, Árnyasi M, Kent M, Barson NJ, Lien S. 2022. The emergence of supergenes from inversions in Atlantic salmon. Philosophical Transactions of the Royal Society B: Biological Sciences 377.

64. Stephens ZD, Hudson ME, Mainzer LS, Taschuk M, Weber MR, Iyer RK. 2016. Simulating next-generation sequencing datasets from empirical mutation and sequencing models. PLoS One 11.

65. Su J, Zheng Z, Ahmed SS, Lam T-W, Luo R. 2022. Clair3-trio: high-performance Nanopore long-read variant calling in family trios with trio-to-trio deep neural networks. Brief Bioinform 23: 1–31. https://academic.oup.com/bib/article/doi/10.1093/bib/bbac301/6645484.

66. Sundaram L, Gao H, Padigepati SR, McRae JF, Li Y, Kosmicki JA, Fritzilas N, Hakenberg J, Dutta A, Shon J, et al. 2018. Predicting the clinical impact of human mutation with deep neural networks. Nat Genet 50: 1161–1170.

67. Supernat A, Vidarsson OV, Steen VM, Stokowy T. 2018. Comparison of three variant callers for human whole genome sequencing. Sci Rep 8.

68. Wagner J, Olson ND, Harris L, Khan Z, Farek J, Mahmoud M, Stankovic A, Kovacevic V, Yoo B, Miller N, et al. 2022. Benchmarking challenging small variants with linked and long reads. Cell Genomics 2: 100128. 10.1016/j.xgen.2022.100128.

69. Webb S. 2018. Deep learning for biology. Nature 554: 555–557.

70. Whalen S, Schreiber J, Noble WS, Pollard KS. 2021. Navigating the pitfalls of applying machine learning in genomics. Nat Rev Genet 23: 169–181. 10.1038/s41576-021-00434-9.

71. Yun T, McLean C, Chang P-C, Carroll A. 2018. Improved non-human variant calling using species-specific DeepVariant models. DeepVariant Blog. https://google.github.io/deepvariant/posts/2018-12-05-improved-non-human-variant-calling-using-species-specific-deepvariant-models/.

72. Zhao S, Agafonov O, Azab A, Stokowy T, Hovig E. 2020. Accuracy and efficiency of germline variant calling pipelines for human genome data. Sci Rep 10.

73. Zhou Y, Yang L, Han X, Han J, Hu Y, Li F, Xia H, Peng L, Boschiero C, Rosen BD, et al. 2022a. Assembly of a pangenome for global cattle reveals missing sequences and novel structural variations, providing new insights into their diversity and evolutionary history. Genome Res 32: 1585–1601. 10.1101/gr.276550.122.

74. Zhou Y, Yang L, Han X, Han J, Hu Y, Li F, Xia H, Peng L, Boschiero C, Rosen BD, et al. 2022b. Assembly of a pangenome for global cattle reveals missing sequences and novel structural variations, providing new insights into their diversity and evolutionary history. Genome Res 32: 1585–1601.

75. Zook JM, Chapman B, Wang J, Mittelman D, Hofmann O, Hide W, Salit M. 2014. Integrating human sequence data sets provides a resource of benchmark SNP and indel genotype calls. Nat Biotechnol 32: 246–251.

76. Zou J, Huss M, Abid A, Mohammadi P, Torkamani A, Telenti A. 2019. A primer on deep learning in genomics. Nat Genet 51: 12–18. 10.1038/s41588-018-0295-5.

