## Supplemental Materials for "Overcoming Limitations to Deep Learning in Domesticated Animals with TrioTrain"

### SUPPLEMENTAL RESULTS

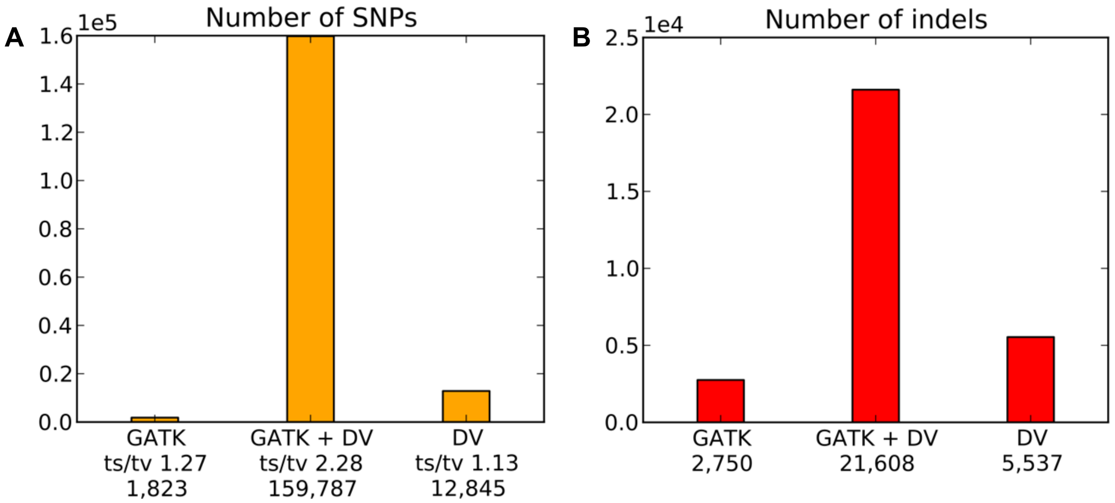

**Supplemental Figure 1. Preliminary comparison between DeepVariant and GATK.** Before pursuing re-training, we used a single bovine genome (UMAG lab ID 828) to compare between approaches. The y-axis represents raw counts with more total SNPs (A) than INDELs (B). The middle bar contains the overlapping variants between both methods, while the other two outer bars contain variants unique to each. The lower Transition to Transversion (Ts/Tv) ratio for SNPs unique to DV indicates that the 1kBulls Run8 callset workflow identified more unique high-quality novel variation than DeepVariant (v1.0). Note that the GATK results are not from the VQSR-optimized GATK workflow used to generate training labels, and an earlier version of the DV was used (1.0). Both approaches are now depreciated; however, we include these results to assist others considering re-training.

**Supplemental Note 1. Reviewing loss plots to identify appropriate filters for training data.** The loss plots produced during training with the 1kBulls Run 8 data indicated overfitting and potentially noisy labels (Supplemental Figure 2). We observed a drastic separation between the two loss curves, the optimal checkpoints occurred early in the epoch, and training performance worsened with better-quality labels. For example, the early checkpoints used between 6 and 18 percent of all labeled examples, meaning more data did not help improve the model. Combined, these contradictory results revealed that filters were not applied consistently between the truth VCF and the PopVCF.

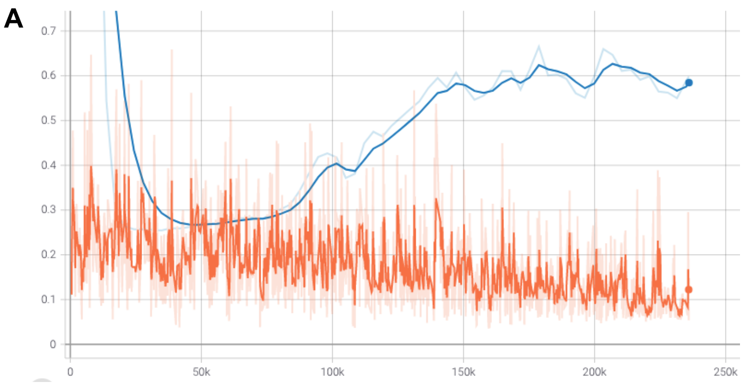

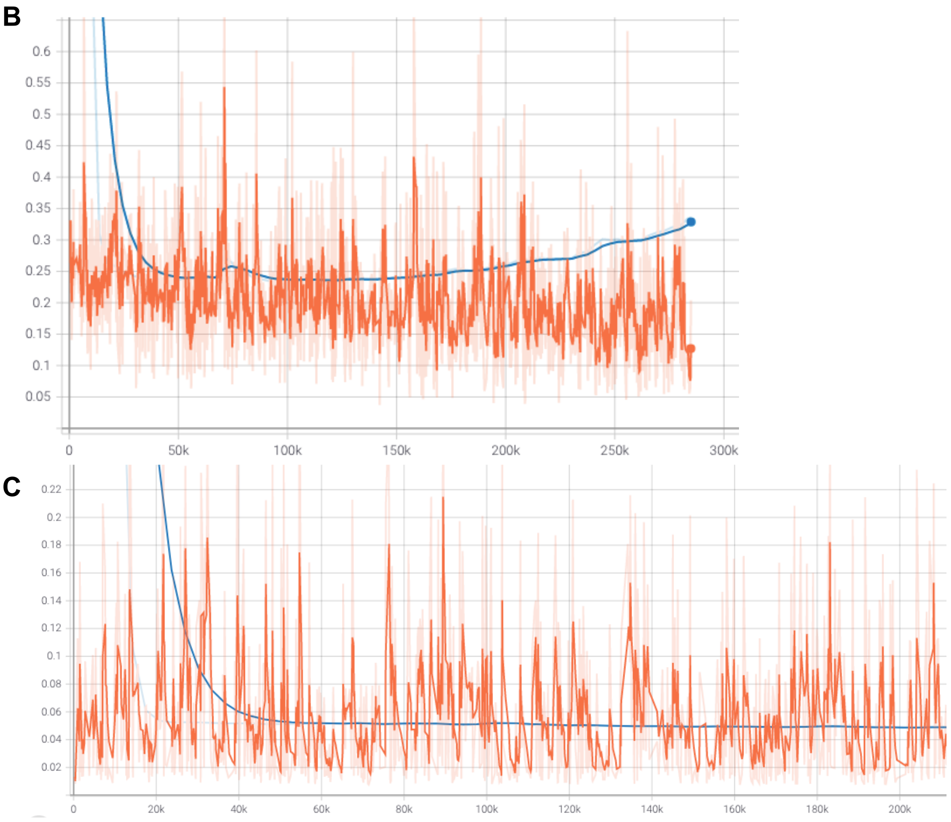

**Supplemental Figure 2. Training and Tuning Loss Plots with Different Quality Truth Labels.**
Line plots illustrating changes to model loss, as calculated by the optimization function during training of DeepVariant. The darker orange line represents the smoothed curve for loss in the parent, while the dark blue line the same curve in the offspring. The x-axis indicates the number of cumulative training steps completed when calculating metrics, where the number of labeled examples is equal to the number of steps x batch size (32). Decreasing values for loss indicate model improvement. (A) This loss plot illustrates an example of a low-quality training iteration that occurred due to mismatched filtering criteria. The filtering criteria is less strict in one dataset, and stricter in another which causes the erratic performance. The separation between training (orange) and tuning (blue) indicates overfitting. (B) This loss plot also indicates an example of low-quality training but demonstrates how we identified the filtering mismatch. Overfitting in the offspring is reduced without any filters based on QUAL or GQ value in the truth VCF. If filtering was appropriate, strict filters would achieve higher quality labels to reduce error. Instead, our initial training with the raw 1kBulls Run8 genotypes resulted in an optimal checkpoint after covering 33.5% of the examples created, whereas applying filters resulted in optimal training halting at 7.4% of the examples. Insufficient quality with truth labels results in an early optimal checkpoint model, as the noisy labels do not improve learning. (C) This loss is an example of training with the UMAGv1 callset with correct filters, as opposed to the 1kBulls Run8 data in figures A and B. Notice the lack of separation between training (orange) and tuning (blue) loss lines as more labeled examples are given to the model. The optimal checkpoint occurs after covering 76.6% of the total examples created. Additionally, note that training loss at the best checkpoint drops by an order of magnitude with improved truth genotypes (1kBulls Run8 = 0.23469; UMAGv1 = 0.01514)

**Supplementary Note 2. Evaluating concordance across 1000 Bulls Callsets**

The 1000 Bulls Project has been the flagship consortium effort in animal agriculture to aggregate genome sequence data from across the globe, standardize data processing and distribute a final variant call set to participants (Hayes and Daetwyler, 2019). Starting with Run7 in 2019, the data processing pipeline was significantly rewritten to transition to the Broad Institute's Genome Analysis Tool Kit (**GATK,** McKenna et al., 2010) and the new ARS-UCD1.2 reference (Rosen et al., 2020).Since then, there have been three genotyping runs; Run7 in 2019 (N=3818), Run8 in 2020 (N=4931), and Run9 in 2021 (N=6191). In total, 3,659 samples are present in all three genotyping runs. Using our assumed truth set (Run 9), we can compare Run7 to Run8, Run8 to Run9, and Run7 to Run9, calculating pairwise recall and precision metrics. To determine the consistency of variant calls over time, we developed a pipeline to extract data for these samples using only variants that PASS VQSR filtering, process through hap.py software, and summarize results (Krusche et al., 2019). Without an established ground truth for cattle, we are only able to measure variability between runs using metrics (recall and precision) as quantities. However, most researchers consider the genotypes generated for a sample to be constant across 1kBulls runs. Instead, our results indicate significantly lower recall and precision within the 1000 Bulls data than expected (Supplemental Table 1).The variability in genotypes is due to differences in analysis parameters between runs and the inclusion of additional information (more individuals), among other factors.

**Supplemental Table 1**. The average recall, precision, and F1 score for 3,659 samples within multiple 1000 Bulls genotyping runs. Values below one indicate False Negatives (recall) and False Positives (precision), where perfect calls would have values of unity. Typically, recall and precision values exceeding 0.99 are considered high quality.

| | **SNP** | | | **INDEL** | | |
|---|---|---|---|---|---|---|
| **TRUTH - QUERY** | Recall | Precision | F1 | Recall | Precision | F1 |
| **8-7** | 0.9605 | 0.9619 | 0.9612 | 0.9566 | 0.9753 | 0.9658 |
| **9-8** | 0.9462 | 0.9733 | 0.9595 | 0.9161 | 0.8468 | 0.8799 |
| **9-7** | 0.9295 | 0.9575 | 0.9432 | 0.8975 | 0.8459 | 0.8707 |

**Supplemental Table 2. Comparing species-specific training approaches.** Comparison of the training, validation, and testing data splits to create species-specific DeepVariant models according to inconsistently reported methods. We note that all bovine models benefit from using v1.4.0 of DeepVariant as a starting point; models built with more examples tend to be more accurate. Mosquitoes, Kākāpō, and cattle lack a GIAB-quality truth set; however, the non-human species have a bigger sample size. Thus, the raw data were split by sample rather than by chromosome, producing examples from the autosomes and the X chromosome (mosquitos: chr 2-3L/R + X; cattle: chr 1-29 + X, Kākāpō: unreported). In contrast, the human models use sequencing replicates to supplement sample size and produce examples for a subset of the autosomes (human: training = chr1-19, validation = chr21-22, testing = chr20, HG003 left out). The deprecated models selected the optimal checkpoint based on minimal validation loss, while v1.4 uses the highest validation F1-score.

| | | | | | **Sample Size** | | |
|---|---|---|---|---|---|---|---|
| *Version* | *Species* | *Model* | *Additional Channels* | *Cumulative Examples* | *Training* | *Evaluation* | *Testing* |
| *0.7* | Human | default | - | 158,571,078 | 2 offspring | 2 offspring | 2 offspring |
| | Mosquito | default | - | NA | 5 offspring | 1 offspring | 2 parents, 1 offspring |
| *0.9* | Human | default | - | 325,202,093 | 2 offspring | 2 offspring | 2 offspring |
| | Kākāpō * | default | - | 478,202,093 | 169◇‡ | NA | 169◇ |
| *1.4* | Human | DeepTrio-Parent | insert_size | 457,374,516 | 4 parents, 2 offspring | 4 parents, 2 offspring | 2 parents, 1 offspring |
| | Human | DeepTrio-Child | insert_size | 704,228,446 | 6 parents, 3 offspring | 6 parents 3 offspring | 2 parents, 1 offspring |
| | Human | WGS.AF | insert_size, allele_freq | 517,209,566 | ** | ** | ** |
| | Human | default | insert_size | 517,209,566 | 3 parents, 3 offspring | 3 parents, 3 offspring | 3 parents, 3 offspring |
| | Cattle | Phase1 | insert_size, allele_freq | 560,587,902 | 12 parents | 6 offspring | 19 cattle, 6 humans |
| | Cattle | Phase2 | insert_size, allele_freq | 584,973,310 | 18 parents | 9 offspring | 19 cattle, 6 humans |
| | Cattle | Phase3 | insert_size, allele_freq | 676,487,582 | 22 parents | 11 offspring | 19 cattle, 6 humans |
| | Cattle | Phase4 | insert_size, allele_freq | 740,019,358 | 28 parents | 14 offspring | 16-18 cattle, 6 humans |

| | | | | | | |
|---|---|---|---|---|---|---|
| Cattle | Phase5 | insert_size, allele_freq | 760,098,558 | 30 parents | 15 offspring | 15 cattle, 6 humans |

** Number of examples is estimated from batch size (306) and approximate number of training steps (~500,000)* (Guhlin et al. 2023)*)*
*‡ SNP genotypes were altered by correcting Mendelian discordant SNPs from a DV-derived callset (v0.9) based on 16 nuclear families (~80 individuals)* (Guhlin et al. 2023)
*◇ Represents the nearly the entire Kākāpō population.*
*** Identical model to default, but with another channel encoding 1000Genomes Allele Frequencies*

**Supplemental Table 3. Excel File, Separate Attachment**

**Supplemental Table 4. Training performance.** The table below describes the F1-score values, stratified by genotype class and variant type. Note that the metric for model selection (F1/All) was identical to F1/HomRef. We observe a lower F1-score with INDELs, which are under-represented in our truth labels due to relying on SRS data for label curation.

| | **Trio** | | | | **F1-Score** | | | | |
|---|---|---|---|---|---|---|---|---|---|
| *Breed* | *Name* | *Iteration* | *Training Genome* | *Steps Covered* | *HomRef* | *Het* | *HomAlt* | *SNPs* | *INDELs* |
| | | | | ***Phase 1*** | | | | | |
| *Angus (AA)* | Trio1 | 1 | Sire | 84,518 | 0.992378 | 0.990173 | 0.995597 | 0.993990 | 0.981645 |
| | | 2 | Dam | 201,011 | 0.992497 | 0.990754 | 0.995881 | 0.994061 | 0.982073 |
| *Hereford (HE)* | Trio2 | 3 | Sire | 134,175 | 0.980533 | 0.978597 | 0.992152 | 0.989170 | 0.919398 |
| | | 4 | Dam | 77,316 | 0.980109 | 0.977662 | 0.992260 | 0.989187 | 0.916280 |
| *Brown Swiss (BS)* | Trio3 | 5 | Sire | 76,110 | 0.986626 | 0.986306 | 0.993298 | 0.991148 | 0.952780 |
| | | 6 | Dam | 99,664 | 0.986563 | 0.986160 | 0.993366 | 0.991106 | 0.952563 |
| *Holstein (HO)* | Trio4 | 7 | Sire | 15,507 | 0.987717 | 0.986423 | 0.992806 | 0.991310 | 0.961992 |
| | | 8 | Dam | 113,822 | 0.988590 | 0.987312 | 0.993216 | 0.991775 | 0.965884 |
| *Hereford (HE)* | Trio5 | 9 | Sire | 127,591 | 0.980117 | 0.975715 | 0.993709 | 0.989418 | 0.915555 |
| | | 10 | Dam | 63,019 | 0.980370 | 0.976392 | 0.993673 | 0.989498 | 0.916762 |
| *Tyrolean Grey (TG)* | Trio6 | 11 | Sire | 159,745 | 0.991886 | 0.989977 | 0.996388 | 0.993501 | 0.980971 |
| | | 12 | Dam | 203,095 | 0.991871 | 0.989765 | 0.996361 | 0.993499 | 0.980877 |
| | | | | ***Phase 2*** | | | | | |
| *Holstein-Jersey (HJ)* | Trio7 | 13 | Sire | 101,101 | 0.988295 | 0.984281 | 0.995218 | 0.993712 | 0.951608 |

| | | | | | | | | | |
|---|---|---|---|---|---|---|---|---|---|
| | | 14 | Dam | 120,966 | 0.988345 | 0.984364 | 0.995407 | 0.993740 | 0.951701 |
| *Holstein-Jersey (HJ)* | Trio8 | 15 | Sire | 225,421 | 0.988391 | 0.986733 | 0.993765 | 0.991686 | 0.965026 |
| | | 16 | Dam | 116,248 | 0.988385 | 0.986736 | 0.993737 | 0.991724 | 0.964678 |
| *Holstein-Hereford (HH)* | Trio9 | 17 | Sire | 77,464 | 0.978856 | 0.975292 | 0.992142 | 0.987511 | 0.917724 |
| | | 18 | Dam | 120,844 | 0.979315 | 0.976319 | 0.992190 | 0.987687 | 0.920014 |
| | | | | ***Phase 3*** | | | | | |
| *Bison (BI)* | Trio10 | 19 | Sire | 228,738 | 0.989776 | 0.995272 | 0.979898 | 0.992136 | 0.967921 |
| | | 20 | Dam | 892,234 | 0.990404 | 0.995668 | 0.980929 | 0.992639 | 0.969567 |
| *Bison (BI)* | Trio11 | 21 | Sire | 803,481 | 0.989700 | 0.995110 | 0.980646 | 0.992271 | 0.966043 |
| | | 22 | Dam | 935,368 | 0.989998 | 0.995659 | 0.980880 | 0.992370 | 0.968051 |
| | | | | ***Phase 4*** | | | | | |
| *Angus/Brahman F1 (AA/BR)* | Trio12 | 23 | Sire | 59,354 | 0.980774 | 0.979656 | 0.990326 | 0.984428 | 0.950244 |
| | | 24 | Dam | 217,570 | 0.980420 | 0.980100 | 0.989698 | 0.983825 | 0.952273 |
| *Yak/Highlander F1 (YK/HI)* | Trio13 | 25 | Sire | 275,488 | 0.986110 | 0.954403 | 0.997429 | 0.988671 | 0.962370 |
| | | 26 | Dam | 432,983 | 0.986396 | 0.955538 | 0.997896 | 0.988876 | 0.963399 |
| *Bison/Simmental F1 (BI/SI)* | Trio14 | 27 | Sire | 204,026 | 0.978195 | 0.949967 | 0.992479 | 0.982472 | 0.936515 |
| | | 28 | Dam | 282,383 | 0.982670 | 0.948754 | 0.996569 | 0.986763 | 0.942704 |
| | | | | ***Phase 5*** | | | | | |
| *Synthetic AA/BR F1* | Trio15 | 29 | Sire | 290,765 | 0.948610 | 0.953821 | 0.968833 | 0.960721 | 0.851486 |
| | | 30 | Dam | 336,710 | 0.979190 | 0.955451 | 0.995209 | 0.990430 | 0.890876 |
| **MEAN** | | | | | 0.984436 | 0.978945 | 0.991399 | 0.989311 | 0.947299 |

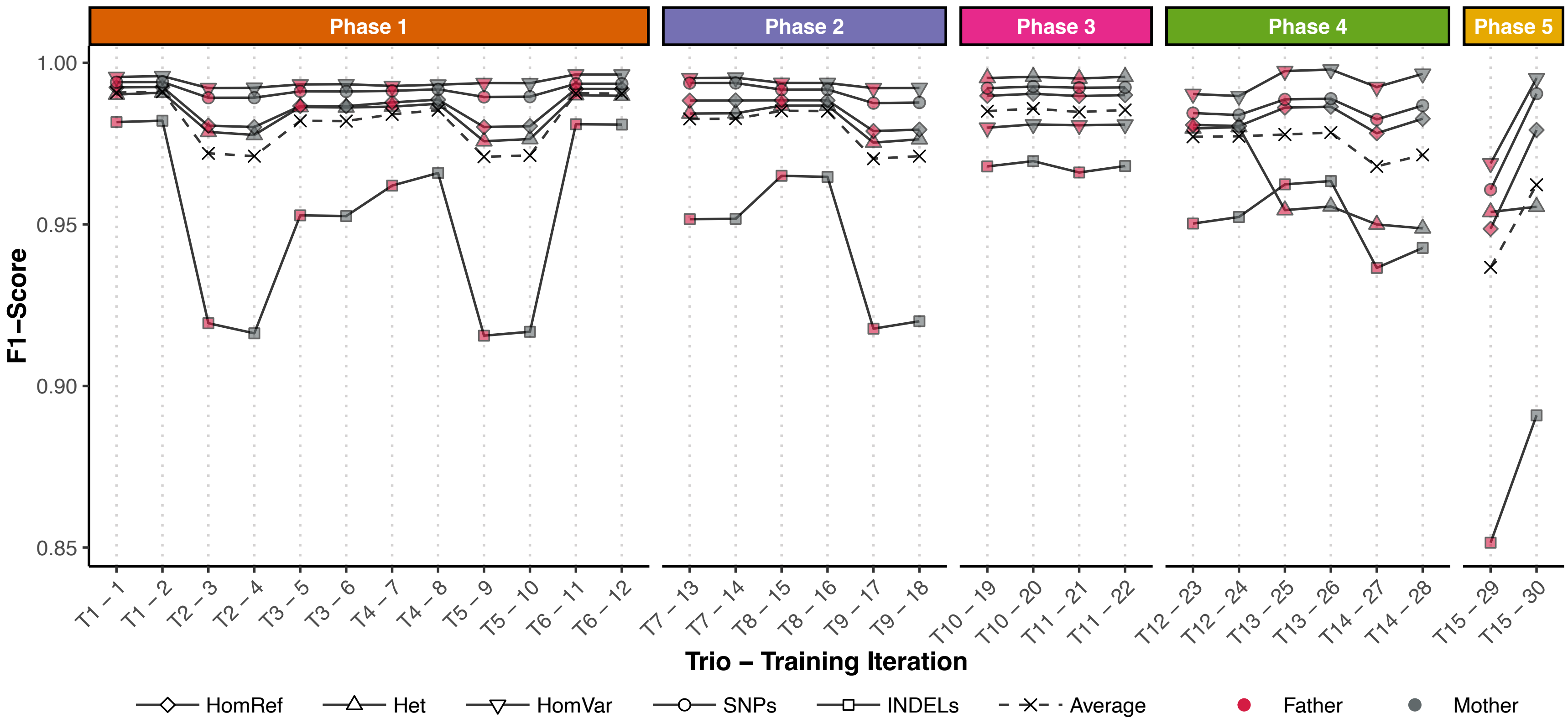

**Supplemental Figure 3. Model training performance across all phases.** Each box-and-whisker represents the F1-score with the GIAB human trios (n = 6). Phase 5 is from training on the same parental genomes in the first two iterations of Phase 4; however, the offspring used for model tuning was replaced with a synthetic diploid (SynDip), created by sampling from the reconstructed parental haplotype assemblies and merged. As the assemblies have a higher per-base error rate than the real Illumina sequencing data used during Phase 4, we observe a large shift in performance due to differing error rates between training and tuning datasets. We find that SynDip samples created from current bovine assemblies with moderate per-base error rates are not ideal for training data, unless the aim is to use a model to call variants in synthetic data. As such, we halted any further training with our synthetic sample

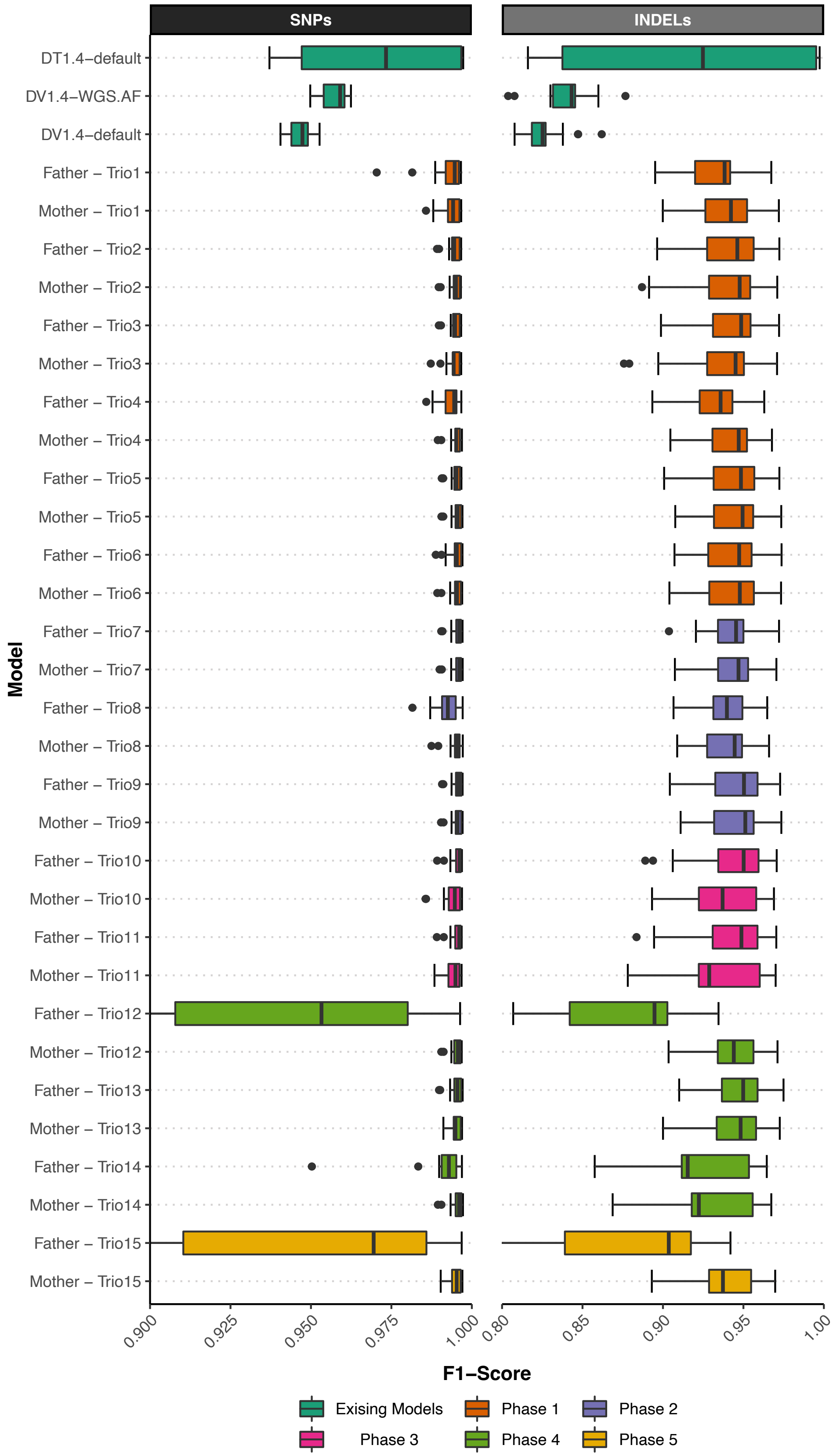

SNPs
INDELs
DT1.4−default
DV1.4−WGS.AF
DV1.4−default
Father – Trio1
Mother – Trio1
Father – Trio2
Mother – Trio2
Father – Trio3
Mother – Trio3
Father – Trio4
Mother – Trio4
Father – Trio5
Mother – Trio5
Father – Trio6
Mother – Trio6
Father – Trio7
Mother – Trio7
Father – Trio8
Mother – Trio8
Father – Trio9
Mother – Trio9
Father – Trio10
Mother – Trio10
Father – Trio11
Mother – Trio11
Father – Trio12
Mother – Trio12
Father – Trio13
Mother – Trio13
Father – Trio14
Mother – Trio14
Father – Trio15
Mother – Trio15
Model
0.900
0.925
0.950
0.975
1.000
0.80
0.85
0.90
0.95
1.00
F1−Score
Exising Models
Phase 1
Phase 2
Phase 3
Phase 4
Phase 5

**Supplemental Figure 4. Comparing variants from bovine test genomes across all phases. (**previous page) Each box-and-whisker represents the F1-score with an independent set of bovine samples previously unseen by the model (n = 19 for all models prior to phase 4, except DT where n = 6; in phase 4, sample size decreases, n-1, for each trio resulting in n = 16 in the final test set). The x-axis scale begins at 0.9 and 0.8 for SNPs and INDELs, respectively, where an F1-score of 1 would indicate perfect prediction. Phase 5 is from training on the identical parental genomes in the first two iterations of Phase 4; however, the offspring used for model tuning was replaced with a synthetic diploid (SynDip), created by sampling from the reconstructed parental haplotype assemblies and merged. Note that iteration 29 (Father-Trio15) is the first time the model was given examples from a synthetic diploid hybrid-cross offspring. This unique case extends the whisker SNPs out of view (minimum F1-Score = 0.863). As the assemblies have a higher per-base error rate than the real Illumina sequencing data used during Phase 4, we observe a sizeable change in performance due to the model adjusting to the new error profile. As we do not intend to genotype synthetic reads, we halted any further training with our synthetic samples.

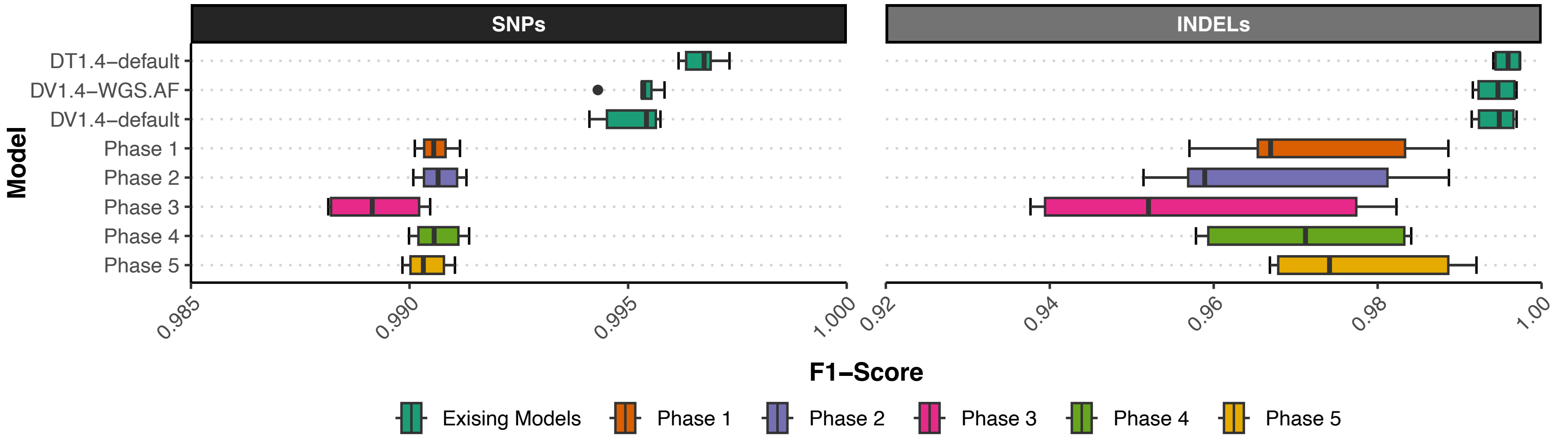

**Supplemental Figure 5. Comparing variants from human genomes.** Each box-and-whisker represents the F1-score with the GIAB human trios (n = 6). Note that the x-axis scale begins at 0.985 and 0.94 for SNPs and INDELs, respectively, where an F1-score of 1 would indicate perfect prediction. We observe a slight reduction in SNP F1-score for Phase 5 due to the higher per-base error rate in the synthetic reads during testing. However, INDEL performance improves further, demonstrating that with a more balanced set of truth labels, we may recover some of the INDEL performance lost by training with truth labels based on real SRS data, which is less sensitive to INDELs.

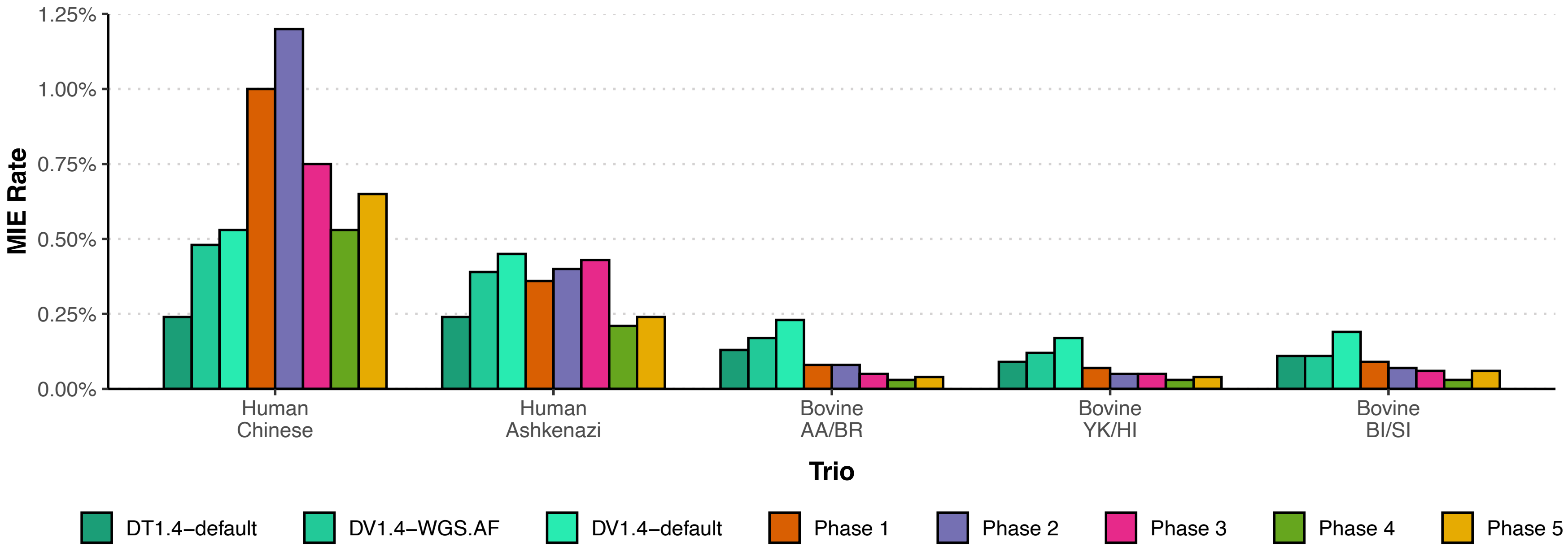

**Supplemental Figure 6. Inheritance error rate in human and bovine trios.** The above bar plot contains the number of discordant variants proportional to the total number of variants analyzed. Each group represents the offspring's PASS variants, which violate Mendelian inheritance expectations, with each bar representing a unique model checkpoint, with the addition of the Phase 5 checkpoint. As expected, training with synthetic data created a model checkpoint with less genotyping accuracy for all trios and halted any further training.

### SUPPLEMENTAL METHODS
### UMAG WGS Processing Methods

#### Supplemental Note 3. Public Samples from SRA

Relevant samples are identified from NCBI SRA based on taxon ID, platform, layout and sequencing strategy. For example:

```
https://www.ncbi.nlm.nih.gov/sra/?term=(txid9913[All
Fields])+AND+((("illumina"[Platform]) AND "paired"[Layout]) AND
"wgs"[Strategy])
```

The RunInfo report is stored in a local database and samples are selected by aggregating runs based on BioSampleID and selecting samples with at least 3-fold raw sequence coverage. SRA runs are downloaded using the SRA Toolkit for each Run. The SRA Toolkit versions ranged from 2.9.1 to 2.11.0 as new data are downloaded periodically over time. (https://github.com/ncbi/sra-tools)

```
prefetch --progress --max-size 200G $Run
```

The prefetched data are then converted to fastq using either fastq-dump or fasterq-dump.

```
fastq-dump --split-e --outdir $CWD $Run
fasterq-dump -f -p -e $DumpThreads --split-3 --outdir $CWD $Run
```

The resulting fastq files are renamed to our UMAG nomenclature and compressed with pigz (https://zlib.net/pigz/).

#### Supplemental Note 4. Sequence Data Processing

Data are processed in stages from FASTQ to final analysis ready BCF following GATK Best Practices (Depristo et al. 2011; Van der Auwera and O'Conn) and implemented using various custom Perl scripts. Each stage is described below, and a representative command line is shown. As data have been continuously processed over time, starting in 2018, exact software versions have varied. However, the general workflow has been consistent and only specific changes based on software version options have been varied.

#### Supplemental Note 5. Sequence Quality Control

Low quality bases and adapters are removed using trimmomatic (Bolger et al. 2014) (https://github.com/usadellab/Trimmomatic) Versions 0.36 to 0.39.

```
java -Djava.io.tmpdir=${cwd}/tmp -XX:+UseParallelGC -
XX:ParallelGCThreads=2 -jar $Trimmomatic \
PE -phred${InputQV} \
-threads $Cpu_Trim \
-summary ${TrimOutBaseFile}.TRIM.SUMMARY \
$for_file $rev_file \
${TrimOutBaseFile}.1.P.fq ${TrimOutBaseFile}.1.U.fq \
${TrimOutBaseFile}.2.P.fq ${TrimOutBaseFile}.2.U.fq\
MINLEN:35 TOPHRED33
ILLUMINACLIP:${RefGenome}/${AdapterFile}:2:30:6:1:TRUE \
LEADING:20 TRAILING:20 SLIDINGWINDOW:3:15 AVGQUAL:20 MINLEN:35
```

**Supplemental Note 6. Reference Genome**
The reference genome used is the same as that used by the 1000 Bull Genomes project(Hayes and Daetwyler 2018), ARS-UCD1.2_Btau5.0.1Y. Briefly, because the reference animal was female, the autosomes, X chromosome, mitochondria and unplaced contigs from GCA_002263795.2 (Rosen et al. 2020a) were merged with the Y chromosome from GCA_000003205.6 (Elsik et al. 2009) which is the sire of the reference animal.

**Supplemental Note 7. Genome Alignment**
FASTQ data passing QC was aligned to the reference genome initially with BWA MEM (https://github.com/lh3/bwa) and later with BWA MEM2 (https://github.com/bwa-mem2/bwa-mem2) after appropriate reference genome indexing. During alignment, the appropriate read group tags are populated to facilitate downstream processing.

```
$bwa mem -M -t $Cpu_Align -R
'\@RG\\tID:${base_file}\\tSM:${animal_id}\\tLB:${lib}\\tPL:ILLUMI
NA' $BWA_ref $for_file $rev_file >$bwa_output
```

**Supplemental Note 8. Sort/Merge/Mark Duplicates**
The resulting SAM files are sorted using samtools (https://github.com/samtools/) resulting in a list of sorted BAM files which are subsequently merged using picard (https://github.com/broadinstitute/picard) and duplicates marked using Picard. Importantly, MarkDuplicates is performed in a "read group aware" manner using the appropriate optical pixel distance for traditional flow cells (100) or patterned flow cells (2500) based on the source of the data. For samples that consisted of multiple sequencing libraries, the resulting library specific BAM files are merged and indexed to produce a single BAM file using samtools.

```
$Samtools sort -m ${SortMem}G -\@ $Cpu_Samtools -o $sort_output -
T ${sam_input}.TMP $sam_input

java -Djava.io.tmpdir=${cwd}/tmp -XX:+UseParallelGC -
XX:ParallelGCThreads=2\
-Xmx20g -jar $Picard MergeSamFiles @in
OUTPUT=${BAM_PREFIX}.${_}.merged.bam\
USE_THREADING=TRUE MERGE_SEQUENCE_DICTIONARIES=TRUE
ASSUME_SORTED=TRUE\
VALIDATION_STRINGENCY=LENIENT TMP_DIR=${cwd}/tmp

java -Djava.io.tmpdir=${cwd}/tmp -XX:+UseParallelGC -
XX:ParallelGCThreads=6 \
-Xmx50g -jar $Picard MarkDuplicates \
-INPUT ${BAM_PREFIX}.${_}.merged.bam -OUTPUT
${BAM_PREFIX}.${_}.bam \
-METRICS_FILE ${BAM_PREFIX}.${_}.DUP.METRICS -MAX_RECORDS_IN_RAM
5000000 \
-MAX_FILE_HANDLES_FOR_READ_ENDS_MAP 1000 -ASSUME_SORTED TRUE \
-VALIDATION_STRINGENCY LENIENT -TMP_DIR ${cwd}/tmp \
-OPTICAL_DUPLICATE_PIXEL_DISTANCE ${Pixels} -COMPRESSION_LEVEL 0

$Samtools merge -\@ $Cpu_Samtools -p -f -b
${BAM_PREFIX}_MERGE_files.txt ${BAM_PREFIX}.bam
```

```
$Samtools index -\@ 8 ${BAM_PREFIX}.bam
```

**Supplemental Note 9. INDEL Realignment**

While indel realignment was removed from the Best Practices, we have elected to retain it as part of our normal processing in order to facilitate other analyses not performed in the current work. We process each chromosome independently to generate indel targets, realign and then merge all chromosome back into a single indexed bam file. The GATK version used remained constant 3.8-1-0-gf15c1c3ef.

```
java -Djava.io.tmpdir=${cwd}/tmp -XX:+UseParallelGC -
XX:ParallelGCThreads=2 \
-Xmx8g -jar $GATK -nt $Cpu_RTC -T RealignerTargetCreator \
-R ${RefGenome}/${ref}.fa -L $_ -I ${BAM_PREFIX}.bam \
-o ${BAM_PREFIX}.${_}.forIndelRealigner.intervals

java -Djava.io.tmpdir=${cwd}/tmp -XX:+UseParallelGC -
XX:ParallelGCThreads=2 \
-Xmx${JavaMem}g -jar $GATK -R ${RefGenome}/${ref}.fa -I
${BAM_PREFIX}.bam \
-T IndelRealigner \
-targetIntervals ${BAM_PREFIX}.${_}.forIndelRealigner.intervals \
-L $_ -o ${BAM_PREFIX}.${_}.${BAM_SUFFIX}.bam --maxReadsInMemory
300000

$Samtools merge -\@ $Cpu_Samtools -f -c -p \
-b ${BAM_PREFIX}_MergeRealignedFiles.list
${BAM_PREFIX}.${BAM_SUFFIX}.bam
$Samtools index -\@ 8 ${BAM_PREFIX}.${BAM_SUFFIX}.bam
```

**Supplemental Note 10. Base Quality Score Recalibration (BQSR)**

Sequencing data were generated over the course of many years and many platforms, thus requiring base quality recalibration. The initial known variant file used was that from the 1000 Bull Project and subsequently modified over time to account for additional data. Briefly, the initial known variant file was constructed based on remapping variants from dbSNP build 150, containing approximately 104 million variants, onto the new UCD1.2_Btau5.0.1Y reference genome. Additional unique variants were added from 1000 Bull Project Runs7 and Run8 based on Taurus and Indicus genomes. Additionally, variants from closely related species such as Bison, Gaur, Banteng and Yak (referred to as outgroups below) were also added. The final file contains 174.3 M SNPs and 22.9 M INDELs (UMAG_9913_BQSR_ARS1.2_V4.vcf.gz). Over the course of testing, it was determined that the outgroup species were having their base qualities over-corrected, likely due to their genetic distance from the Hereford (*Bos taurus*) reference genome. It was determined that setting the following parameters alleviated the over-correction while not adversely affecting *Bos taurus* or *Bos indicus* samples:

```
--bqsrBAQGapOpenPenalty 45
--deletions_default_quality 45
--insertions_default_quality 45
```

Optimal BQSR is obtained by building a recalibration model using the entire genome and all available information, however this requires lengthy run times. Extensive testing was performed to evaluate using smaller intervals to build the model. Because the quality of the recalibration model is dependent on the amount of data provided, we reasoned that a sufficient model could be obtained using smaller genomic intervals based on the total amount of sequencing data available for a sample. Given that the Bos taurus genome is approximately 3 Gb in size and contains 29 autosomes plus the X chromosome, we developed “target” intervals of size 5 Mb, 10 Mb and 20 Mb for each of the 30 chromosomes starting at position 1 Mb on each chromosome. We developed “testing” intervals of the same size starting at position 20 Mb. The “target” intervals are used to build the recalibration model used for BQSR. The “testing” intervals are used to generate a new recalibration model after the data have been recalibrated and used to evaluate the overall quality before and after recalibration. Because the on-disk size of a BAM file is proportional to the total amount of sequence data contained within the file, we determine the interval size to use for BQSR based on the BAM file size where files <10 GB use the entire genome, <35 GB use 20 Mb, <70 GB use 10 Mb and >70 GB use 5 Mb target/testing intervals. This results in similar amounts of data used for model generation/evaluation for each sample while significantly decreasing run time. Through extensive testing (data not shown) we determined that the reduced intervals based on total data amount produced similar results to using the full genome. Importantly, because our target and testing intervals are distinct, a well-calibrated testing model indicates that the model used for the entire genome was also well calibrated. GATK BaseRecalibrator and PrintReads used GATK version 3.8-1-0-gf15c1c3ef. PrintReads is run in parallel by chromosome and chromosomes are merged and indexed. GATK version 4.1.9.0 was used for AnalyzeCovariate plots.

```
java -Djava.io.tmpdir=${cwd}/tmp -XX:+UseParallelGC -
XX:ParallelGCThreads=2 \
-Xmx20g -jar $GATK -nct $Cpu_BQSR -T BaseRecalibrator \
-R ${RefGenome}/${ref}.fa -I ${BAM_PREFIX}.${BAM_SUFFIX}.bam \
-L ${RefSNP}/${snp_dir}/BQSR_${BqsrSize}MB_target.interval_list \
-knownSites ${RefSNP}/${snp_dir}/${KnownSites} \
--bqsrBAQGapOpenPenalty $bqsrBAQGOP --deletions_default_quality
$indelQUAL \
--insertions_default_quality $indelQUAL \
-o ${BAM_PREFIX}.recalibration_report.grp \
-U ALLOW_N_CIGAR_READS --quantizing_levels 24

java -Djava.io.tmpdir=${cwd}/tmp -XX:+UseParallelGC -
XX:ParallelGCThreads=2 \
-Xmx12g -jar $GATK -nct $Cpu_PR -T PrintReads -R
${RefGenome}/${ref}.fa \
-L $_ -I ${BAM_PREFIX}.${BAM_SUFFIX}.bam \
-BQSR ${BAM_PREFIX}.recalibration_report.grp \
-o ${BAM_PREFIX}.${_}.${BAM_SUFFIX}.recalibrated.bam -U
ALLOW_N_CIGAR_READS

$Samtools merge -\@ $Cpu_Samtools -l 9 -f -c -p -b \
${BAM_PREFIX}_MergeRealignedRecalibratedFiles.list \
${BAM_PREFIX}.$BAM_SUFFIX.bam

$Samtools index -\@ 4 ${BAM_PREFIX}.${BAM_SUFFIX}.bam
```

```
java -Djava.io.tmpdir=${cwd}/tmp -XX:+UseParallelGC -
XX:ParallelGCThreads=2 \
-Xmx10g -jar $GATK4 AnalyzeCovariates \
-before ${BAM_PREFIX}.recalibration_report.grp \
-after ${BAM_PREFIX}.recalibration_report2.grp -plots
${BAM_PREFIX}.BQSR3.pdf
```

**Supplemental Note 11. Summary Metrics**
We collect summary metrics using GATK DepthOfCoverage and Picard CollectMultipleMetrics. The output from these are reformatted and stored within our database for downstream QC and summary metrics.

DepthOfCoverage
```
java -Djava.io.tmpdir=${cwd}/tmp -XX:+UseParallelGC -
XX:ParallelGCThreads=2 \
-Xmx10g -jar $GATK -nt $Cpu_DOC -R ${RefGenome}/${ref}.fa \
-I ${BAM_PREFIX}.${BAM_SUFFIX}.bam -T DepthOfCoverage -L $_ \
-o ${BAM_PREFIX}.${_}.${BAM_SUFFIX}.bam.coverage -omitBaseOutput
\
-omitIntervals --omitLocusTable -ct 5 -ct 10 -ct 15 -ct 20 -ct 25
\
-ct 30 -ct 40 -ct 50 -ct 80 -ct 90 -ct 100 -ct 150 --
minBaseQuality 15 \
--minMappingQuality 30 --start 1 --stop 1000 --nBins 999 -dt NONE
\
-U ALLOW_N_CIGAR_READS
```

CollectMultipleMetrics
```
java -Djava.io.tmpdir=${cwd}/tmp -XX:+UseParallelGC -
XX:ParallelGCThreads=2 \
-Xmx10g -jar $Picard CollectMultipleMetrics \
-INPUT ${BAM_PREFIX}.${BAM_SUFFIX}.bam -OUTPUT ${BAM_PREFIX} \
-METRIC_ACCUMULATION_LEVEL LIBRARY \
-REFERENCE_SEQUENCE ${RefGenome}/${ref}.fa\
-PROGRAM CollectAlignmentSummaryMetrics \
-PROGRAM CollectInsertSizeMetrics \
-STOP_AFTER $CollectMetricsNumReads \
-TMP_DIR ${cwd}/tmp -ASSUME_SORTED TRUE
```

**Supplemental Note 12. HaplotypeCaller & CallableRegions**
GATK HaplotypeCaller is used to generate gVCF files for each sample and each chromosome in parallel. Based on initial estimates from the Bovine HapMap project (Gibbs et al. 2009) we set the `--heterozygosity` parameter to 0.0015. We also generate CallableRegions for each chromosome to facilitate downstream applications.

```
java -Djava.io.tmpdir=${cwd}/tmp -XX:+UseParallelGC -
XX:ParallelGCThreads=2 \
-Xmx15g -jar $GATK -nct $Cpu_HC -ERC GVCF -T HaplotypeCaller \
-R ${RefGenome}/${ref}.fa -L $_ -I
${BAM_PREFIX}.${BAM_SUFFIX}.bam \
-o ${BAM_PREFIX}.${_}.g.vcf.gz --heterozygosity $Heterozygosity \
```

```
--pcr_indel_model NONE --useNewAFCalculator

java -Djava.io.tmpdir=${cwd}/tmp -XX:+UseParallelGC -
XX:ParallelGCThreads=2 \
-Xmx15g -jar $GATK -T CallableLoci -R ${RefGenome}/${ref}.fa -L
$_ \
-I ${BAM_PREFIX}.${BAM_SUFFIX}.bam \
-summary ${BAM_PREFIX}.CallableLoci.${_}.summary.txt \
-o ${BAM_PREFIX}.CallableLoci.${_}.bed
```

**Supplemental Note 13. Joint Genotype Calling**

An initial set of 5,612 genomes were available to use for joint genotype calling. The gVCF files, for each chromosome, were combined using groups of 100 samples resulting in 57 gVCF files per chromosome. These combined gVCF files are then passed as a list to GenotypeGVCs for joint genotype calling. In order to reduce total run time, chromosomes are genotyped in 10 Mb chunks in parallel and then concatenated/merged back into a single vcf per chromosome and finally indexed using bcftools (GATK 3.8-1-0-gf15c1c3ef, bcftools 1.14).

```
java -Djava.io.tmpdir=$cwd/tmp -XX:ParallelGCThreads=2 -Xmx20g -
jar $GATK \
-T CombineGVCFs -R ${RefGenome}/${ref}.fa -V
${InputFile}.${i}.${Chr}.list \
-o ${cwd}/gvcf/${Cohort}.${i}.${Chr}.g.vcf.gz

java -Djava.io.tmpdir=$tmpdir -XX:+UseParallelGC -
XX:ParallelGCThreads=2 \
-Xmx${Mem_Node}g -jar $GATK -nt $Cpu_Node -T GenotypeGVCFs \
-R ${RefGenome}/${ref}.fa -V ${Cohort}.${Chr}.list \
-L ${Chr}:${StartPosition}-${EndPosition} \
-o $outdir/${Cohort}.${Chr}.${IntervalChunk}.vcf.gz \
--heterozygosity ${Heterozygosity} --useNewAFCalculator \
--standard_min_confidence_threshold_for_calling 5

bcftools concat --naive --file-list CAT.${c}.list -Oz9 -o
${COHORT}.$c.vcf.gz
bcftools index --tbi ${COHORT}.$c.vcf.gz
```

We create SitesOnly files for downstream use with Variant Quality Score Recalibration using GATK 4.2.6.1.

```
$GatkBinary MakeSitesOnlyVcf --COMPRESSION_LEVEL 9 \
-I vcf/${YYMMDD}.${CNAME}.$c.vcf.gz \
-O vcf/SitesOnly/${YYMMDD}.${CNAME}.$c.vcf.gz
```

**Supplemental Note 14. Variant Quality Score Recalibration (VQSR)**

The effectiveness of VQSR is dependent on the overall quality and quantity of the variants used as truth sets, which are generally lacking for non-model organisms. Additionally, settings for appropriate priors for each of the truth sets should be optimized in addition to evaluation of the annotations used for VQSR. In order to develop these resources we developed a method to evaluate many parameters in parallel and iterating until a final set of values are determined to be acceptable. The general principle is that we brute force test all combinations of parameters and evaluate the number of variants in the resulting tranches at each iteration. At each iteration, we choose an "optimal" set of parameters, recalibrate the SitesOnly files, evaluate Mendelian discordance in trios and select PASS variants to generate a new "truth" set. Here, "optimal" is subjective and is based on examining the behavior of the number of variants in each tranche as well as the clustering of variants in the summary PDF files where more tightly clusters variants separating known/novel, filtered/retained, and positive/negative are indicative of better recalibration. Quantitatively, we count Mendelian errors by tranche in a series of trios where lower error rate indicate better performing models.

<u>Tranches</u>: In order to evaluate the finer scale impact of parameter changes we use 21 total tranches with 10 tranches between 90-99 and 10 tranches between 99.0 and 100 for both SNP and INDEL, which are evaluated independently. PASS variants are set as tranche 90 for evaluation purposes.

<u>Truth sets</u>: The final set of variants used for VQSR after seven iterations are shown in Supplemental Table 6. Each of the resources represent unique variants starting with the most confident variants. The HD_F250 are loci contained on the Illumina Bovine HD (Illumina, San Diego, CA) and Geneseek GGPF250 (Rowan et al. 2019) assays that were filtered on variant call rate and Mendelian error rate metrics from >10,000 samples genotyped on each assay. The remaining SNP truth/training sets were selected based on the criteria below:

```
UMAG1_SNP_TRUTH1U: tranche PASS, 0.05<AF<1.0, SnpGap 10, No trio errors
UMAG1_SNP_TRUTH2U: tranche PASS, 0.01<AF<0.05, SnpGap 10, No trio errors
UMAG1_SNP_TRUTH3U: tranche PASS, 0.001<AF<0.01, SnpGap 10, No trio errors
UMAG1_SNP_train1U: tranche 90.0-98.0, 0.05<AF<1.0, SnpGap 10, No trio errors
UMAG1_SNP_train2U: tranche 90.0-98.0, 0.01<AF<0.05, SnpGap 10, No trio errors
UMAG1_SNP_train3U: tranche 90.0-98.0, 0.001<AF<0.01, SnpGap 10, No trio errors
```

The INDEL truth/training sets used the same criteria except used IndelGap 10.

**Supplemental Table 5. VQSR Truth Set Iterations**

| RESOURCENAME | KNOWN | TRAINING | TRUTH | PRIOR | NUMVARIANTS |
|---|---|---|---|---|---|
| **HD_F250** | false | true | true | 15 | 862,012 |
| AFFYBOS1U | false | true | true | 15 | 316,018 |
| UMAG1_SNP_TRUTH1U | false | true | true | 12 | 17,140,772 |
| UMAG1_SNP_TRUTH2U | false | true | true | 12 | 12,865,155 |
| UMAG1_SNP_TRUTH3U | false | true | true | 12 | 12,303,684 |
| UMAG1_SNP_TRAIN1U | false | true | true | 10 | 4,194,140 |
| UMAG1_SNP_TRAIN2U | false | true | true | 10 | 2,930,294 |
| UMAG1_SNP_TRAIN3U | false | true | true | 10 | 19,590,532 |

| | | | | | |
|---|---|---|---|---|---|
| UMAG1_BQSRV4 | true | false | false | 7 | 174,362,380 |
| **UMAG1_INDEL_TRUTH1U** | false | true | true | 12 | 2,651,881 |
| UMAG1_INDEL_TRUTH2U | false | true | true | 12 | 1,517,732 |
| UMAG1_INDEL_TRUTH3U | false | true | true | 12 | 1,375,626 |
| UMAG1_INDEL_TRAIN1U | false | true | true | 10 | 372,192 |
| UMAG1_INDEL_TRAIN2U | false | true | true | 10 | 319,137 |
| UMAG1_INDEL_TRAIN3U | false | true | true | 10 | 1,251,861 |
| UMAG1_BQSRV4 | true | false | false | 2 | 174,362,380 |

<u>Parameters evaluated:</u> Below is a summary of the parameters and values tested, their defaults and the optimal values. For each of the variables, they were run combinatorically such that only one was varied while holding all others constant to evaluate the impact of each variable in isolation and with respect to others. This results in approximately 135-150 VQSR runs per iteration. After the first couple of rounds of iterations, it was determined that max-gaussians of 9 or 10 and max-negative-gaussians of 5 or 6 performed very poorly and were abandoned in later iterations. As a general observation, we found the bad-lod-score-cutoff to have the most significant impact. As part of the pipeline we evaluate the distribution of the number of variants in the positive and negative model versus the VQSLOD score for each run (**Supplemental Figure 7**). It was noticed early in the testing that the variants included in the negative model produced a long tailed distribution. The variants in the long tail represent the loci that consistently produce very negative LOD scores indicating that they are very poor quality. However, the majority of the density is contained near the bad-lod-score-cutoff and where this parameter is set greatly impacts the ability of the model to differentiate “good” from “bad” variants. Extreme values (more negative) for the bad-lod-score-cutoff result in too few variants to effectively model the variation. Conversely, the default value of -5 includes far too many variants and makes it difficult to differentiate “good” from “bad” variants. All of these variables are interdependent and are also dataset specific which makes obtaining truly optimal values very difficult (**Supplemental Figure 8 – 9**). As stated in the GATK documentation “VQSR is probably the hardest part of the Best Practices to get right.” We have performed similar analyses in dogs, pigs and honeybees and each results in different optimal values.

max-gaussians: 4,5,6,7,8,9,10, [default 8], Optimal SNP=4, INDEL=4
max-negative-gaussians: 2,3,4,5,6 [default 2] , Optimal SNP=4, INDEL=2
minimum-bad-variants: 1000 [default 1000]
maximum-training-variants: 10000000 [default 2500000]
bad-lod-score-cutoff: -5,-7,-9,-11,-13,-15,-17,-19,-21,-23,-25 [default -5] , Optimal SNP=-25, INDEL=-25
SNPS Annotations: '-an QD -an ReadPosRankSum -an FS -an MQ -an SOR -an DP'
INDEL Annotations: '-an QD -an ReadPosRankSum -an FS -an SOR -an DP'

```
java -Djava.io.tmpdir=tmp_${LABEL} -XX:ParallelGCThreads=2 -
Xmx${JMEM}g \
-jar $GATKJAR \
VariantRecalibrator \
-nt $Cpu_Node \
-R $REF \
```

```
-V $SitesOnlyList \
-tranche 100.0 -tranche 99.90 -tranche 99.80 -tranche 99.70 -
tranche 99.60 \
-tranche 99.50 -tranche 99.40 -tranche 99.30 -tranche 99.20 -
tranche 99.10 \
-tranche 99.0 -tranche 98.0 -tranche 97.0 -tranche 96.0 -tranche
95.0 \
-tranche 94.0 -tranche 93.0 -tranche 92.0 -tranche 91.0 -tranche
90.0 \
--mode $MODE \
--seconds-between-progress-updates 300 \
--trust-all-polymorphic \
$Annotations \
-resource:\
${key},known=${Known},training=${Training},truth=${Truth},prior=$
{Prior} $KVPath \
--max-gaussians $MG \
--max-negative-gaussians $MNG \
--minimum-bad-variants $NUMBAD \
--maximum-training-variants $NUMTRAIN \
--max-iterations $MaxIterations \
--max-attempts 5 \
--bad-lod-score-cutoff $BADLOD \
--output ${LABEL}.recal \
--tranches-file ${LABEL}.tranches \
--rscript-file ${LABEL}.R \
--output-model ${LABEL}.model \
--tmp-dir tmp_${LABEL}
```

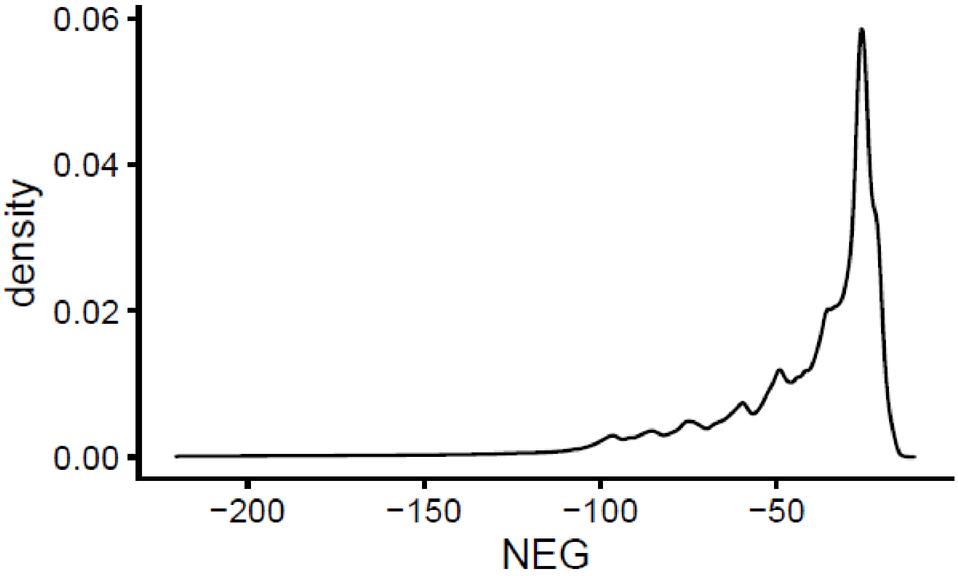

**Supplemental Figure 7**. Example of the distribution of the number of loci in the negative model used for VQSR in the optimal SNP model with a bad-lod-score-cutoff = -25.

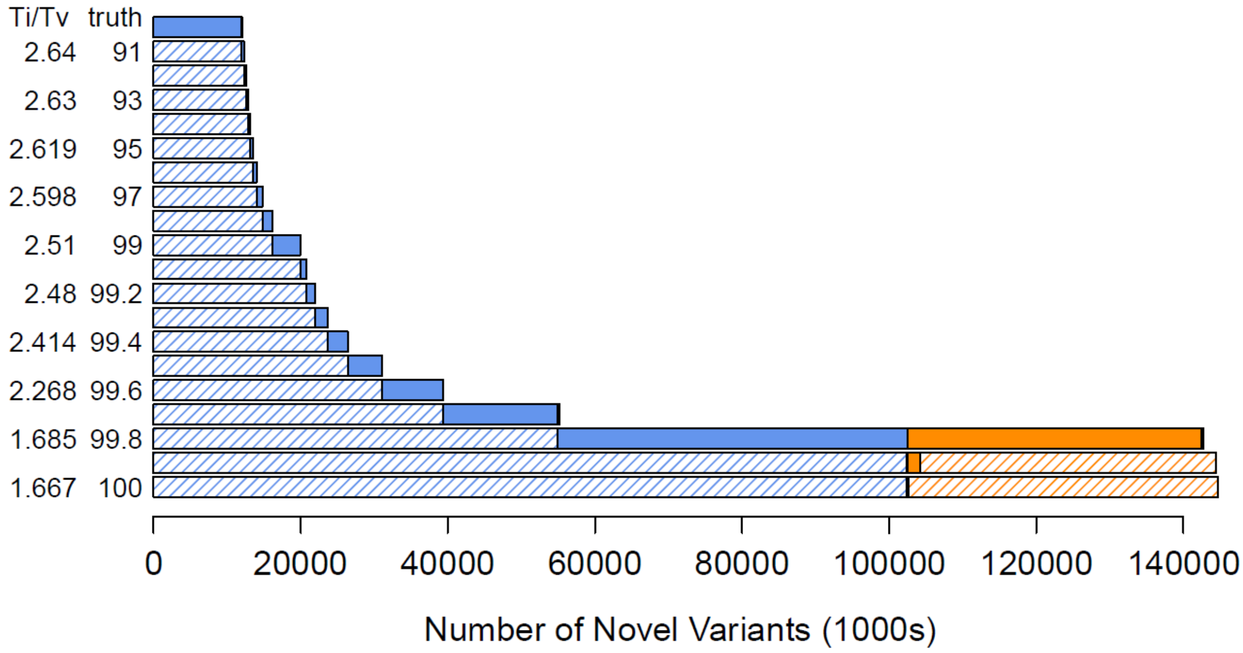

**Supplemental Figure 8.** Tranche plot for the optimal SNP model parameters.

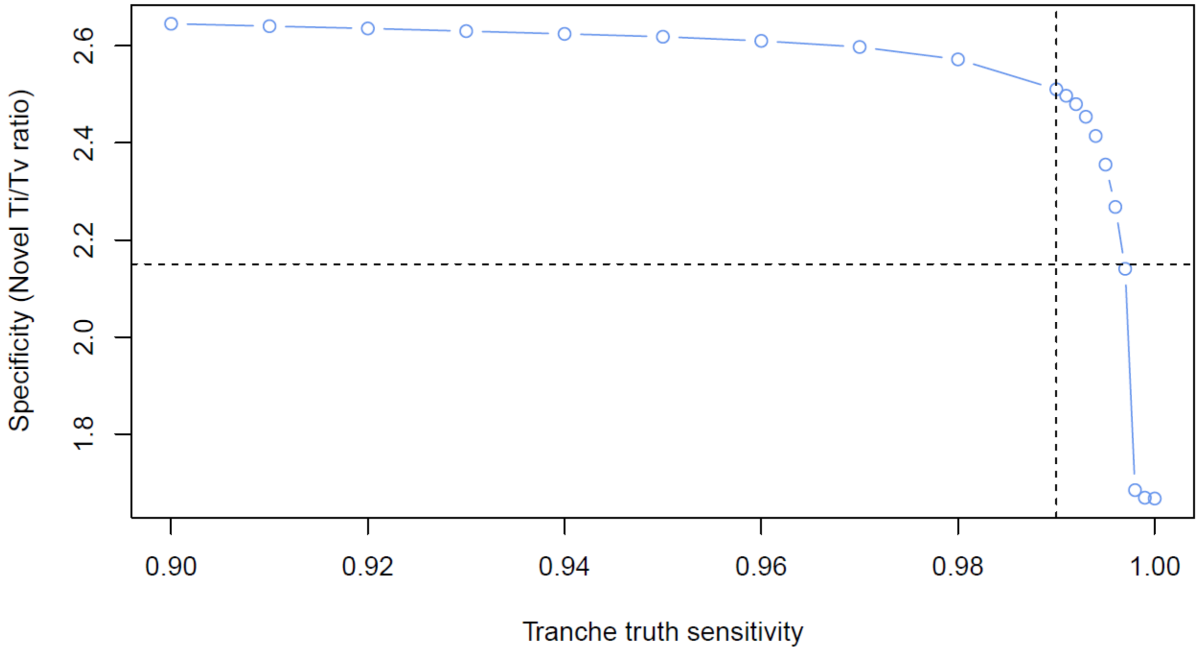

**Supplemental Figure 9.** Tranche sensitivity plot for the optimal SNP model parameters.

**Supplemental Note 15. VQSR Mendelian Error Evaluation:** A total of 15 trios were identified where each member of the trio was sequenced at a minimum of 10x coverage. These trios were used for evaluating the Mendelian Error rate for chromosome 25 for the optimal VQSR parameters to assist with determining the appropriate tranche sensitivity value. For each of the 21 tranches, we calculate the Mendelian error rate after recalibration in the 15 trios for only the loci in the specific tranche. This allows us to evaluate the impact of increasing sensitivity which will increasingly include lower quality variants. **Supplemental Figures 10 and 11** demonstrate an increase in SNP Mendelian error rate as sensitivity increases with a noticeable increase in error rate at tranche 99.7. **Supplemental Figures 12 and 13** show the same trend for indels but with a much higher baseline error rate which is typical for indels with a noticeable increase at tranche 99.8. We also contrast the error rates from our optimized VQSR against the GATK defaults, and the 1kBulls Run9 in **Supplemental Figure 14**.

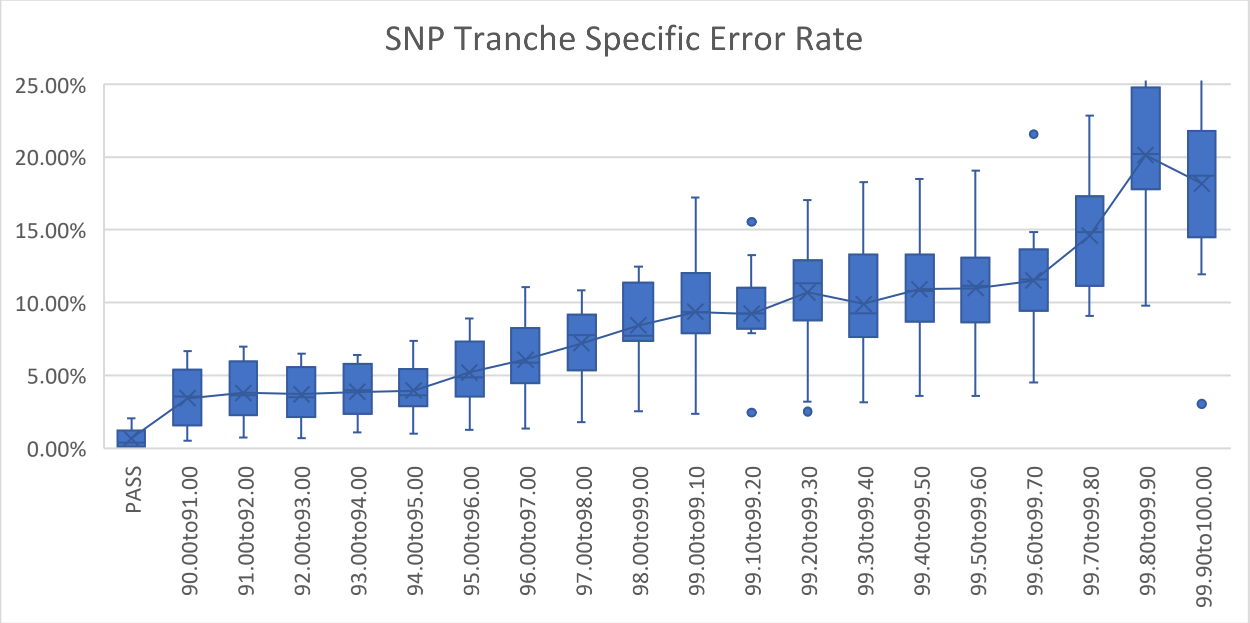

**Supplemental Figure 10.** Tranche specific Mendelian error rate for SNP variants exclusive to each tranche with a noticeable increase in error rate after tranche 99.7.

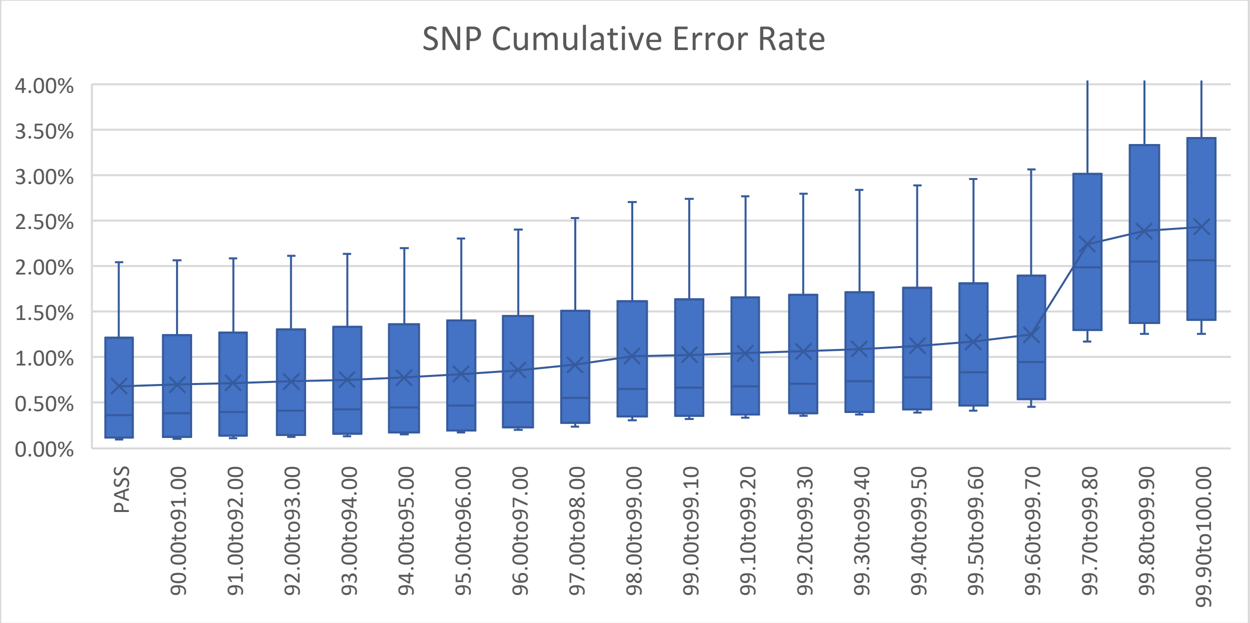

**Supplemental Figure 11.** Cumulative Mendelian error rate by tranche where each tranche includes loci at lower sensitivities. There is a steady increase in error rate up till tranche 99.7 where the remaining 0.3% of variants more than double the error rate. Based on Mendelian error profiles and other factors, a final threshold of tranche 99.7 was chosen to set as PASS.

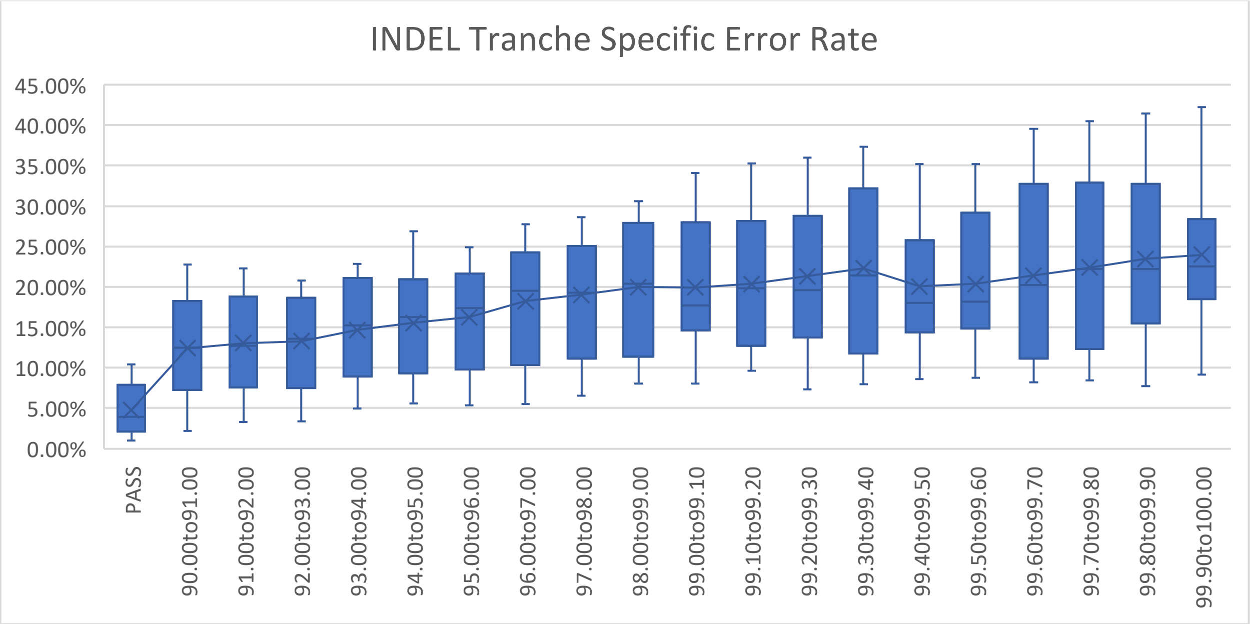

**Supplemental Figure 12.** Tranche specific Mendelian error rate for INDEL variants exclusive to each tranche with a noticeable increase in error rate at tranche 91.

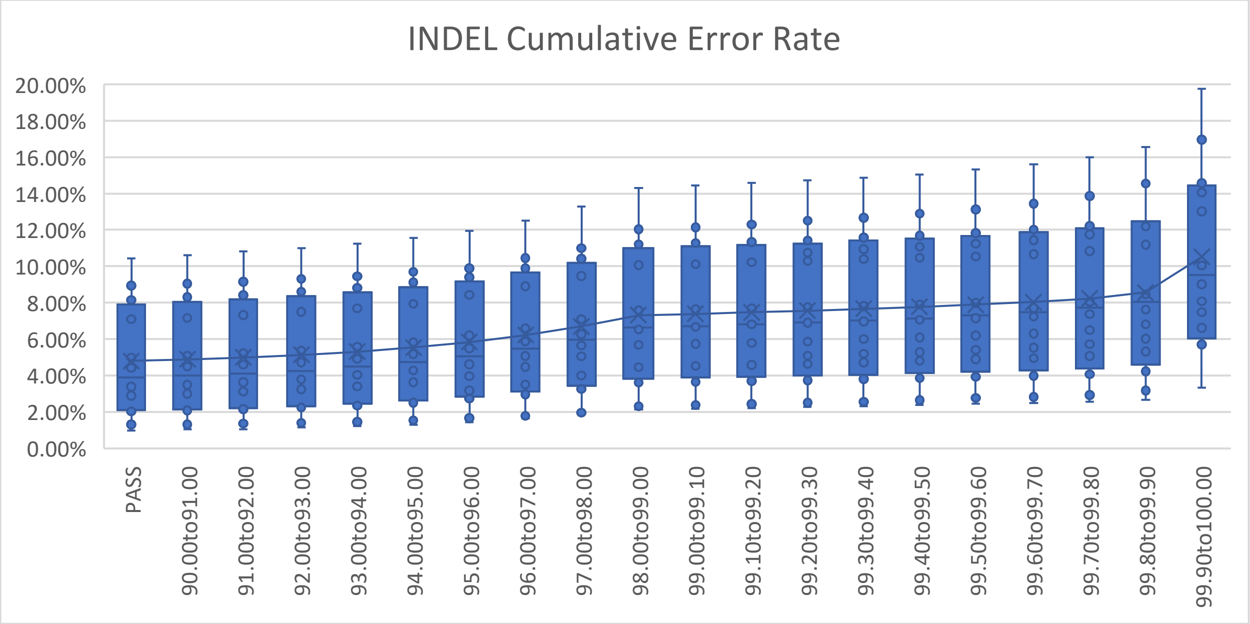

**Supplement Figure 13.** Cumulative Mendelian error rate by tranche where each tranche includes loci at lower sensitivities. There is a steady increase in error rate up till tranche 99.9. Based on Mendelian error profiles and other factors, a final threshold of tranche 99.8 was chosen to set as PASS.

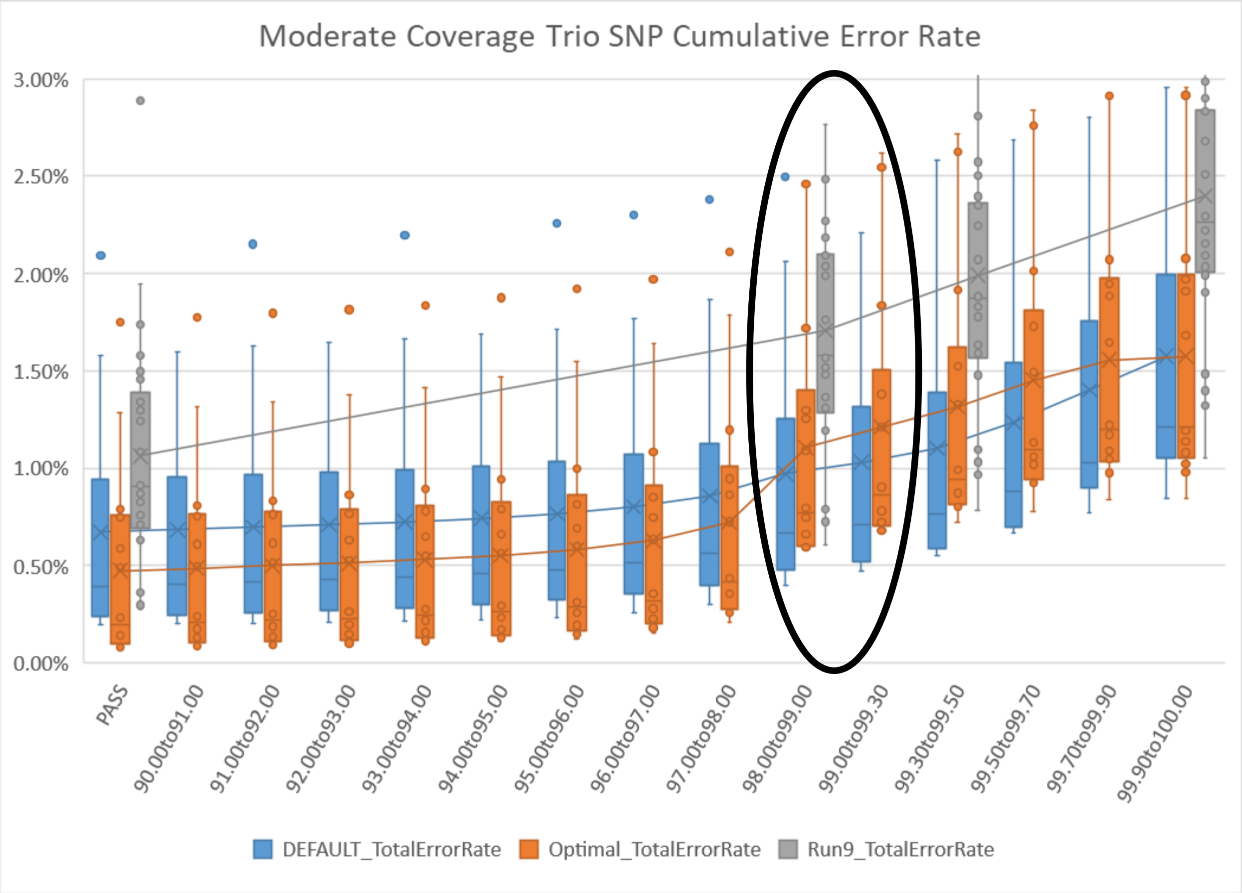

**Supplemental Figure 14. Comparing VQSR approaches.**
Box and whisker plots showing the Mendelian discordance rate for 13 trios using the default (blue), the optimal (orange), and the 1kBulls Run9 (grey) VQSR parameter values. The optimal parameter set indicated a relatively flat and low error rate up to the tranche value of 97 with an elevated error rate after tranche 99 (circled).

**Supplemental Note 16. ApplyVQSR:** After VQSR parameter evaluation, we set PASS variants at SNP tranche sensitivity 99.7 and INDEL sensitivity 99.8 using the respective optimal models for each chromosome in parallel using GATK 4.3.0.0.

```
java -Djava.io.tmpdir=tmp_SNP_${c} -XX:ParallelGCThreads=2 -
Xmx${Jmem}g \
-jar $GatkJar ApplyVQSR -R $Ref \
-V ${YYMMDD}_${COHORT}/vcf/${YYMMDD}.${COHORT}.$c.vcf.gz \
-O ${COHORT}.SNP.${c}.vcf.gz \
-L $c \
--truth-sensitivity-filter-level $SnpSensitivityTranche \
--tranches-file ${OptimalSnpLabel}.tranches \
--recal-file ${OptimalSnpLabel}.recal \
--tmp-dir tmp_SNP_${c} \
--seconds-between-progress-updates 120 \
-mode SNP
```

```
java -Djava.io.tmpdir=tmp_INDEL_${c} -XX:ParallelGCThreads=2 -
Xmx${Jmem}g \
-jar $GatkJar ApplyVQSR -R $Ref \
-V ${COHORT}.SNP.${c}.vcf.gz \
-O ${COHORT}.SNP.INDEL.${c}.vcf.gz \
-L $c \
--truth-sensitivity-filter-level $IndelSensitivityTranche \
--tranches-file ${OptimalIndelLabel}.tranches \
--recal-file ${VqsrDirIndel}/${OptimalIndelLabel}.recal \
--tmp-dir tmp_INDEL_${c} \
--seconds-between-progress-updates 120 \
-mode INDEL
```

**Supplemental Note 17. Initial Quality Control (QC)**
We examined the initial call set using bcftools 1.16 smpl-stats plugin to identify low quality samples to be excluded. As expected, sample coverage was the primary determinant of the number of PASS genotypes per sample (**Supplemental Figure 15**). Samples with a call rate < 0.85 were flagged to be excluded (N=47). Samples with a call rate between 0.85 – 0.90 were retained at this stage because other metrics such as Ts/Tv, number of singletons and heterozygous to homozygous ratio were all within the normal range (**Supplemental Figure 16**).

```
bcftools +smpl-stats -i 'FILTER=\"PASS\"' -o tmp.$c.PASS.tsv
${TaxonID}.UMAG${UmagVersion}.ENSEMBL${AnnotationVersion}.$c.bcf
bcftools view --threads $CPU_view -S ^${SamplesToRemove} -Ou \
$InputDir/${TaxonID}.UMAG${UmagVersion}.ENSEMBL${AnnotationVersio
n}.$c.bcf |\ bcftools --trim-alt-alleles -e 'ALT=\".\" -Ob -o \
${OutputDir}/${TaxonID}.UMAG${UmagVersion}.ENSEMBL${AnnotationVer
sion}.$c.bcf
bcftools index --threads $CPU_index \
${OutputDir}/${TaxonID}.UMAG${UmagVersion}.ENSEMBL${AnnotationVer
sion}.$c.bcf
```

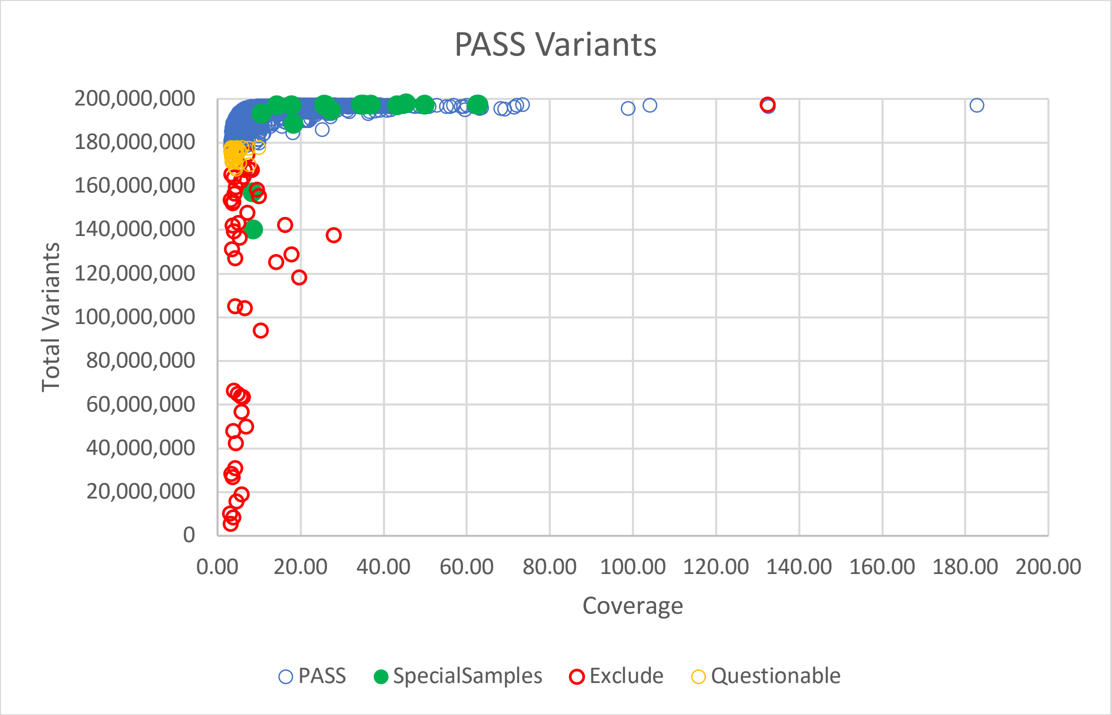

**Supplemental Figure 15.** Initial QC of samples demonstrating the impact of genome coverage. Samples with a total call rate < 0.85 were flagged to be excluded (red). The one excluded outlier at 132x coverage with 99.99% call rate was a sample from SRA which was later discovered to be the same sample used to create the reference genome which was already present using a different identifier. The “special samples” in green represent samples present in trios that were used to generate alternate genome assemblies and three ancient bovid samples.

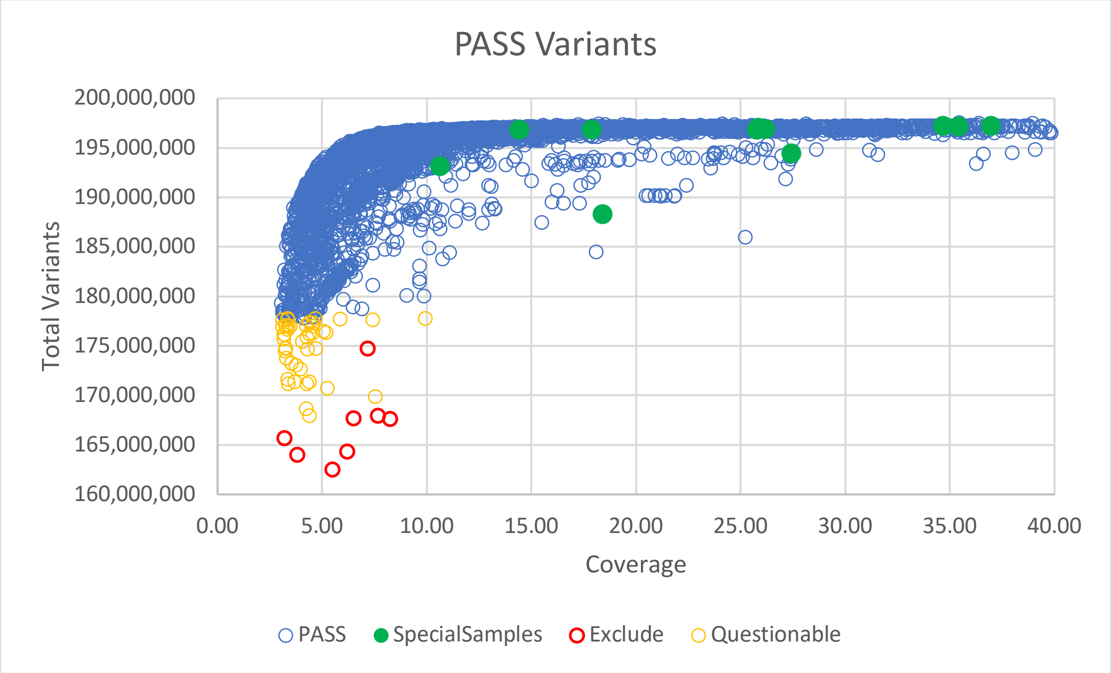

**Supplement Figure 16.** Initial QC of samples with axes truncated. The samples labeled as “questionable” have call rate between 0.85 and 0.90.

**Supplemental Note 18. Training Sample Criteria and Phases**
Cattle genomics data frequently includes families. While multiple offspring can occur per generation in cattle, they typically result via artificial insemination using one sire for several dams rather than multiple full siblings in mosquitos. Publicly available cattle sequencing data includes pedigrees across generations, offering a larger sample size than the human GIAB trios.

Phases 1 and 2: Consists of 18 training iterations (2 parents per trio x 9 cattle trios) across 26 unique genomes. These genomes represent six purebred trios and three crossbred trios. Additionally, five trios had increased similarity to the Hereford-built reference genome, while the remaining four trios represent more divergent breeds relative to the reference. These samples were randomized to identify any trio-specific characteristics that enable gains in training performance.

Phase 3: Consists of 4 training iterations (2 parents per trio x 2 Bison trios) across five unique genomes, as the offspring shared a sire. Bison samples were included to explore the impact of outgroup species upon training.

Phase 4: Consists of 6 training iterations (2 parents per trio x 3 real F1-hybrid trios) across nine unique genomes. The training was extended for these samples to determine if higher-quality truth labels could meaningfully impact performance.

Phase 5: Consists of 2 training iterations (2 parents per trio x 1 synthetic F1-hybrid trios) across one unique genome – a SynDip replicate of the Angus-Brahman trio from Phase 4 using the real, short-read WGS for the parents, but synthetic reads for the offspring. A single synthetic trio was used to reserve the remaining two trios for model testing. This phase assessed if synthetic truth labels could be substituted for GIAB-quality truth labels.

**Supplemental Table 6. Training Phases**Re-training DeepVariant for bovine genomes is achieved over 30 iterations, representing 15 family trios. These training iterations are subset into five phases for summary comparisons. The terminal checkpoint for each phase is from the second iteration of the last trio within a phase

| **Trio** | | | **Pedigree** | | | | **Mean Coverage** | | |
|---|---|---|---|---|---|---|---|---|---|
| *Breed* | *Name* | *Discordance* | *Offspring Lab ID* | *Sex* | *Father Lab ID* | *Mother Lab ID* | *Offspring* | *Father* | *Mother* |
| ***Phase 1*** | | | | | | | | | |
| *Angus (AA)* | Trio1 | 0.1054% | 118766 | F | 32157 | 118765 | 23.64 | 35.96 | 23.10 |
| *Hereford (HE)* | Trio2 | 0.1653% | 199724 | F | 199730 | 199727 | 14.45 | 17.47 | 12.92 |
| *Brown Swiss (BS)* | Trio3 | 0.1754% | 342161 | U | 342163 | 342162 | 19.10 | 18.00 | 17.30 |
| *Holstein (HO)* | Trio4 | 0.3715% | 342485 | M | 341441 | 341281 | 23.66 | 35.23 | 46.68 |
| *Hereford (HE)* | Trio5 | 0.1754% | 199723 | F | 199731 | 199728 | 15.89 | 13.51 | 15.08 |
| *Tyrolean Grey (TG)* | Trio6 | 0.1007% | 342304 | U | 342303 | 342305 | 24.90 | 25.30 | 24.10 |
| ***Phase 2*** | | | | | | | | | |
| *Holstein-Jersey (HJ)* | Trio7 | 0.1412% | 342472 | F | 341035 | 340965 | 50.83 | 39.35 | 44.75 |
| *Holstein-Jersey (HJ)* | Trio8 | 0.3822% | 341101 | M | 340998 | 342473 | 27.65 | 30.81 | 39.31 |
| *Holstein-Hereford (HH)* | Trio9 | 0.4427% | 199725 | M | 199730 | 199726 | 14.79 | 17.47 | 14.62 |
| ***Phase 3*** | | | | | | | | | |
| *Bison (BI)* | Trio10 | 0.2725% | 20136 | F | 2406 | 20076 | 17.95 | 22.69 | 23.72 |
| *Bison (BI)* | Trio11 | 0.5329% | 20172 | M | 2406 | 20098 | 24.06 | 22.69 | 23.78 |
| ***Phase 4*** | | | | | | | | | |
| *Angus/Brahman F1 (AA/BR)* | Trio12 | 0.6368% | 341496 | M | 194551 | 194550 | 50.01 | 36.99 | 43.32 |
| *Yak/Highlander F1 (YK/HI)* | Trio13 | 0.4829% | 341497 | F | 204543 | 204544 | 17.92 | 34.69 | 10.66 |
| *Bison/Simmental F1 (BI/SI)* | Trio14 | 1.0966% | 341713 | M | 341714 | 339207 | 14.46 | 27.44 | 35.42 |
| ***Phase 5*** | | | | | | | | | |
| *Synthetic AA/BR F1* | Trio15 | 0.6797% | 9341496 | M | 194551 | 194550 | 26.20 | 36.99 | 43.32 |

**Supplemental Table 7. Training Checkpoints** The table below provides the number of training steps covered at the selected best checkpoint, where F1 score in the offspring genome is maximized. We also describe the number of bovine training examples used to re-train the DeepVariant model to achieve the best checkpoint. In total, we provided the model with nearly 400M new example variants across the entire training period, but only 230M were used to generate the optimal checkpoints across all 30 iterations.

| Trio | | | | Checkpoint | | Examples | | |
|---|---|---|---|---|---|---|---|---|
| *Breed* | *Name* | *Iteration* | *Training Genome* | *Steps Covered* | *Loss* | *Used* | *Created* | *%* |
| | | | ***Phase 1*** | | | | | |
| *Angus (AA)* | *Trio1* | *1* | *Sire* | 84,518 | 0.05812 | 2,704,576 | 8,273,689 | 33% |
| | | *2* | *Dam* | 201,011 | 0.04866 | 6,432,352 | 8,638,474 | 74% |
| *Hereford (HE)* | *Trio2* | *3* | *Sire* | 134,175 | 0.09964 | 4,293,600 | 7,252,806 | 59% |
| | | *4* | *Dam* | 77,316 | 0.10315 | 2,474,112 | 7,052,315 | 35% |
| *Brown Swiss (BS)* | *Trio3* | *5* | *Sire* | 76,110 | 0.07617 | 2,435,520 | 8,557,378 | 28% |
| | | *6* | *Dam* | 99,664 | 0.07889 | 3,189,248 | 8,521,779 | 37% |
| *Holstein (HO)* | *Trio4* | *7* | *Sire* | 15,507 | 0.10058 | 496,224 | 8,081,303 | 6% |
| | | *8* | *Dam* | 113,822 | 0.07635 | 3,642,304 | 8,309,599 | 44% |
| *Hereford (HE)* | *Trio5* | *9* | *Sire* | 127,591 | 0.09844 | 4,082,912 | 6,692,877 | 61% |
| | | *10* | *Dam* | 63,019 | 0.09767 | 2,016,608 | 7,183,727 | 28% |
| *Tyrolean Grey (TG)* | *Trio6* | *11* | *Sire* | 159,745 | 0.04826 | 5,111,840 | 8,701,670 | 59% |
| | | *12* | *Dam* | 203,095 | 0.04837 | 6,499,040 | 8,799,190 | 74% |
| | | | ***Phase 2*** | | | | | |
| *Holstein-Jersey (HJ)* | *Trio7* | *13* | *Sire* | 101,101 | 0.06093 | 3,235,232 | 8,356,023 | 39% |
| | | *14* | *Dam* | 120,966 | 0.05960 | 3,870,912 | 8,646,939 | 45% |
| *Holstein-Jersey (HJ)* | *Trio8* | *15* | *Sire* | 225,421 | 0.06192 | 7,213,472 | 8,268,746 | 87% |
| | | *16* | *Dam* | 116,248 | 0.06444 | 3,719,936 | 8,131,368 | 46% |
| *Holstein-Hereford (HH)* | *Trio9* | *17* | *Sire* | 77,464 | 0.10359 | 2,478,848 | 7,252,806 | 34% |
| | | *18* | *Dam* | 120,844 | 0.10119 | 3,867,008 | 7,082,305 | 55% |
| | | | ***Phase 3*** | | | | | |
| *Bison (BI)* | *Trio10* | *19* | *Sire* | 228,738 | 0.06509 | 7,319,616 | 28,833,745 | 25% |
| | | *20* | *Dam* | 892,234 | 0.05785 | 28,551,488 | 30,638,318 | 93% |
| *Bison (BI)* | *Trio11* | *21* | *Sire* | 803,481 | 0.06344 | 25,711,392 | 28,833,044 | 89% |
| | | *22* | *Dam* | 935,368 | 0.05978 | 29,931,776 | 30,540,725 | 98% |

| | | | | | | | | |
|---|---|---|---|---|---|---|---|---|
| | | | | ***Phase 4*** | | | | |
| *Angus/Brahman F1 (AA/BR)* | *Trio12* | *23* | *Sire* | 59,354 | 0.10298 | 1,899,328 | 9,353,296 | 20% |
| | | *24* | *Dam* | 217,570 | 0.11639 | 6,962,240 | 19,640,662 | 35% |
| *Yak/Highlander F1 (YK/HI)* | *Trio13* | *25* | *Sire* | 275,488 | 0.08869 | 8,815,616 | 8,815,636 | 100% |
| | | *26* | *Dam* | 432,983 | 0.08369 | 13,855,456 | 29,085,226 | 48% |
| *Bison/Simmental F1 (BI/SI)* | *Trio14* | *27* | *Sire* | 204,026 | 0.12034 | 6,528,832 | 30,506,602 | 21% |
| | | *28* | *Dam* | 282,383 | 0.12304 | 9,036,256 | 9,036,262 | 100% |
| | | | | ***Phase 5*** | | | | |
| *Synthetic AA/BR F1* | *Trio15* | *29* | *Sire* | 290,765 | 0.17478 | 9,304,480 | 9,353,296 | 99% |
| | | *30* | *Dam* | 336,710 | 0.10775 | 10,774,720 | 19,640,662 | 55% |
| **TOTAL** | | | | | | 226,454,944 | 398,080,468 | |

**Supplemental Note 19. Testing Samples Used Per Phase**

Phases 1 – 3: Consists of the same 19 samples across all three phases. While training was performed for these phases, DeepVariant has never seen any labeled examples from the six F1-hybrid offspring, either real or synthetic. Thus, we used the 13 unrelated samples and the 6 F1-hybrid offspring to assess performance and track generalizability over the first 22 iterations.

Phases 4 and 5: Consists of the 13 unrelated samples while incrementally removing one of the F1-hybrid samples once the first iteration begins. While training was performed for these phases, as soon as DeepVariant saw labels from an F1-hybrid trio, the offspring was never used for model testing again. By eliminating these samples from our testing cohort iteratively, we minimized data bleed because samples are never used for model selection and performance testing.

**Supplemental Table 8. Testing Phases.** Each model iteration is tested using an independent set of previously unseen bovine genomes. After Phase 3, we began to extend training into the F1-hybrid trios. As such, the set of testing genomes decreases by one as training proceeds to another trio, dropping the F1-hybrid offspring from further use as a test genome.

| | | | | | *Phases 1 - 3* | | | *Phase 4* | | | | | | *Phase 5* | |
|---|---|---|---|---|---|---|---|---|---|---|---|---|---|---|---|
| | **Test Genome** | | | | **Iterations Used** | | | | | | | | | | |
| *Breed* | *Name* | *Lab ID* | *Sex* | *Mean Coverage* | *All* | *All* | *All* | *23* | *24* | *25* | *26* | *27* | *28* | *29* | *30* |
| *Angus* | Test1 | 186 | M | 31.4 | x | x | x | x | x | x | x | x | x | x | x |
| *Shorthorn* | Test2 | 71941 | M | 25.7 | x | x | x | x | x | x | x | x | x | x | x |
| *Charolais* | Test3 | 20809 | M | 20.1 | x | x | x | x | x | x | x | x | x | x | x |
| *Gelbvieh* | Test4 | 34095 | M | 20.0 | x | x | x | x | x | x | x | x | x | x | x |
| *Hereford* | Test5 | 33604 | F | 45.5 | x | x | x | x | x | x | x | x | x | x | x |
| *Jersey* | Test6 | 185699 | M | 24.1 | x | x | x | x | x | x | x | x | x | x | x |
| *Limousin* | Test7 | 18336 | M | 20.7 | x | x | x | x | x | x | x | x | x | x | x |
| *MaineAnjou* | Test8 | 87957 | M | 22.2 | x | x | x | x | x | x | x | x | x | x | x |
| *Salers* | Test9 | 51124 | M | 24.9 | x | x | x | x | x | x | x | x | x | x | x |
| *Simmental* | Test10 | 71657 | M | 24.5 | x | x | x | x | x | x | x | x | x | x | x |
| *Holstein* | Test11 | 196818 | M | 52.8 | x | x | x | x | x | x | x | x | x | x | x |
| *Chianina* | Test12 | 194426 | U | 13.5 | x | x | x | x | x | x | x | x | x | x | x |
| *BrownSwiss* | Test13 | 204654 | U | 60.0 | x | x | x | x | x | x | x | x | x | x | x |
| *Angus/Brahman F1 (AA/BR)* | Test14 | 341496 | M | 50.0 | x | x | x | | | | | | | | |
| *Yak/Highlander F1 (YK/HI)* | Test15 | 341497 | F | 17.9 | x | x | x | x | x | | | | | | |
| *Bison/Simmental F1 (BI/SI)* | Test16 | 341713 | M | 14.5 | x | x | x | x | x | x | x | | | | |
| *Synthetic AA/BR F1* | Test17 | 9341496 | M | 26.20 | x | x | x | x | x | x | x | x | x | | |
| *Synthetic YK/HI F1* | Test18 | 9341497 | F | 25.79 | x | x | x | x | x | x | x | x | x | x | x |
| *Synthetic BI/SI F1* | Test19 | 9341713 | M | 25.85 | x | x | x | x | x | x | x | x | x | x | x |

**Supplemental Note 20. Creating Truth Sets for DeepVariant**
After cohort QC, we generated truth sets based on the UMAG1 cohort using GATK derived genotypes. The regions files produced by GATK (3.8-1-0-gf15c1c3ef) CallableLoci were parsed by sample to extract only PASS regions that were used for downstream analyses. We first extract trio and testing samples from the full BCF (all variants) and write to 17 trio specific files, plus a file containing the testing individuals using bcftools 1.14. This is performed per chromosome and then individual chromosomes are concatenated and indexed to produce a single BCF file per trio.

```
bcftools +split --groups-file Group.txt -Ob -o $c
9913.UMAG1.ENSEMBL106.$c.bcf
bcftools concat --file-list Trio${t}.merge.list -Ob -o
RAW.Trio${t}.bcf
bcftools index RAW.Trio${t}.bcf
```

Mendelian errors in the raw BCF file are indicative of poorly genotypes positions within one or more of the members of a trio. We removed Mendelian errors using bcftools 1.14 to produce trio specific BCF files with no errors

```
bcftools +mendelian --rules-file $RulesFile RAW.Trio${t}.bcf \
-Ob -o CLEAN.RAW.Trio${t}.bcf -m d --trio-file Trio${t}.ID.txt
bcftools index CLEAN.RAW.Trio${t}.bcf
```

A third set of files are created using bcftools 1.14 on an individual basis that are used for training. For each sample, we only extract variants that are contained within callable regions for that sample and we exclude all sites where the individual is homozygous for the reference allele, has a missing genotype or contains a spanning deletion (*). Note that the previous removal of Mendelian errors sets the genotype at the site to missing and these sites will be excluded at this step. During initial development, we also extracted a series of files that also restricted the genotypes based on genotype quality (GQ=10,13,20,30) to evaluate the impact of removing lower quality genotypes on training.

```
bcftools view --samples $sample --regions-file
$sample.callable.bed -Ou CLEAN.RAW.Trio${t}.bcf | \
bcftools view --exclude 'GT=\"RR\" | GT=\"mis\" | ALT=\"*\"' -Oz
-o $sample.vcf.gz
bcftools index --tbi $sample.vcf.gz
```

A population VCF was created containing all PASS variants to use the `--use_allele_frequency` option of DeepVariant. The allele frequencies are obtained from the cohort call set based on 5,512 samples representing our best estimate of population allele frequencies. The resulting file contained 141,595,864 SNP and 18,246,855 INDEL.

```
bcftools view -G -f PASS --exclude 'ALT=\"*\"' -Oz -o
POP.$i.vcf.gz \ 9913.UMAG1.ENSEMBL106.$i.bcf
bcftools concat --file-list POP.merge.list -Oz -o
UMAG1.POP.FREQ.vcf.gz
bcftools index --tbi UMAG1.POP.FREQ.vcf.gz
```

**Supplemental Note 21. Synthetic Diploid Reads**

Three phased genome assemblies based on offspring from divergent parents were used to create synthetic reads. In each of these three trios, the sex of the F1 that was sequenced dictated the sex chromosome(s) that could be assembled (Rice et al. 2020; Koren et al. 2018; Low et al. 2020; Heaton et al. 2021; Oppenheimer et al. 2021). To have a full complement of the autosomes, sex chromosomes and mitochondrial genome, when a chromosome was missing from a haploid assembly we added the missing chromosome from another assembly (**Supplemental Table 9**). For each of the six haploid genomes, we generated synthetic reads using NEAT 3.2 (Stephens et al. 2016). The parameters used were: -R 150, -c 15, -E 0, -M 0, --pe 350 70, --rng 1234567. These simulate 15x coverage of each haplotype without error, similar to standard Illumina paired end data. However, the per-base error rate of the assemblies was higher than in the raw SRS data, meaning our sampled synthetic reads are noisy (**Supplemental Table 10**). The reads from each chromosome are then concatenated into a single pair of files representing a diploid genome. The reads are then shuffled to remove the linear genomic order that was used to generate them to avoid issues during the alignment stage.

```
python3 gen_reads.py -r $RefName.$c.fa -R $ReadLen -c $COV -E
$ErrorRate \
-M $MutationRate --pe $Pe --rng $RNG --bam -o
OUT/${OUTPREFIX}.${COV}.$c
cat OUT/${RefName}.${COV}.*_read1.fq.gz
>OUT/${RefName}.${COV}.1.fastq.gz
cat OUT/${RefName}.${COV}.*_read2.fq.gz
>OUT/${RefName}.${COV}.2.fastq.gz
File1=OUT/${RefName}.${COV}.1.fastq.gz
File2=OUT/${RefName}.${COV}.2.fastq.gz
Rand1="OUT/RAND/${RefName}.${COV}.1.fastq"
Rand2="OUT/RAND/${RefName}.${COV}.2.fastq"
paste <(zcat $File1) <(zcat $File2) | paste - - - - | shuf | awk
-F'\t' \
-v r1=$Rand1 -v r2=$Rand2 '{OFS="\n"; print $1,$3,$5,$7 > r1; \
print $2,$4,$6,$8 > r2}'
```

**Supplemental Table 9**. Three genome assemblies used to generate synthetic reads. The short name represents the two breeds or species used to make the sequenced offspring and the origin of the sex chromosomes. If a sex chromosome was not assembled, we used the indicated sex chromosome from a different assembly to represent a full genome for each haploid reference. Note that the Hereford ChrY is the same as that used for the ARS-UCD1.2_Btau5.0.1Y reference.

| Short Name | Autosomes | ChrX | ChrY | MT |
|---|---|---|---|---|
| Highland | GCA_009493655.1 | GCA_009493655.1 | GCA_000003205.6 | GCA_002263795.2 |
| Paternal | Highland | Highland | Hereford | Hereford |
| Yak | GCA_009493645.1 | GCA_009493645.1 | GCA_000003205.6 | GCA_009493645.1 |
| Maternal | Yak | Yak | Hereford | Yak |
| Angus | GCA_003369685.2 | GCA_002263795.2 | GCA_003369685.2 | GCA_002263795.2 |
| Paternal | Angus | Hereford | Angus | Hereford |
| Brahman | GCA_003369695.2 | GCA_003369695.2 | GCA_000003205.6 | GCA_003369695.2 |
| Maternal | Brahman | Brahman | Hereford | Brahman |
| Simmental | GCA_018282465.1 | GCA_018282465.1 | GCA_000003205.6 | GCA_018282465.1 |
| Maternal | Simmental | Simmental | Hereford | Simmental |
| Bison | GCA_018282365.1 | GCA_002263795.2 | GCA_018282365.1 | NC_012346.1 |
| Paternal | Bison | Hereford | Bison | Bison |

**Supplemental Table 10. Assembly per-base error comparison.** We estimated the per-base error rates for the current bovine reference genome and the assemblies from F1-hybrid crosses used to produce synthetic diploid (SynDip) training data. The expected number of errors is contrasted against the Mendelian Inheritance Error counts in the trio VCFs with either the Synthetic or Real SRS data for the offspring. By sampling from the assemblies, the synthetic reads have an order of magnitude more base errors than the real data used during phase 4 of re-training. As assembly quality improves, we recommend evaluating these metrics before using the SynDip approach to create training data.

| *Assembly Name* | *Source* | *Date Available* | *Base QV* | *Error Probability* | *Genome Size* | **Error Count** | | |
|---|---|---|---|---|---|---|---|---|
| | | | | | | *Expected* | *Synthetic* | *Real* |
| *ARS-UCD1.2* | link | 3/19/20 | 48.67 | 1.36E-05 | 2.75915E+09 | 37,478 | - | - |
| *UOA_Angus_1* | link | 4/29/20 | 44.63 | 3.44E-05 | 2.60000E+09 | 89,531 | 45,157 | 3,370 |
| *UOA_Brahman_1* | link | 4/29/20 | 46.38 | 2.30E-05 | 2.70000E+09 | 62,139 | | |
| *Highlander/Yak* | link | 4/3/20 | ** | ** | ** | - | 23,203 | 6,808 |
| *ARS-UCSC_bison1.0* | link | 2/17/21 | 38.88 | 1.29E-04 | 2.83000E+09 | 366,257 | 52,631 | 8,338 |
| *ARS_Simm_1.0* | link | 1/13/21 | 35.01 | 0.0003155 | 2.86172E+09 | 902,874 | | |

*** Unpublished due to technical issues with error estimation*

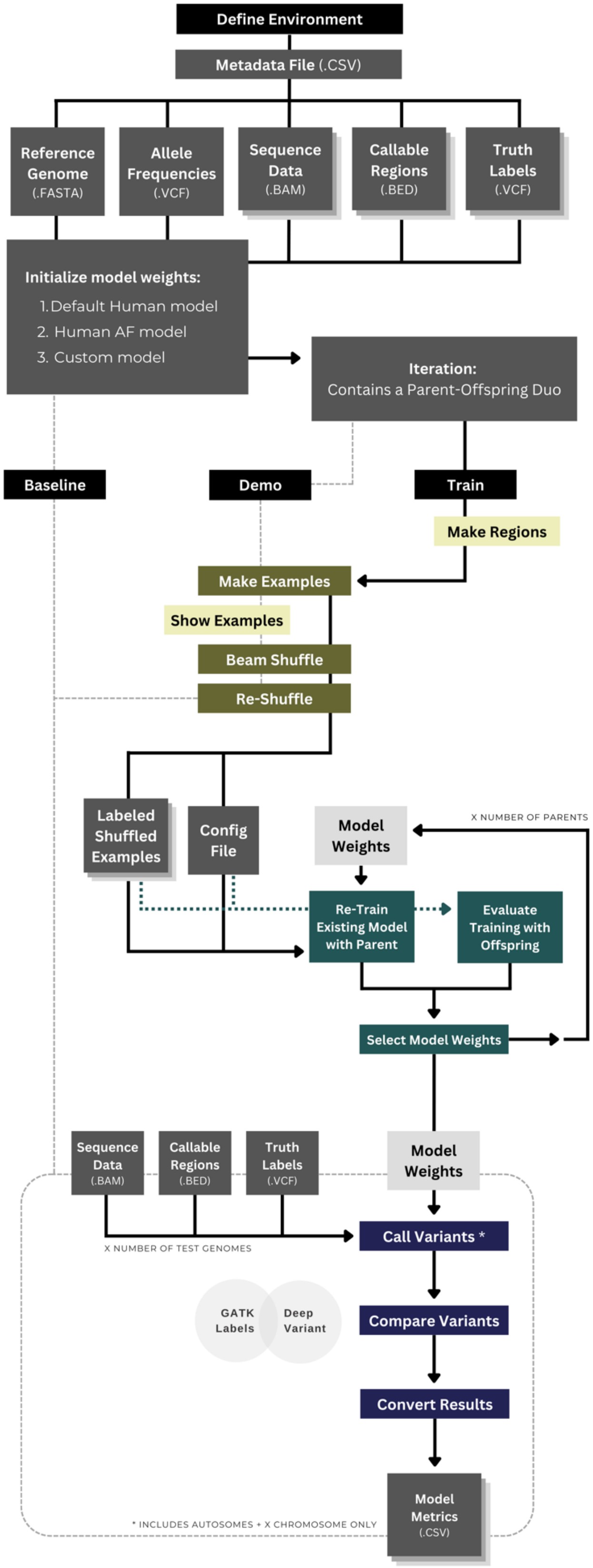
Define Environment
Metadata File (.CSV)
Reference Genome (.FASTA)
Allele Frequencies (.VCF)
Sequence Data (.BAM)
Callable Regions (.BED)
Truth Labels (.VCF)
Initialize model weights:
1. Default Human model
2. Human AF model
3. Custom model
Iteration:
Contains a Parent-Offspring Duo
Baseline
Demo
Train
Make Regions
Make Examples
Show Examples
Beam Shuffle
Re-Shuffle
Labeled Shuffled Examples
Config File
Model Weights
X NUMBER OF PARENTS
Re-Train Existing Model with Parent
Evaluate Training with Offspring
Select Model Weights
Sequence Data (.BAM)
Callable Regions (.BED)
Truth Labels (.VCF)
Model Weights
Call Variants *
X NUMBER OF TEST GENOMES
GATK Labels
Deep Variant
Compare Variants
Convert Results
Model Metrics (.CSV)
* INCLUDES AUTOSOMES + X CHROMOSOME ONLY

**Supplemental Figure 17. Workflow diagram for the TrioTrain pipeline.**
(previous page) Two iterations are performed for each trio: one for each parent. Both iterations use replicates of the same offspring genome for selecting model weights that maximize the F1-score. A training iteration consists of three stages:

(1) *Data Preparation: (yellow) Enables genome-wide shuffling with SLURM.*

Make Regions: Each genome is divided into a set of regions. By default, a region should produce 200,000 or fewer examples so that shuffling fits within memory more effortlessly.

Make Examples: For each region, genome-wide labeled examples are created.

Beam Shuffle: For each region, shuffling of genome-wide occurs in parallel.

Re-Shuffle: The per-CPU shards are merged into per-region tfrecord files; the per-region file order is randomized, and then the paths are written to a new config file.

*(2) Iterative Re-Training: (green) Starts with an existing model checkpoint to sequentially re-train DeepVariant across all trios.*

Train: Shuffled, labeled, merged, and randomized tfrecord files from the parent genome are incrementally provided to the model in groups set by batch size (32) while adjusting model weights using a learning rate (0.005).

Evaluate: As new model weights are produced, genetic variants are called using shuffled, labeled, merged, and randomized examples produced from the Child's genome. Tune the model to accurate genotype inherited variants.

Select: Each checkpoint's performance metrics are stratified by variant type and class. Optimal performance is based on evaluation with the F1-score to select one checkpoint per iteration.

(3) *Testing A Model: (blue) Assess how well the model performs in genomes that the model has not previously seen.*

Call Variants: Use each selected checkpoint for variant calling in genomes outside the trios.

Compare Variants: Use hap.py to assess the new model's variants against a Truth VCF.

Convert Results: Process pattern combinations from hap.py into raw metric counts.

**Supplemental Note 21. Required Inputs for TrioTrain**

Once a starting point is selected at the warm-starting weights, the TrioTrain pipeline automatically prepares the labeled, shuffled examples of each genome in a trio based on a pipeline configuration input file. This metadata file (.csv) contains descriptive information such as mean coverage and the absolute paths to all required input files. Training order is dictated by the row order within the metadata file, while the first parent is set to either “Mother” or “Father” via the --first-parent flag using TrioTrain.

The metatdata file is provided as a command line flag and contains the pedigree and the local file structure for all the input files necessary for re-training DeepVariant. An example of the metadata file can be viewed here:
https://docs.google.com/spreadsheets/d/1hSWD0a5PrWXTQKW4-Q6MTqm-kuI8Bs8cx3kB8u0GEOs/edit?usp=sharing

When using TrioTrain, the following assumptions are made: (1) Each row corresponds to one complete family trio to use in two (2) TrioTrain iterations. (2) Row order determines the sequential order of how data is used during training. However, the number corresponding to a specific test genome (e.g., Test3) does not correspond to the testing order, as testing is performed in parallel.

At a minimum, the pipeline expects the metadata file to have two rows and 24 columns. The minimum rows include a header row, where the second row represents a single trio for re-training. Additional rows correspond to other trios. The first two columns define the order in which data will be given to the model and provide a unique name for each iteration. The following seven columns describe the trio used by providing the sample ID and, optionally, a separate lab ID for the child, father, and mother, as well as the offspring's sex. The following three columns provide the reference genome, the optional population allele frequency VCF file, and the optional regions BED file. The last nine columns provide the input files for the child, father, and mother by providing the absolute paths to three files per genome (BAM/TruthVCF/BED). The last three required columns contain the input files required for each test genome, with three files per test genome (BAM/TruthVCF/BED). Further tests can be achieved by adding three columns for each additional genome. Complete details about the required data and their formats are described further in the TrioTrain user guide: https://jkalleberg.github.io/DV-TrioTrain/user-guide/usage_guide/#assumptions

**Supplemental Note 22. Implementing training on a SLURM-based cluster**

The DeepVariant software has two components: one for genotyping samples and the other for producing new models. While the variant caller is packaged as a single command executed from a Docker container, the training components are not. Additionally, some training steps are incompatible with SLURM-based HPC clusters. Considering these hurdles, we developed TrioTrain for species-agnostic model extension with new data. We automated repeated training rounds from a single command on research computing hardware by re-designing a previously intractable step, the shuffling approach for randomizing genomic order (**Supplemental Figure 18**). Each candidate variant results in a multi-channel tensor record file, known as examples. Channels represent features commonly found in pileup alignment images. The primary difference for training is the addition of an extra channel containing the label (truth genotype). Examples are created in linear genomic order and stored on disk. The total number of examples produced depends on genome divergence from the reference. For example, a typical bovine genome can produce 8–10 million examples, while Bison genomes produce around 30 million (**Supplemental Table 7**). Shuffling is required because training assumes the current example is independent of prior examples. However, achieving randomization of several million images is not trivial.

Our biggest hurdle was ensuring that the labeled examples represented a complete genome yet fit within memory. TrioTrain implements a SLURM-compatible approach which distributes a genome into buckets, called regions, use to handle millions of images created per genome. Before generating examples, we calculate a genome-specific number of regions based on the total truth variants. Each region is defined in a 0-based BED file, where sequential regions overlap by 1b. While the number of genome-wide examples can vary significantly, a region samples the chromosomes proportionally yet produces approximately 200,000 examples. These BED files are used when generating examples, resulting in distributed, independent SLURM jobs for each region.

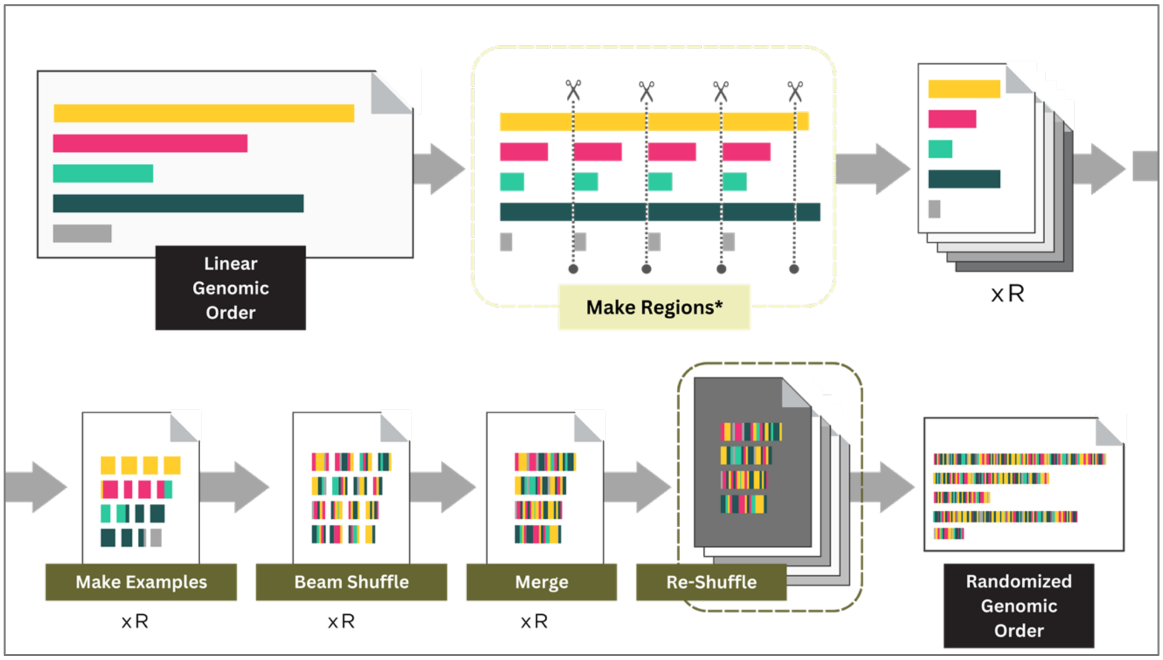

**Supplemental Figure 18. Region Shuffling Approach.** TrioTrain automatically determines how many regions (R) are required to split a complete genome. This calculation is based on a user-defined maximum number of examples per region (default = 200,000), the expected number of examples created per variant, which varies between species (default=1.5), and the number of variants in the corresponding truth VCF. TrioTrain first create R overlapping BED files containing a proportional sample from the autosomes and the X chromosome. Next, within-region labeled examples are made in parallel. To minimize linear genomic order, each genome-wide set of files undergoes shuffling with Apache Beam. The shuffled within-region labels are then concatenated. Finally, the region order is randomized, reducing the chances of providing the model regions sequentially. The respective config file for the re-shuffled examples is given to the model for either training or evaluation, depending on the genome.

**Supplemental Note 23. Defining regions for shuffling**

We calculate the number of regions per genome, RG, as follows. First, TrioTrain requires two user-provided constant values, $\mathrm{N}$ and $\mathrm{E}$, where $\mathrm{N}$ is the approximate number of examples in each region (default = 200,000). $\mathrm{E}$ is the estimated number of examples generated per variant (default = 1.5, estimated using --demo mode based on the number of examples produced from a single chromosome, rounded up to the nearest 0.5). TrioTrain will also internally calculate three additional parameters: $\mathrm{T_g}$, $\mathrm{C}$, and $\mathrm{B}$. $\mathrm{T_g}$ is the total number of variants within the genome-specific truth VCF $\mathrm{C}$ is a set of $\mathrm{n}$ values, where $\mathrm{n}$ is the total number of chromosomes. For $\mathrm{C}$, the values represent the total length (bp) for each chromosome ($\mathrm{c}$) as a proportion of the total genome length (bp). These values are obtained from the Picard reference dictionary file generated automatically by TrioTrain. The series is calculated as $\mathrm{C} = \{\mathrm{c_i} \ldots \mathrm{c_n}\} / \sum_{\mathrm{i=1}}^{\mathrm{n}} \mathrm{C}$. Next, we calculate the number of variants per base pair, as $\mathrm{B} = \frac{\mathrm{T_g}}{\sum_{\mathrm{i=1}}^{\mathrm{n}} \mathrm{C}}$.

We assume that variants are equally distributed across the genome. TrioTrain then creates three series of $\mathrm{n}$ values, where $\mathrm{n}$ is the total number of chromosomes. First, we calculate the number of variants per chromosome as $\mathrm{V} = \{\mathrm{v} \ldots \mathrm{v_n}\} = \mathrm{T_g} * \{\mathrm{c_i} \ldots \mathrm{c_n}\}$. Next, we calculate the length (bp) to sample from each chromosome so that each region BED file will contain the same amount of content per chromosome as the complete genome as $\mathrm{L} = \{\mathrm{l} \ldots \mathrm{l_n}\} = \frac{\{\mathrm{v} \ldots \mathrm{v_n}\}}{\mathrm{B}}$. Lastly, we obtain the number of regions, or B.E.D. files to create as $R_g = \frac{\{c \ldots c_n\}}{\{l \ldots l_n\}} + 1$. We add one additional region for any division remaining across chromosomes. TrioTrain then creates $R_g$ BED files with chromosome number, start, and stop values based on $\{l \ldots l_n\}$ where the start of the subsequent region overlaps the prior region by one base pair. Each BED file is then passed to DeepVariant and executed within $R_g$ SLURM jobs for the make_examples module.

**Supplemental Note 24. Checkpoints used for Warm Starting Training**

Modeler builders should consider how the final model will be used before selecting truth data and a training starting point. For example, we prioritized including the AF channel to enable using our bovine-trained model in legacy, low-coverage samples that are common with animal genomics. However, the warm-starting checkpoint with TrioTrain is user-defined, ensuring species without population-scale resources to avoid including this channel.

The default model checkpoint for DeepVariant v1.4.0, which was used to initialize weights for re-training, was obtained from:
https://console.cloud.google.com/storage/browser/deepvariant/models/DeepVariant/1.4.0/DeepVariant-inception_v3-1.4.0+data-wgs_standard.

Note that the `--make_examples_extra_args` flag is specific to the WGS Allele Frequency (WGS.AF) model and the bovine model checkpoints and, thus, not used with the default WGS model or DeepTrio.

The population allele frequency model for DeepVariant v1.4.0, which was used to compare against our new model, was obtained from:
https://console.cloud.google.com/storage/browser/brain-genomics-public/research/allele_frequency/pretrained_model_WGS/1.4.0;tab=objects?pageState=(%22StorageObjectListTable%22:(%22f%22:%22%255B%255D%22))&prefix=&forceOnObjectsSortingFiltering=false

**Supplemental Note 25. Human Benchmarking Datasets**

The GRCh38 human reference genome was obtained from:
https://ftp.ncbi.nlm.nih.gov/genomes/all/GCA/000/001/405/GCA_000001405.15_GRCh38/seqs_for_alignment_pipelines.ucsc_ids/

The Beam shuffling script was obtained from:
https://raw.githubusercontent.com/google/deepvariant/r1.4/tools/shuffle_tfrecords_beam.py

We used the GIAB benchmarks (4.2.1) for the GRCh38 reference genome, obtained from:
https://ftp-trace.ncbi.nlm.nih.gov/ReferenceSamples/giab/release/

The population VCF for human genomes was obtained from the Google Cloud Storage bucket and will require an active account with Google (i.e., Gmail) to view:
https://console.cloud.google.com/storage/browser/brain-genomics-public/research/cohort/1KGP/cohort_dv_glnexus_opt/v3_missing2ref;tab=objects?pageState=(%22StorageObjectListTable%22:(%22f%22:%22%255B%255D%22))&prefix=&forceOnObjectsSortingFiltering=false

**Supplemental Note 26. Evaluating Performance in Bovine Genomes**

After a training iteration, the selected best checkpoint is immediately used as a custom checkpoint with the one-step variant caller. Although we generate truth labels for these samples, these are not given to the model to call variants but used to assess how the new model changed the resulting callset. DeepVariant is run using:

```
/opt/deepvariant/bin/run_deepvariant \
      --model_type WGS \
      --ref /ref_dir/ARS-UCD1.2_Btau5.0.1Y \
      --reads /bam_dir/<test_genome>.cram \
      --customized_model /start_dir/<new_checkpoint>.ckpt \
      --make_examples_extra_args=
      "use_allele_frequency=true,population_vcfs=/popVCF_dir/<allele_fr
      equencies>.vcf \
      --output_vcf /out_dir/<prefix>.vcf.gz
      --intermediate_results_dir /out_dir/tmp/<prefix> \
      --num_shards (nproc - 1) \
      --exclude_regions Y
```

The resulting compressed VCF is then passed to hap.py (v0.3.12), the tool recommended by the Global Alliance for Genomics and Health (GA4GH) Benchmarking Team (Krusche et al. 2019) and used the vcfeval engine for comparison (Cleary et al. 2015). Although we used software designed for benchmarking VCFs, we could only calculate performance metrics relative to the GATK-derived genotypes we used as truth labels. As such, a True Positive (**TP**) is an identical variant between GATK and DV, a False Positive (**FP**) is a variant missing from our GATK truth labels but detected by DV, and a False Negative (**FN**) is a variant present in our GATK truth labels that DeepVariant missed. We note that hap.py uses an obsolete version of Python (v2.7), so we obtained a working copy within an existing container from Docker (docker://jmcdani20/hap.py:v0.3.12).

For all 30 iterations, we run hap.py in parallel as separate SLURM jobs for each testing genome, with the number of testing samples varying by testing phase. We execute hap.py using:

```
/opt/hap.py/bin/hap.py \
      /truth/<truthVCF> \
      /query/<custom_model_DVoutputVCF> \
      -r /ref/ARS-UCD1.2_Btau5.0.1Y.fa \
      -f /callable/<truthBED> \
      -o /output/<prefix> \
      --write-counts \
      --keep-scratch \
      --scratch-prefix /output/scratch \
      --engine vcfeval \
      --threads $(nproc - 1) \
      --location <autosomes_withX.file>
```

We then used custom Python scripts to process the output VCF from hap.py into CSV files with summary metrics, including precision, recall, and F1 score, and stratified by variant class (SNPs vs INDELs). We calculated precision as $\mathrm{TP} / (\mathrm{TP} + \mathrm{FP})$ and recall as $\mathrm{TP} / (\mathrm{TP} + \mathrm{FN})$. From these, we calculated F1 score as $2 * [(\mathrm{precision} * \mathrm{recall}) / (\mathrm{precision} + \mathrm{recall})]$. After all tests and comparisons with hap.py are complete, the per-sample CSV files with these metrics are concatenated with those from the entire phase-specific testing group. The group's performance metrics are then used to assess model performance within bovine genomes. The F1 score for all tests used the bovine training iterations was plotted using R.

### Supplemental Note 27. Calculating Mendelian Inheritance Errors

First, we created a reference format SDF file required by rtg-tools for the GRCh38 reference and the ARS-UCD1.2_Btau5.0.1Y reference (Rosen et al. 2020). For example, for the human reference, we used:

```
rtg format \
      -o ./triotrain/variant_calling/data/GIAB/reference/rtg_tools/ \
      ./triotrain/variant_calling/data/GIAB/reference/GRCh38_no_alt_ana
      lysis_set.fasta
```

A pedigree file required for rtg-tools mendelian was created for each trio using a custom Python script using trio metadata (UMAGv1 cohort for bovine, NIST for human). We next converted the single-sample VCFs from the various DeepVariant models to BCFs using Bcftools convert (1.14), with an index created using Bcftools index (1.14). We then created a trio VCF constrained to the autosomes and X chromosome only using Bcftools merge (1.14) to combine the per-sample BCFs, with the sample order defined as offspring, father, mother. The trio BCF was also indexed using Bcftools. These files were then passed to rtg-tools using:

```
conda run --no-capture-output ./miniconda_evns/beam_v2.30 \
      rtg mendelian \
      --input <trioVCF> \
      --output <mieVCF> \
      --template <referenceSDF> \
      --pedigree <ped.txt>
```

We note that this results in a VCF that, by default, contains only the PASS variants. Additionally, the variants detected in the offspring with missing genotypes in either parent result in an “uncertain” designation, and thus, excluded from calculations. The reported MIE rate is then extracted from the log file for each trio, with the results plotted in R.

### REFERENCES

Bolger AM, Lohse M, Usadel B. 2014. Trimmomatic: A flexible trimmer for Illumina sequence data. *Bioinformatics* **30**: 2114–2120.

Cleary JG, Braithwaite R, Gaastra K, Hilbush BS, Inglis S, Irvine SA, Jackson A, Littin R, Rathod M, Ware D, et al. 2015. Comparing Variant Call Files for Performance Benchmarking of Next-Generation Sequencing Variant Calling Pipelines. *bioRxiv* **Aug. 3**: 023754. http://biorxiv.org/content/early/2015/08/03/023754.abstract.

Elsik CG, Tellam RL, Worley KC, Gibbs RA, Muzny DM, Weinstock GM, Adelson DL, Eichler EE, Elnitski L, Guigó R, et al. 2009. The genome sequence of taurine cattle: A window to ruminant biology and evolution. *Science* **324**.

Gibbs RA, Taylor JF, Van Tassell CP, Barendse W, Eversole KA, Gill CA, Green RD, Hamernik DL, Kappes SM, Lien S, et al. 2009. Genome-wide survey of SNP variation uncovers the genetic structure of cattle breeds. *Science (80- )* **324**: 528–532.

Guhlin J, Le Lec MF, Wold J, Koot E, Winter D, Biggs PJ, Galla SJ, Urban L, Foster Y, Cox MP, et al. 2023. Species-wide genomics of kākāpō provides tools to accelerate recovery. *Nat Ecol Evol*.

Hayes BJ, Daetwyler HD. 2018. 1000 Bull Genomes Project to Map Simple and Complex Genetic Traits in Cattle: Applications and Outcomes. *Annual Review of Animal Biosciences* **7**: 89–102.
Heaton MP, Smith TPL, Bickhart DM, Vander Ley BL, Kuehn LA, Oppenheimer J, Shafer WR, Schuetze FT, Stroud B, Mcclure JC, et al. 2021. A Reference Genome Assembly of Simmental Cattle, Bos taurus taurus. *Journal of Heredity* **112**: 184–191.
Koren S, Rhie A, Walenz BP, Dilthey AT, Bickhart DM, Kingan SB, Hiendleder S, Williams JL, Smith TPL, Phillippy AM. 2018. De novo assembly of haplotype-resolved genomes with trio binning. *Nature Biotechnology* **36**: 1174–1182.
Krusche P, Trigg L, Boutros PC, Mason CE, De La Vega FM, Moore BL, Gonzalez-Porta M, Eberle MA, Tezak Z, Lababidi S, et al. 2019. Best practices for benchmarking germline small-variant calls in human genomes. *Nat Biotechnol* **37**: 555–560. http://dx.doi.org/10.1038/s41587-019-0054-x.
Low WY, Tearle R, Liu R, Koren S, Rhie A, Bickhart DM, Rosen BD, Kronenberg ZN, Kingan SB, Tseng E, et al. 2020. Haplotype-resolved genomes provide insights into structural variation and gene content in Angus and Brahman cattle. *Nature Communications* **11**.
Oppenheimer J, Rosen BD, Heaton MP, Vander Ley BL, Shafer WR, Schuetze FT, Stroud B, Kuehn LA, McClure JC, Barfield JP, et al. 2021. A Reference Genome Assembly of American Bison, Bison bison bison. *The Journal of heredity* **112**: 174–183.
Rice ES, Koren S, Rhie A, Heaton MP, Kalbfleisch TS, Hardy T, Hackett PH, Bickhart DM, Rosen BD, Ley B Vander, et al. 2020. Continuous chromosome-scale haplotypes assembled from a single interspecies F1 hybrid of yak and cattle. *GigaScience* **9**: 1–9.
Rosen BD, Bickhart DM, Schnabel RD, Koren S, Elsik CG, Tseng E, Rowan TN, Low WY, Zimin A, Couldrey C, et al. 2020b. De novo assembly of the cattle reference genome with single-molecule sequencing. *Gigascience* **9**.
Rowan TN, Hoff JL, Crum TE, Taylor JF, Schnabel RD, Decker JE. 2019. A multi-breed reference panel and additional rare variants maximize imputation accuracy in cattle. *Genetics Selection Evolution* **51**: 1–16.
Stephens ZD, Hudson ME, Mainzer LS, Taschuk M, Weber MR, Iyer RK. 2016. Simulating next-generation sequencing datasets from empirical mutation and sequencing models. *PLoS ONE* **11**: 1–18.
